# Structure-based discovery of highly bioavailable, covalent, broad-spectrum coronavirus-M^Pro^ inhibitors with potent in vivo efficacy

**DOI:** 10.1101/2025.01.16.633443

**Authors:** Tyler C. Detomasi, Gilles Degotte, Sijie Huang, Rahul K. Suryawanshi, Amy Diallo, Luca Lizzadro, Francisco J. Zaptero-Belinchón, Taha Y. Taha, Jiapeng Li, Alicia Richards, Eric R. Hantz, Zain Alam, Mauricio Mantano, Maria McCavitt-Malvido, Rajesh Gumpena, James R. Partridge, Galen J. Correy, Annemarie F. Charvat, Isabella S. Glenn, Julia Rosecrans, Jezrael L. Revalde, Dashiell Anderson, Judd F. Hultquist, Michelle R. Arkin, R. Jeffrey Neitz, Danielle L. Swaney, Nevan J. Krogan, Brian K. Shoichet, Kliment A. Verba, Melanie Ott, Adam R. Renslo, Charles S. Craik

## Abstract

The main protease (M^Pro^) of SARS-CoV-2 is crucial for viral replication and is the target of nirmatrelvir (the active ingredient of Paxlovid) and ensitrelvir. The identification of new agents with differentiated pharmacokinetic and drug resistance profiles will increase therapeutic options for COVID-19 patients and bolster pandemic preparedness generally. Starting with a lead-like dihydrouracil chemotype from a large-library docking campaign, we improved M^Pro^ inhibition >1,000-fold by engaging additional sub-sites in the M^Pro^ active site, most notably by employing a latent propargyl electrophile to engage the catalytic Cys145. Advanced leads from this series, including **AVI-4516** and **AVI-4773** show pan-coronavirus antiviral activity in cells, very low clearance in mice, and for **AVI-4773** a rapid reduction in viral titers more than a million-fold after just three doses, more rapidly and effectively than the approved drugs, nirmatrelvir and ensitrelvir. Both **AVI-4516** and **AVI-477**3 are well distributed in mouse tissues, including brain, where concentrations ten or fifteen-thousand times the EC_90_, respectively, are observed eight hours after an oral dose. As exemplar of the series, **AVI-4516** shows minimal inhibition of major CYP isoforms and human cysteine and serine proteases, likely due to its latent–electrophilic warhead. **AVI-4516** also exhibits synergy in cellular infection models in combination with the RdRp inhibitor molnupiravir, while related analogs strongly inhibit nirmatrelvir-resistant M^Pro^ mutant virus in cells. The *in vivo* and antiviral properties of this new chemotype are differentiated from existing clinical and pre-clinical M^Pro^ inhibitors, and will advance new therapeutic development against emerging SARS-CoV-2 variants and other coronaviruses.

**One sentence summary:** This manuscript describes the discovery of a new class of potent inhibitors of the SARS-CoV-2 major proteases (M^Pro^) with a unique mechanism of inhibition, pan coronaviral activity *in cellulo*, exquisite selectivity vs. the human proteome, and exceptional *in vivo* efficacy in SARS-CoV-2 infection models that surpasses that of currently approved agents.

## Introduction

Four years after the start of the COVID-19 pandemic, the persistent threat of highly transmissible, pathogenic, and immune-evading SARS-CoV-2 variants remains a global concern. SARS-CoV-2 variants are expected to continue emerging and thus, to stop the cycle of infections and emergence of new variants, it is crucial to develop effective direct-acting antiviral therapeutics. Proteolytic processing of the SARS-CoV-2 polyprotein is essential for viral replication and depends on the action of nsp5(*1, 2*) which encodes the main protease (M^Pro^), also referred to as 3CL^Pro^. Targeting the proteases involved in viral replication has a long track record of success in delivering antiviral therapeutics(*3*). Indeed, M^Pro^ is a clinically validated target for SARS-CoV-2, with the M^Pro^ inhibitors nirmatrelvir(*4*) and ensitrelvir(*5*) used clinically to treat COVID-19. SARS-CoV-2 will continue to mutate and generate new resistant variants, which calls for new agents with pan-coronavirus activity and enhanced antiviral spectrum to treat these new infections. Moreover, given the ongoing risk of future pandemics arising from coronaviral reservoirs in bats and in other small mammals(*6, 7*), it is crucial to identify molecules that target evolutionarily conserved domains of M^Pro^ to elicit pan coronaviral antiviral activity. This approach is an essential preparative measure beyond the current pandemic(*8*).

Herein, we describe the discovery of uracil-based, non-peptidic M^Pro^ inhibitors exemplified by the advanced lead molecule **AVI-4516**. Attractive features of this new chemotype include a simple achiral and easily diversified chemical scaffold that exhibits potent biochemical and *in vitro* antiviral activity against multiple SARS-CoV-2 variants as well as other known human coronaviruses. The most efficacious analogs from this series employ an unactivated, latent-electrophilic alkyne to engage the active site cysteine (Cys145) of M^Pro^, leading to potent, irreversible inhibition, both *in vitro* and *in vivo*. In using this very weak electrophile, cross-reactivity with important mammalian proteases is wholly avoided, as are interactions with other important off-targets, such as receptors and ion channels, including the hERG channel. We propose that reactivity with Cys145 is promoted by precise positioning of the latent electrophile adjacent to the oxyanion hole in M^pro^ (*9*), thereby stabilizing the developing negative charge (*9*) in the transition state of the nucleophilic attack. Overall, advanced leads **AVI-4516** and **AVI-4773** manifest many differentiated properties that augur well for the discovery of antiviral development candidates for SARS-CoV2 and related coronaviruses.

## Results

### Docking Campaign Reveals Dihydrouracil Core

As we described previously(*10*), an initial docking screen of 862 million “tangible”, make-on-demand molecules against a deposited [PDB: 6Y2G] M^Pro^ structure returned several scaffolds with mid-low µM inhibition, including the inhibitor **AVI-1084** (Z3535317212, IC_50_: 29 µM). Informed by the docking poses, we sought to improve its potency by exploring the much larger 48 billion molecule space represented by the tangible library, an approach we have used previously(*11–14*). The docked pose of **AVI-1084** suggested favorable hydrogen bond interactions between its dihydrouracil core and the backbone of Glu166 and of Gly143 (**Fig. S1**). With a simple structure, this scaffold was amenable to initial structure activity relationship (SAR) expansion using the SmallWorld search engine(*15*) in ZINC22 (*16*), whereby we identified 17,123 purchasable analogs of **AVI-1084**. Each of these was docked into the M^Pro^ structure to evaluate complementarity with the binding site. From this effort a total of 29 compounds (**Fig. 1A, S1**, **Table S1**) were prioritized for synthesis and tested for activity in an M^Pro^ activity assay. Seven of the analogs showed improved activity, with the most potent, **AVI-3570**, exhibiting an IC_50_ of 1.5 μM (**Fig. 1B**). The improved potency of these analogs stemmed from the introduction of chloro or fluoro substituents augmenting non-polar interactions with the M^Pro^ S2 pocket. We note that the 29 analogs predominantly focused on modifying the thiophene ring of **AVI-1084**, targeting the M^Pro^ S2 pocket, without addressing optimization of the inhibitor’s crucial pyridinone moiety that was modeled to bind the M^Pro^ active site in its S1 pocket. Cognizant that isoquinoline is a privileged structure for the S1 sub-site in M^Pro^ we replaced the pyridinone ring with isoquinoline, leading to **AVI-3778**, **AVI-3779** (**Fig. 1E**), and **AVI-3780**; these analogs had sub-micromolar potencies, about 50-fold improved over the initial docking hit, **AVI-1084**.

**Figure 1.**
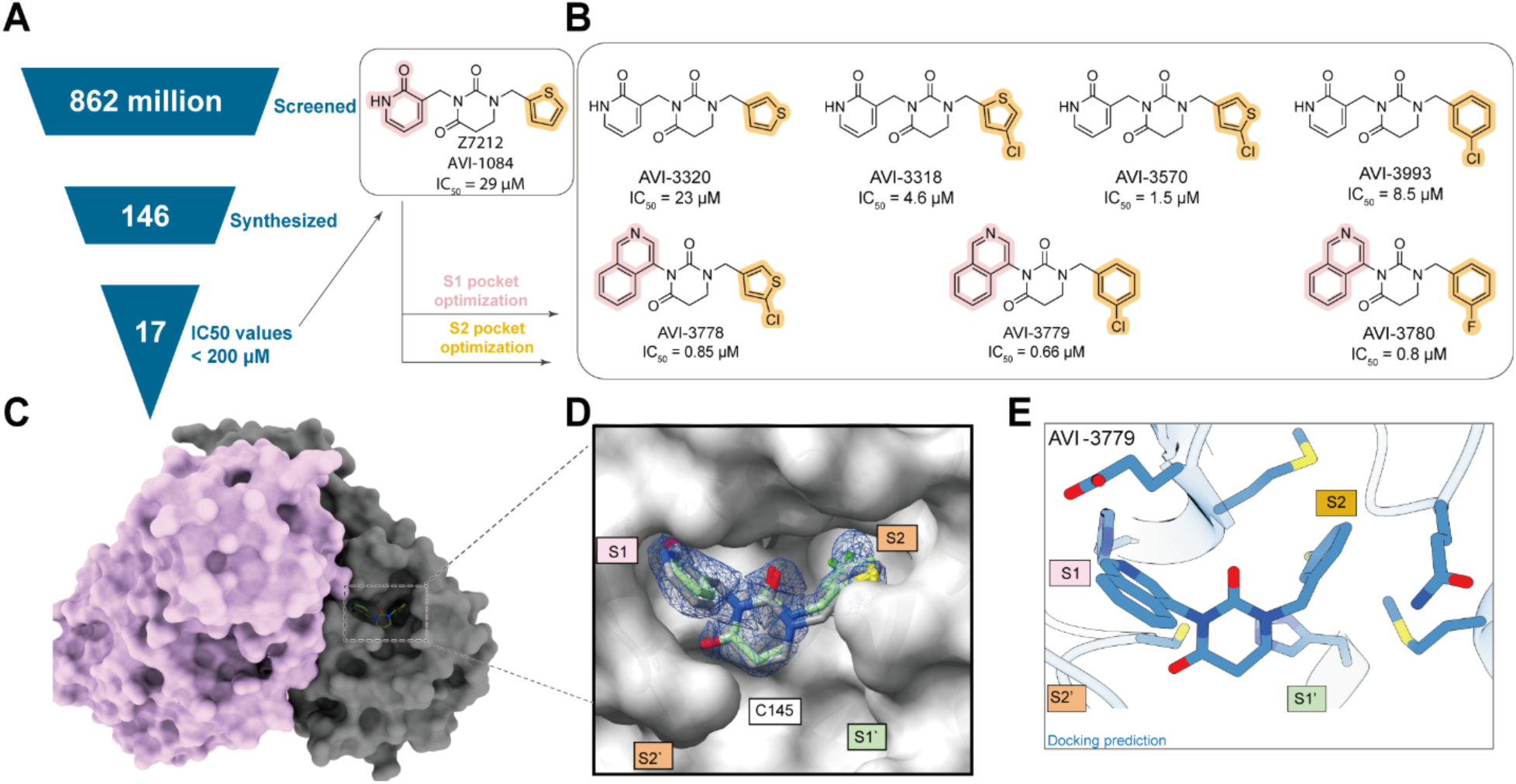
Initial discovery and early structure-based optimization of the AVI-4673 series. A: Large-library docking led to 17 diverse inhibitors, of which Z7212 (AVI-1084) is shown(*10*). **B:** Structure-based optimization of one of them Z7212, explored side chains modeled to bind in the S2 and S1 pockets, leading to sub-µM inhibitors. **C:** Surface representation of the homodimer of M^Pro^. Protomer A is in pink, protomer B is in light brown. **D:** Superposition between the docking predicted (green carbons) and the crystal structure (grey carbons) of the inhibitor AVI-3318 (from panel a). **E:** Docked pose of AVI-3779 the most potent inhibitor based on docking.

To inform further optimization of this scaffold, the x-ray crystal structure of the low µM inhibitor **AVI-3318** in complex with M^Pro^ was determined to 1.96 Å resolution (**Fig. 1C,D, Table S12**). The docking pose of the inhibitor closely superposed with its experimental structure, with an RMSD of 0.45 Å. With only minor discrepancies between the orientation of the chlorine substituent on the thiophene ring, all major interactions predicted by docking were confirmed in the crystal structure – the pyridinone side chain bound within the S1 pocket and in hydrogen-bonding contact with His163, the hydrophobic chlorothiophene group occupying the S2 pocket.

### Optimization of uracil scaffold and discovery of a latent electrophilic moiety affords low-nM inhibitors

Fortified by the correspondence between the docking poses and the structure of **AVI-3318**-M^Pro^ complex, we sought to expand the SAR beyond commercially available compounds, which were limited to substitutions of the dihydrouracil nitrogen atoms. By introducing side chains at the remaining two positions of the dihydrouracil core, we envisaged engaging the S1′ pocket to further improve potency. We also moved to an unsaturated uracil core, as this enabled the synthesis of putatively S1′-targeting analogs without introducing a stereocenter. Docking studies supported the potential of the proposed uracil-derived analogs to target multiple sub-sites in the active site.

To access the desired uracil analogs, we employed a convergent synthesis involving a cyclization reaction between an enaminone and isoquinoline carbamic ester (see SI, Experimental Procedures). The enaminones were either commercially available or synthesized via the Blaise reaction(*17*). This synthesis afforded novel analogs with N3 and C6 substitution, but wherein the N1 position was necessarily unsubstituted. To introduce a C5 substituent we applied a previously described amination procedure in the synthesis of the enaminone intermediate(*18*). Promising early analogs from this effort included **AVI-4301** which was roughly equipotent to **AVI-3318**, and the C5 benzotriazole analog **AVI-4303** (**Fig. 2A,G S4**) that was encouragingly ∼10-fold more potent, with an IC_50_ = 300 nM (**Fig. 2A**). A structure of **AVI-4303** bound to M^Pro^ at 1.58 Å (**Fig. 2D, Table S12**) revealed that the isoquinoline occupied S1 as expected, whereas the benzotriazole unexpectedly bound the S2 pocket and the C6 chlorophenyl side chain stacked in a region between S1′ and S2 positioned near the catalytic dyad residue His41. Compared to the binding of **AVI-3318** then, the uracil core in **AVI-4303** was flipped ∼180 degrees, albeit still anchored by the strong preference for isoquinoline at S1. Further gains in affinity were realized with the introduction of fluoro and choloro substituents on the benzotriazole and aryl rings, respectively, leading to **AVI-4673** with an IC_50_ value of 67 nM (**Fig. 2A, S4**).

**Figure 2.**
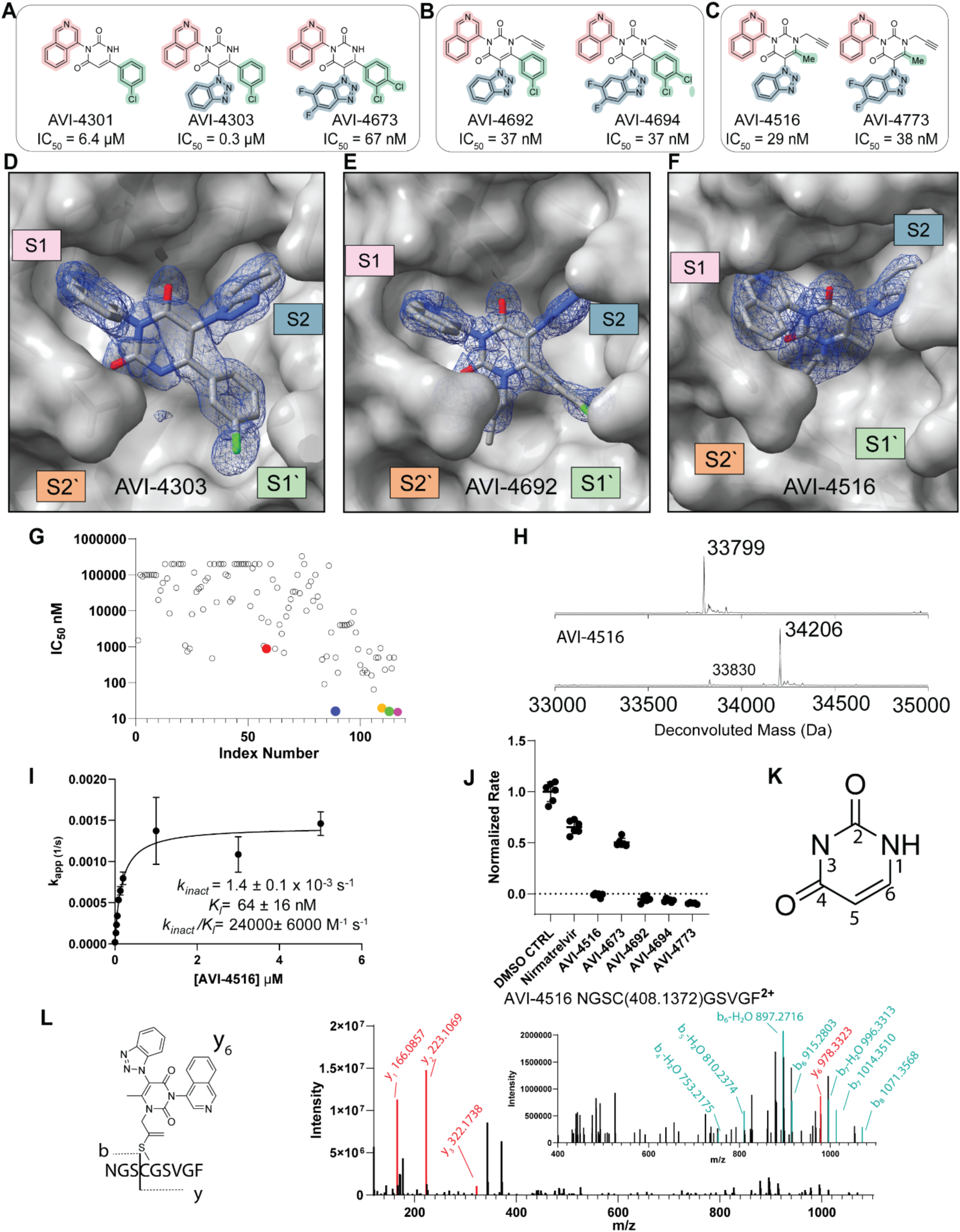
Medicinal chemistry optimization of scaffold and identification of latent electrophilic warhead. The arms of the compounds are highlighted according to the corresponding M^Pro^ subsite. Pink S1, green S1’, blue S2. **A:** Structures of notable compounds during optimization of noncovalent series with uracil core; AVI-4673 is the most potent noncovalent. **B:** Structure of the most potent C6-aryl covalent compounds **C:** The structures of C6-methyl version of covalent alkyne compound, AVI-4516 and AVI-4773. **D-F:** Structures of M^Pro^ bound to inhibitors with subsites anotated;1.58 Å resolution crystal density shown in structure of M^Pro^ with AVI-4303 bound **(D)** with AVI-4692 bound at 1.85 Å resolution **(E)** and AVI-4516 bound at 2.35 Å resolution **(F)**. **G:** Overall comparison of screened compounds with measured dose responses. AVI-4303 is red, AVI-4516 is blue, AVI-4692 is yellow, AVI-4694 is green and AVI-4773 is purple. **H:** Deconvoluted whole protein denaturing mass spectrum of M^Pro^ alone, and M^Pro^ treated with AVI-4516 indicating one modification. **I:** Concentration of AVI-4516 plotted against *k_app_*, and calculated inhibitor kinetic parameters. **J:** Dialysis experiment demonstrating that AVI-4516, AVI-4773, AVI-4692, and AVI-4694 are irreversible covalent inhibitors. **K:** Uracil ring atom numbering nomenclature. **L:** MS2 spectra of chymotrypic peptide of AVI-4516 and the structure of y_6_ ion observed with proposed adduct bound to Cys 145.

With the **AVI-4303** series binding primarily the S1 and S2 pockets (**Fig. 2A,D**), we noted that the unsubstituted N1 position offered a potential vector towards the shallow and comparatively solvent-exposed S1′ pocket, which is also adjacent the catalytic Cys145. To explore binding preferences at S1′, we prepared N1 propargyl analog **AVI-4516**, which we expected would provide access to diverse S1′ targeting analogs via Cu(I)-catalyzed azide–alkyne cycloaddition (CuAAC) reactions of the propargyl function. Surprisingly, while various CuAAC adducts of **AVI-4516** exhibited only low µM potencies, **AVI-4516** itself was very potent, with an IC_50_ of 29 nM (**Fig. 2C, S4**), representing a 100-fold improvement over the des-propargyl comparator **AVI-4375** (IC_50_ 7.4 µM) (**Fig. S2**). A close analog of **AVI-4516** bearing difluoro substitution of the benzotriazole ring (**AVI-4773**) was similarly potent at 38 nM (**Fig. 2C, S4**). We then explored the effect of an N1 propargyl side chain in the context of C6-aryl analogs, finding that both **AVI-4692** and **AVI-4694** possessed potent M^Pro^ inhibition with identical IC_50_ values of 37 nM (**Fig. 2B, S4**).

To confirm that inhibition was due to drug-like binding at the active site, **AVI-4516**, **AVI-4673**, **AVI-4773**, **AVI-4692**, and **AVI-4694** were tested for (artifactual) aggregation induced inhibition. None of the compounds inhibited aggregation-prone inhibition of enzymes like β-lactamase and malate dehydrogenase up to a concentration of 10 µM (**Fig. S3A,C**). Of the five compounds, only **AVI-4694** formed particles by Dynamic Light Scattering (DLS) (**Fig. S3B**), but the critical aggregation concentration (CAC) was 4.3 µM (**Fig. S3D**), a concentration 100-fold higher than required for M^Pro^ inhibition. Thus, dual lines of evidence suggested that C6-methyl (**AVI-4516/4773**) and C6-aryl (**AVI-4673/4694/4692**) analogs act by a drug-like mechanism and not by aggregation at inhibitory concentrations.

### N1 Propargyl Side Chain is a Latent Electrophile

Given the expected proximity of Cys145 to the propargyl group in **AVI-4516**, we hypothesized that nucleophilic attack on the alkyne function might explain the compound’s markedly (∼100-fold) improved potency compared to its des-propargyl comparator. Although rare and generally underappreciated, the reactivity of catalytic cysteines with unactivated alkynes has in fact been demonstrated for deubiquitinases(*9*), cathepsin K(*19*), and even M^Pro^ (*20*). Accordingly, we sought to confirm covalent engagement using several orthogonal methods. First, we evaluated several analogs of **AVI-4516** in which the propargyl side chain was replaced by allyl (**AVI-4690**); homo-propargyl (**AVI-5764**) or butynyl (**AVI-4774**) side chains. All three analogs were devoid of potent M^Pro^ activity with measured IC_50_ > 0.5 mM, a finding consistent with the hypothesis of covalent modification requiring precise positioning of a terminally unsubstituted alkyne (**Fig. S5**). The nitrile congener **AVI-4689** was more active, but surprisingly ∼5-fold less potent than **AVI-4516**. Next, we attempted to detect an **AVI-4516**–M^Pro^ adduct (**Fig. 2H**) by denatured, intact-protein mass-spectrometry (MS) and were pleased to observe a single modification consistent with the mass of **AVI-4516**. Next, we confirmed modification at cysteine (Cys145) by a chymotryptic-digested, fragment MS analysis (**Fig. 2L**). Whole protein MS experiments were also performed with the other propargyl analogs **AVI-4773**, **AVI-4692** and **AVI-4694**; all showed mass shifts consistent with modification at a single site (**Fig. S6A**). Modification of Cys145 by **AVI-4694** was confirmed by analysis of chymotryptic peptides similar to **AVI-4516** (**Fig. S6B**). Consistent with the latent (weakly) electrophilic nature of the propargyl group and the hypothesis of proximity-based reactivity, no covalent adduct was observed in incubations of **AVI-4516** with excess deuterated beta-mercaptoethanol, as observed by ^1^H-NMR (**Fig. S8**).

Next, kinetic parameters of inhibition by **AVI-4516** then measured to test if these compounds were consistent with covalent modification. **AVI-4516** exhibited *k_inact_* and *K_I_* values of 1.4 ± 0.1 x 10^-3^ s^-1^ and 64 ± 16 nM, respectively (**Fig. 2I, S7A**). The modest *k_inact_* is consistent with a weak electrophile, and values reported for other alkyne warheads.(*19*) Overall the inactivation efficiency was reasonable for **AVI-4516** (*k_inact_ /K_I_*= 22,000 ± 6,000 M^-1^ s^-1^), **AVI-4773** (*k_inact_ /K_I_* = 24,400 ± 700 M^-1^ s^-1^), **AVI-4692** (*k_inact_ /K_I_* = 27,000 ± 2,200 M^-1^ s^-1^), and **AVI-4694** (*k_inact_ /K_I_* =48,000 ± 6,000 M^-1^ s^-1^) (**Fig. 2I, S7E-G**), and comparable to *k_inact_ /K_I_* values reported for other covalent protease inhibitors(*21–24*). The *k_inact_/K_I_* values were derived from a linear fit from the *k_app_* vs inhibitor concentration plot; importantly, the raw traces (**Fig. S7A-D**) match what is expected for covalent inhibitors (loss of all activity over time) and the trend in inactivation efficiency matches the potency observed in cells (*vide infra*). To determine if the presumed thioenol ether adduct was subject to hydrolytic instability and regeneration of functional enzyme, a dialysis experiment was performed wherein M^Pro^ was treated with a slight excess of compound (1.5 equiv.) and incubated for 4 hours, after which dialysis was performed for 20 hours (**Fig. 2J**) or for 7 days (**Fig. S7H**). In these experiments M^Pro^ activity was not regained after inhibition by **AVI-4516**, **AVI-4773**, **AVI-4692**, or **AVI-4694**, confirming that the modification is irreversible and the adduct is stable over the timescale examined. In contrast, the thioimidate adduct formed upon incubation with nirmatrelvir was not stable, and partial activity was restored on dialysis, consistent with the expected reversible-covalent reactivity of nitrile warheads (**Fig. 2K, S7H**). These results also suggest that our IC_50_s are over-estimated and are very often close to approximately half the enzyme used, and cell-based readouts provide a more appropriate measure to compare within the series and to other inhibitors.

Lastly, we were able to obtain crystal structures of N1 propargyl analogs **AVI-4516** and **AVI-4692** bound to M^Pro^ (**Fig. 2E,F**). Although **AVI-4516** was solved at 2.35 Å resolution, **AVI-4692** was solved at 1.85 Å allowing for a detailed analysis of the binding mode (**Table S12**). Both compounds retained the same global binding mode of des-propargyl progenitor **AVI-4303**, with their isoquinoline moieties bound in S1 and benzotriazole substituents in S2. The density surrounding Cys145 is most consistent with a thioenol ether adduct formed by reaction at the internal carbon of the alkyne function (**Fig. S16**). This mode of reactivity is also that suggested by structures of M^Pro^ bound to an alkyne analog of nirmatrelvir [PDB:8B2T] and reports of other propargyl-based cysteine warheads(*9, 19*). Both the **AVI-4516** and **AVI-4692** structures exhibited partial occupancy and extra density near Cys145 that is suggestive of partial oxidation of the sulfur, a plausible result of the extended soaks performed to generate these structures.

### M^Pro^ inhibitors exhibit efficacy in SARS-CoV-2 infected cells

Prior to antiviral efficacy assessment in cells, all compounds were evaluated for permeability in a parallel artificial membrane permeability (PAMPA) assay and for cytotoxicity in A549 ACE2 cells (**Table S2**). After demonstrating high passive permeability and a lack of acute cellular toxicity, the most promising compounds were evaluated for antiviral efficacy using a previously described SARS-CoV-2 replicon assay(*25, 26*). In this assay, the SARS-CoV-2 Spike coding sequence is replaced with luciferase and fluorescence reporters to conduct single-round infection and rapid testing of many compounds (**Fig. 3B**). The reporter activity from replicon infected cells has been validated as a surrogate of viral RNA replication(*25*). Several of the lead molecules inhibited viral RNA replication in cells (**Fig. 3A, S9A**); the C6-aryl analogs, **AVI-4692** and **AVI-4694**, exhibited excellent EC_50_ values of 26 nM and 13 nM, respectively.

**Figure 3.**
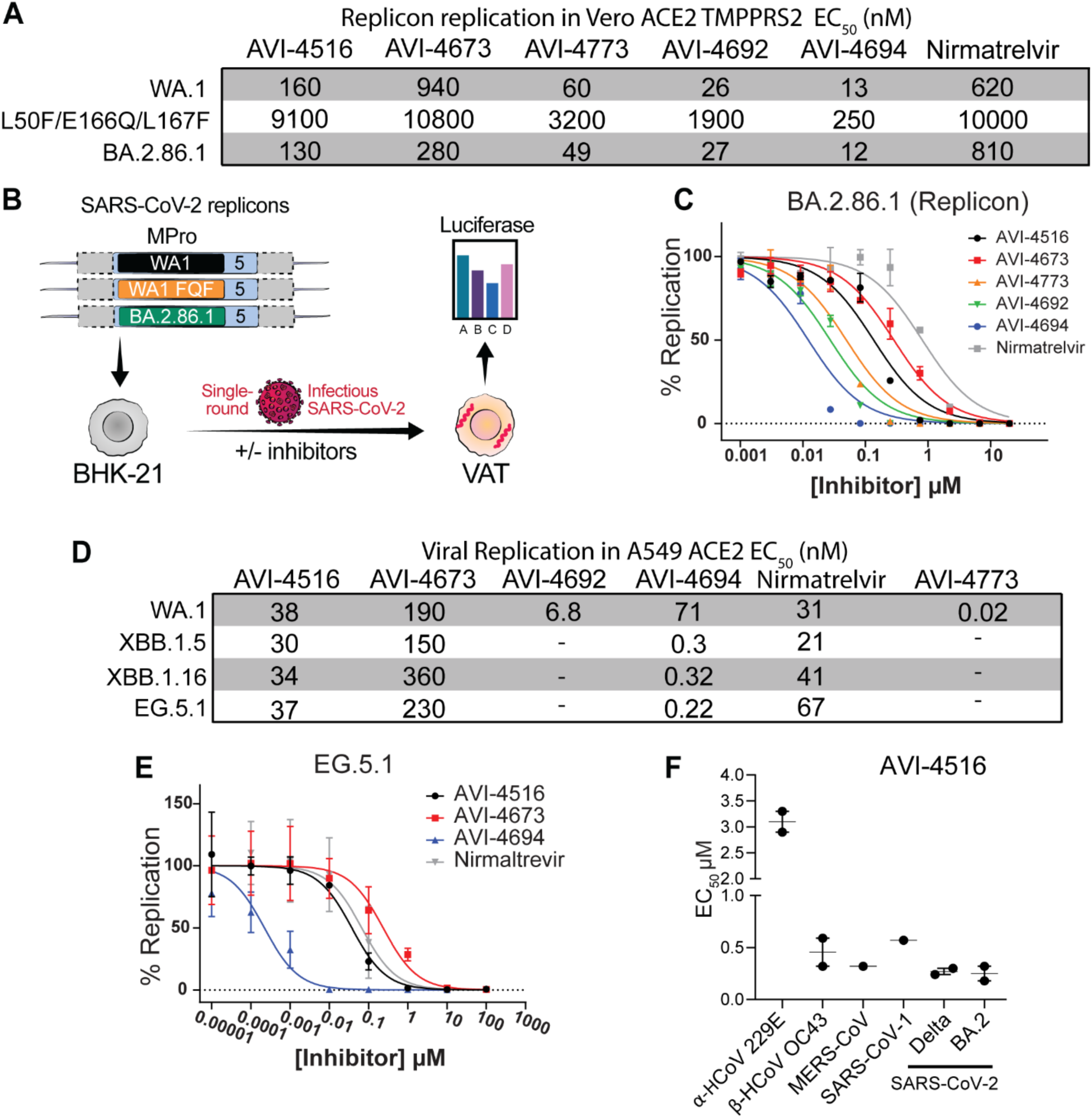
*In cellulo* efficacy of M^Pro^ inhibitors against SARS-CoV-2 Strains, related coronaviruses and Nirmatrelvir resistant mutations. **A:** A table of EC_50_s for dose reponse inhibition of viral replication in replicon based assay. BA.2.86.1 curve is **c** and the rest are located in **Fig. S9A,B**. **B:** a schematic cartoon of the replicon assay for measurement of antiviral potency. **C:** dose response curves for selected compounds in the BA.2.86.1 replicon assay. **D:** A table of EC_50_s for dose reponse inhibition of viral replication in live virus based assay. EG.5.1 curve is **e** and the rest are located in **Fig. S9C-E**. **E:** Dose response curves for selected compounds against EG.5.1 live virus SARS-CoV-2 variants in A549 ACE2 cells **F:** AVI-4516 EC_50_s for pan coronavirus antiviral efficacy screen determined through CPE. All error bars are plotted as ±SD.

To assess the lead compounds’ activity in the context of nirmatrelvir-resistant M^Pro^ (E166Q, A173V, and S144A(*27*)), we evaluated their activity on purified M^Pro^ containing these specific mutations. **AVI-4516** maintained low-nM potency only against the E166Q enzyme with an IC_50_ of 25 nM (**Fig. S10A-D**) while the other tested compounds lost considerable activity (≥10-fold) when compared to their WT M^Pro^ IC_50_ values (**Fig. 3A,B,E**). To determine if this trend held in the replicon system, compounds were then tested against a triple mutant that contains in addition to E166Q two more mutations in M^Pro^ (L50F, L167F). These are the most prevalent resistant mutations at each respective position in natural sequences on the GISAID database (**Fig. 3A, S9B**). Although **AVI-4516,** and **AVI-4773** maintained potency similar to nirmatrelvir in this triple mutant, **AVI-4692** and **AVI-4694** exhibited enhanced potency, likely due to a combination of better permeability and improved inhibitory properties. Notably, all tested compounds exhibited slightly better efficacy against a recent Omicron strain (BA.2.86.1) that was an ancestor of the currently circulating KP.3 variant and contains the relatively fixed P132H mutation(*28, 29*) (**Fig. 3A,C**), suggesting that P132H likely does not affect compound binding.

To determine the inhibitory activity of compounds in authentic replicating SARS-CoV-2, we employed a novel Incucyte-based HTS antiviral screen using multiple SARS-CoV-2 variants. In this assay, we used a genetically encoded fluorescent reporter virus, where the Orf7a and Orf7b coding sequences were replaced with the reporter mNeonGreen (mNG), icSARS-CoV-2-mNG(*30*). This reporter virus was then used to generate recent Omicron variant viruses. The infectivity and replication of the WA1, XBB.1.5, XBB.1.16, and EG.5.1 were evaluated to optimize signal-to-noise ratio. As expected, we found that nirmatrelvir was potent against all viruses tested with an EC_50_ ranging from 21-67 nM consistent with previously reported values(*31*) and has better potency than in the replicon-infected Vero ACE2 TMPRSS2 cells, which have high expression of the xenobiotic transporter, P-glycoprotein (P-gp)(*32*). The non-covalent analog **AVI-4673**, as well as covalent C6-methyl, **AVI-4516,** and C6-aryl **AVI-4694** showed potent antiviral efficacy against the ancestral SARS-CoV-2 WA.1 variant, (**AVI-4516** EC_50_ = 38 nM, **AVI-4673** EC_50_ = 190 nM, **AVI-4694** EC_50_ = 71 nM) while **AVI-4694** was remarkably ≥100-fold more potent than nirmatrelvir against the recent Omicron variants XBB.1.5, XBB.1.16, and EG.5.1, with EC_50_ of 0.30 nM, 0.32 nM, and 0.22 nM, respectively (**Fig. 3B-D, S9D,E**). Taken together, the biochemical, replicon, and live virus assay data suggests that the combination of C6-aryl substitution with the propargyl warhead (as in **AVI-4694**) has great potential to produce an agent that effectively targets recently emergent variants of SARS-CoV2.

Promisingly, the lead molecules high potency against SARS-CoV-1 M^Pro^ (**Fig. S10**) in biochemical assays. To test if this was a general result and if the lead molecules exhibited pan-coronavirus activity, two covalent analogs (one C6-methyl, one C6-aryl) were further tested against live viruses. The viruses tested included human coronaviruses (α-HCoV 229E and β-HCoV OC43), MERS-CoV, SARS-CoV, and various SARS-CoV-2 variants (Delta, BA.2). **AVI-4516** and **AVI-4694** demonstrated pan-coronavirus activity with low EC_50_ values (<3 µM) and high selectivity indices (SI_50_ >170) against all tested variants (**Fig. 3F**, **Table S11**). These findings suggest that **AVI-4516** and **AVI-4694** could serve as effective pan-coronavirus inhibitors. The pan-coronavirus activity is consistent with reactivity directed by the presence of an oxyanion hole positioned near a reactive cysteine and conserved S1 and S2 pockets.

There have been some reports of M^Pro^ inhibitors that exhibit synergy(*33*) when used in combination with RdRp inhibitors. We tested our scaffold to determine if these inhibitors could synergize with an RdRp inhibitor. The inhibitor, molnupiravir, was chosen as it is clinical approved and available orally which could facilitate further studies if successful. Cells that were infected with either WA.1 or XBB.1.16 were then tested with **AVI-4516** and molnupiravir to measure synergy between a C6-methyl compound and the RdRp inhibitor (**Fig. S17A-F**). For the WA.1 strain infected cells, a minor effect was observed in the direction of positive synergy. However, for XBB.1.16 strain of SARS-CoV-2 (**Fig. S17D-F**) synergy was observed at several concentrations when using a ZIP synergy analysis(*34*).

### AVI-4516 has an excellent pharmacokinetic and *in vitro* off-target safety profile

To guide our lead optimization efforts, we performed a panel of standard *in vitro* ADME assays for all new analogs exhibiting potent biochemical activity. Among the leads described herein, **AVI-4516**, **AVI-4773**, **AVI-4692**, and **AVI-4694** all exhibited excellent stability in mouse and human liver microsomes (MLM and HLM T_1/2_ >120 min, **Fig. 4A**). The C6 methyl analogs **AVI-4516** and **AVI-4773** showed plasma protein binding (PPB) that was low to moderate at 61% and 83%, respectively, while C6 aryl analogs **AVI-4692**, and **AVI-4694**, by contrast, had very high PPB at >99% (**Fig. 4A**). Permeability in MDCK-MDR1 monolayers that express P-gp were high in the apical to basolateral direction for **AVI-4516** (13.3 x 10^-6^cm/s), **AVI-4773** (17.8 x 10^-6^cm/s) and **AVI-4694** (8.2 x 10^-6^cm/s) while efflux ratios were reasonable to low at 3.4, 3.99, and 1.7 respectively (**Fig. 4A**). The combined *in vitro* antiviral and ADME data suggested excellent potential for **AVI-4516**, **AVI-4773**, and **AVI-4694** to achieve efficacious plasma and cells/tissues concentrations in animals and thus were nominated for *in vivo* pharmacokinetic profiling.

**Figure 4.**
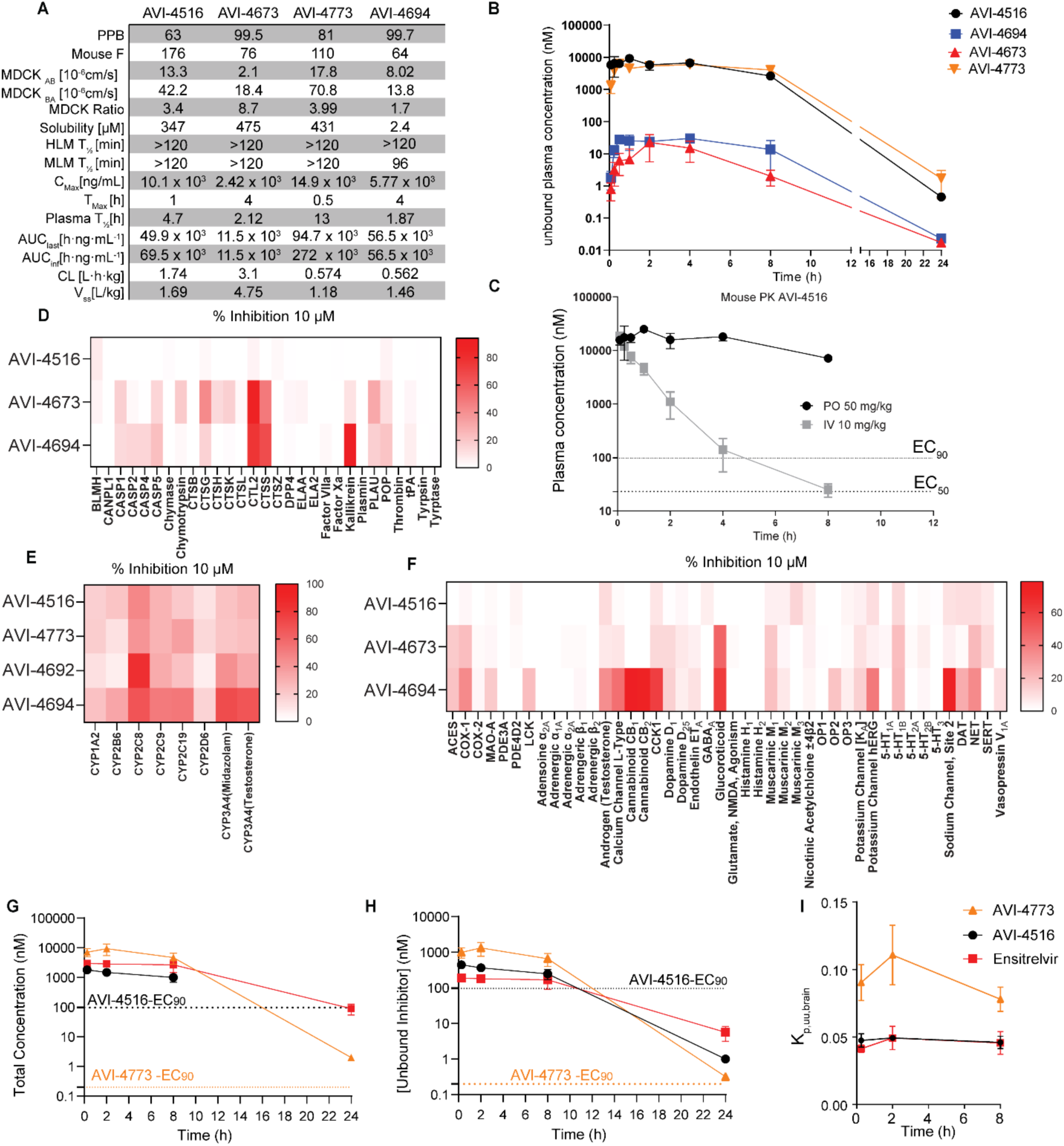
ADME, PK, and safety properties of M^Pro^ inhibitors. A: ADME and PK properties of AVI-4516, AVI-4773, AVI-4692, AVI-4694**. B:** Fraction unbound concentration in mouse plasma of AVI-4516, AVI-4773, AVI-4692 and AVI-4694 after PO 50 mg/kg dosing. **C:** Comparison of mouse plasma concentration of AVI-4516 when dosed 10mg/kg IV or 50 mg/kg PO. The concentration at 24 h was below the LOQ. **D:** Percent inhibition of mammalian peptidase panel when treated with 10 µM of AVI-4516, AVI-4673, AVI-4694. AVI-4516 has low inhibition across the panel. Proteases that inhibited >50% have dose response curves that generated IC_50_s in **Fig. S13**. **E:** Percent inhibition of human CYP panel of AVI-4516, AVI-4773, AVI-4692 and AVI-4694 compared to Nirmatrelvir and Ensitrelvir. **F:** Percent inhibition of mammalian receptor panel when treated with 10 µM of AVI-4516, AVI-4673, AVI-4694. AVI-4516 has low inhibition across the panel. Proteins that were inhibited >50% have dose response curves that generated IC50s in **Fig S13 A,B**. **G:** Total brain and plasma concentration of AVI-4516, AVI-4773 and ensitrelvir after PO 100mg/kg dose. all error bars are plotted as ± SD. **H**: Unbound concentration of AVI-4516, AVI,4773, and ensitrelvir in mouse brain. **I**: unbound brain to plasma partioning coefficient for AVI-4516, AVI-4773, and ensitrelvir.

We chose doses of 50 mg/kg PO and 10 mg/kg IV pharmacokinetic (PK) experiments with **AVI-4516**, **AVI-4773**, **AVI-4694,** and **AVI-4673**, which were performed in male CD-1 mice (**Fig. 4B S11A-D, Table S4-7**). All four compounds showed low clearance, and remarkably so for **AVI-4694** and **AVI-4773** with *in vivo* clearance just ∼7% of hepatic blood flow in the mouse. Total exposure by AUC was highest for C6 methyl analogs **AVI-4773** and **AVI-4516**, and lowest for **AVI-4673**, consistent with the considerably lower permeability of this analog in the MDCK-MDR1 assay. Overall, the plasma exposure profiles, and oral bioavailability of the lead compounds were excellent, with the oral bioavailability (%F) values of **AVI-4516** and **AVI-4773** exceeding 100%. These very high apparent F values may reflect a slow intestinal absorption process(*35*) or might be due to saturation of clearance mechanisms at the rather high oral dose of 50 mg/kg (as compared to 10 mg/kg in the IV arm). Using measured PPB values to correct for plasma protein binding returned a free concentration of 2,615 nM for **AVI-4516** and 4,045 nM for **AVI-4773** at the 8-hour timepoint (**Fig. 4B**), values approximately 26-fold higher than the cellular EC_90_ = 97 nM for **AVI-4516** in the WA.1 strain (**Fig. 3A**). Accordingly, we predicted that a 50 mg/kg or higher dose of either **AVI-4516** or **AVI-4773** should retain efficacious antiviral concentrations at 12 hours and that twice-daily (BID) dosing would be effective in mouse efficacy studies.

Paxlovid, ensitrelvir, as well as other recently reported(*4, 5, 36–38*) pre-clinical agents are known to be inhibitors of CYP3A4, with the potential for drug-drug interactions that must be monitored and can lead to adverse events in some patient populations. Accordingly, we evaluated **AVI-4516**, **AVI-4773**, **AVI-4692**, and **AVI-4694**, for inhibition of important human CYP isoforms at a fixed concentration of 10 µM. Both **AVI-4516** and **AVI-4773** exhibited minimal inhibition (≤ 25% at 10 µM) across the panel, with the exception of CYP2C8 (∼45% at 10 µM). The C6-aryl analog **AVI-4692** by contrast was a somewhat more potent inhibitor of both CYP2C8 and CYP3A4 while **AVI-4694** showed the overall poorest profile, inhibiting several CYP isoforms >50% at 10 µM (**Fig. 4E**). We next evaluated three exemplar analogs: **AVI-4673** (non-covalent analog), **AVI-4516** (C6 methyl analog) and **AVI-4694** (C6 aryl analog) for off-target activity across a panel of 40 receptors, ion channels (including the hERG channel), and serine and cysteine proteases (**Fig. 4D,F**). Of the three leads, latent-electrophilic analog **AVI-4516** bearing C6 methyl substitution showed the most exceptional *in vitro* safety profile, demonstrating no significant inhibition or interference with any of the off-targets at 10 µM. Of the other leads, noncovalent analog **AVI-4673** was found to be a low micromolar inhibitor of cathepsin L2 (**Fig. S12A**), while C6 aryl analog **AVI-4694** showed low micromolar inhibition of nine enzymes/receptors in the panel (**Fig. S12B**). To evaluate **AVI-4516** against a complex proteome we turned to a thermal proteome profiling (TPP) assay(*39*). To identify concentrations where effective binding could be observed in a TPP assay, we first used increasing concentrations of **AVI-4516** with M^Pro^ protein alone and observed a significant increase in T_m_ due to binding at all concentrations tested (**Fig. S12A**). From this experiment we selected 50 µM **AVI-4516** for TPP analysis of cellular lysate from A549 cells. Here, we observed no proteins with a statistically significant T_m_ increase that would be consistent with binding, and 4 proteins with a moderate decrease in T_m_ that could be the result of either direct binding or secondary impacts. In summary, these studies revealed the N-propargyl, C6 methyl chemotype exemplified by **AVI-4516** (and also present in **AVI-4773**) as devoid of significant off-target binding with human proteins (**Fig. S12B**).

Given the superior off-target selectivity profile of the N-propargyl/C6-methyl chemotype, we sought to explore the distribution of **AVI-4516** across mouse tissues following a 100 mg/kg oral dose. Of particular interest was exposure in lung, bronchial alveolar fluid (BALF), and brain, given that infection is centered in the respiratory tract, while reservoirs of virus may persist in brain(*40*). Encouragingly, we found that **AVI-4516** and **AVI-4773** were more significantly distributed into pharmacologically relevant compartments like lung and BALF than was ensitrelvir. Thus, **AVI-4516** maintains very high exposure compared to its cellular EC_90_ (WA.1 live virus **Fig 3B, S9C**) at 8 hours in mouse heart (227x EC_90_), lungs (47x EC_90_) and BALF (11.8x EC_90_) (**Fig. S11E,F Table S8**) while **AVI-4773** exhibited even higher exposures in these tissues, which was especially notable given its even lower cellular EC_90_ values. (**Fig. S11G,H Table S10**). For both analogs, exposure in brain was considerably lower than in plasma or other tissues, but total brain concentration of **AVI-4516** at 8 hours (**Fig. 4G**) was still ∼10-fold greater than its antiviral EC_90_ vs. the WA.1 strain. On account of its ∼4.7-fold higher exposure in brain and picomolar EC_90_ value, the total brain concentration of **AVI-4773** at 8 hrs was some 15,000-fold greater than its cellular EC_90_ vs. the WA.1 strain. Using measured binding to mouse plasma protein and brain homogenate, we calculate an unbound brain-to-plasma partitioning coefficient (K_P,uu,brain_) of ∼5%, and ∼8% for **AVI-4516** and **AVI-4773**, respectively at 8 hours. Overall, the biodistribution studies demonstrate that **AVI-4773** is more favorably partitioned into the lung, heart, and BALF when compared to ensitrelvir, whilst exhibiting ∼10-fold higher free drug concentrations in brain 8 hours after a single oral dose (**Fig. 4H**). While directly analogous data for nirmatrelvir is unavailable, a study of nirmatrelvir in the rat (when dosed at an allometrically scaled dose of 60 mg/kg nirmatrelvir and 20 mg/kg ritonavir/day) revealed brain concentrations only 3 times the respective EC_90_ value(*41*). In summary, the favorable PK profile, significant free fraction, and favorable biodistribution profile of **AVI-4516** and **AVI-4773** nominated these compounds as promising lead compounds for *in vivo* studies of antiviral efficacy.

### AVI-4516 and AVI-4773 demonstrate potent *in vivo* antiviral efficacy

We next turned to a mouse infection model to elucidate the *in vivo* antiviral effects of irreversible M^pro^ inhibition by **AVI-4516** and **AVI-4773**. An initial study in C57BL/6 mice infected with the SARS-CoV-2 Beta compared nirmatrelvir (300 mg/kg BID) and **AVI-4516** (100 mg/kg BID) with vehicle-treated animals. Of note, the Beta variant contains a natural mutation (N501Y) in its Spike protein that allows non-lethal infection of wild-type mice, while M^Pro^ of the Beta variant contains the K90R mutation(*42*). As shown in the schematic (**Fig. 5A**), treatment began at 4 hours post-infection with oral BID dosing continuing for 5 days post-infection, during which we closely monitored body weight as a marker for severity of infection (**Fig. S15A**). At 2-, 4-, and 7-days post-infection, a subset of mice (n=5) from each group was euthanized to determine the virus titers through plaque assays (**Fig. 5B**).

**Figure 5.**
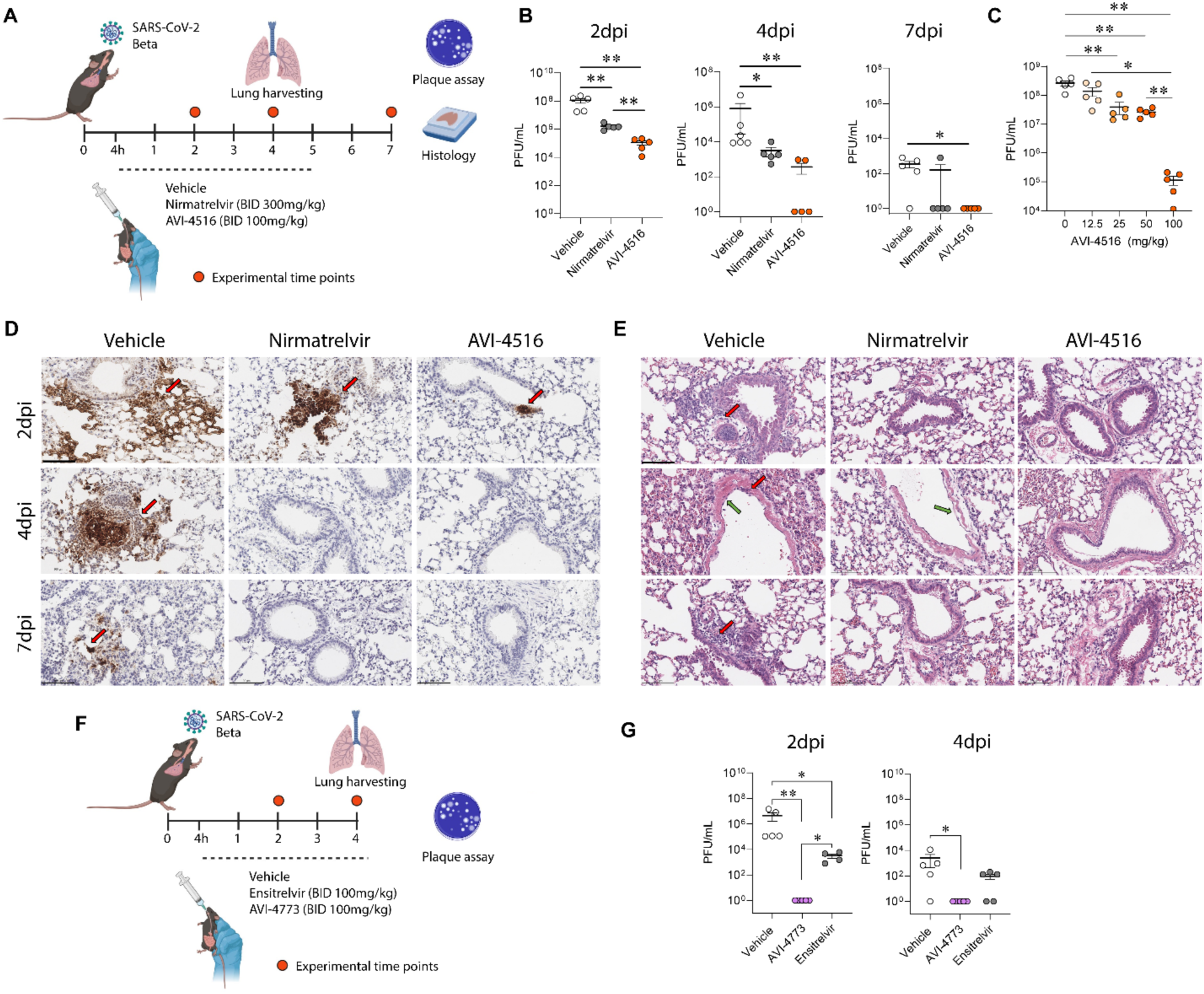
Oral administration of AVI-4516 limits virus replication. A: Schematic of antiviral efficacy experiment. Wild type mice were intranasally infected with 10^3^ plaque forming units of the SARS-CoV-2 Beta variant. Infected animals were orally dosed (BID) with either vehicle, nirmatrelvir or AVI-4516. The lung tissues were harvested and processed for further analysis at 2-, 4-, and 7-days post-infection (dpi) (n=5 per group per time point). **B:** Graphs presenting mature virus particles from the lungs of infected mice at the indicated time points are presented. The data are shown as the mean ± SEM. for each time point and were analyzed using a two-tailed unpaired Student’s t-test. Each dot represents the infectious virus titer in an individual mouse. **C:** A separate group of mice were infected with the SARS-CoV-2 Beta variant and treated with various concentrations of AVI-4516. Virus titers were measured at 2 dpi. **D:** Representative images of immunohistochemistry for the SARS-CoV-2 N protein in the left lung lobe of mice from different treatment groups at the specified time points. Scale bars represent 100µm. **E:** Hematoxylin and eosin (H&E) staining of lung tissue. Significant mononuclear cell infiltrations were marked by red arrows, and severe injury in respiratory epithelia, characterized by epithelial cell debris in the lumen and incomplete epithelial regeneration, was highlighted by green arrows Scale bars represent 100µm.

As expected, for this model, we observed no weight loss in any treatment group until day 7 (**Fig. S14A**). Encouragingly, measurement of mature virus particles using plaque assays revealed the potent antiviral efficacy of **AVI-4516**, with a 3- to 4-log reduction in virus replication compared to the vehicle, while the positive control, nirmatrelvir, showed a 2-log reduction (**Fig. 5B**). Histology data indicated heightened staining for the SARS-CoV-2 N protein in the vehicle treatment group through immunohistochemistry. In contrast, the nirmatrelvir and **AVI-4516** treatment groups exhibited minimal or no viral protein present at all time points (**Fig. 5D, S13A**). Peribronchiolar and perivascular infiltrations by mononuclear cells and respiratory epithelial cell injury were observed in the vehicle-treated group, to a minimal extent in the nirmatrelvir-treated group, and were notably absent in the **AVI-4516**-treated mouse lungs, underscoring the rapid action and superior efficacy of **AVI-4516** compared to a higher dose of nirmatrelvir in this model (**Fig. 5E, S13B**).

In an independent experiment the dose-dependent antiviral efficacy of **AVI-4516** was determined using SARS-CoV-2 Beta-infected WT mice, where the mice were treated orally with **AVI-4516** at doses ranging from 12.5 to 100 mg/kg. The animals were euthanized at day 2 post-infection to determine the virus titers in the lung tissues. The results of this study demonstrated a dose-dependent decrease in virus replication and allowed the calculation of an IC_50_ value of 14.7 mg/kg (**Fig. 5C**). From this data it is evident that three doses of **AVI-4516** at 25 mg/kg is sufficient to significantly reduce viral load at 2 days post-infection. In summary, **AVI-4516** exhibits significant dose-dependent antiviral efficacy, demonstrating a significant reduction in mature virus particle production compared to the vehicle- and nirmatrelvir-treated mice. Additionally, the histology data show that early treatment with **AVI-4516** mitigates signs of lung inflammation.

Encouraged by the superior efficacy of **AVI-4516** as compared to nirmatrelvir, we next compared the efficacy of **AVI-4516** and difluoro congener **AVI-4773** to ensitrelvir, which is regarded as more efficacious than single-agent nirmatrelvir in mouse models(*43*) and in our models performed better at a third the dose. Thus, a series of experiments were performed comparing **AVI-4516** or **AVI-4773** to ensitrelvir using a 100 mg/kg BID dosing regimen. In the experiments with **AVI-4516** and ensitrelvir, we observed similar reductions in viral titers for the two test articles, as compared vehicle-treated controls 2dpi (**Fig. S15B**). More strikingly, we found that **AVI-4773** conferred a dramatic and rapid reduction of viral titers to below detectable levels by day two, after just three doses (**Fig. 5F**). This represented a >3-log reduction in viral load at day 2 when compared to the ensitrelvir-treated arm and a ∼6-log reduction compared to vehicle-treated animals at day 2 (**Fig. 5G**). In a second, identical study, we confirmed this powerful pharmacodynamic effect, with a reduction of viral titers to below detectable levels after just three doses of **AVI-4773** (**Fig. S15C**). The remarkable pharmacodynamics of **AVI-4773** can be understood in light of the compound’s high exposure in BALF of ∼4,400 nM at 8 hours after a single oral dose (**Fig. S11H**) and potent, low- or sub-nM antiviral effects in cellular models.

## Discussion

The approval of nirmatrelvir and ensitrelvir for clinical use was followed rapidly by efforts from various academic and industrial groups to identify improved, next-generation inhibitors of SARS-CoV-2 M^Pro^. Many of these next-generation compounds are inspired by, or based on nirmatrelvir, differing in the side chains and the nature of the cysteine-targeting warhead (*36–38, 44, 45*). A second class of inhibitors are entirely non-peptidic in nature and based on cyclic uracil, dihydrouracil, or hydantoin cores (*5, 33, 46–49*) from which aromatic or aliphatic arms are displayed. These latter compounds, like ensitrelvir, act by reversible, non-covalent mechanisms of inhibition. Here we describe a distinct chemotype (**Fig. S3F**) that combines the general trifold architecture of ensitrelvir with a latent-electrophilic warhead, resulting in a unique mechanism of M^Pro^ inhibition and differentiated antiviral and pharmacodynamic properties. These improvements are exemplified by the exquisite off-target safety profile and potent pan-coronaviral *in cellulo* activity of **AVI-4516**, and the superior pharmacokinetics, and tissue distribution of **AVI-4773**, which together with potent antiviral activity, produce a remarkable pharmacodynamic effect in infected mice.

The rapid reduction of viral titers conferred by **AVI-4516,** and especially **AVI-4773**, suggest the potential for more convenient, less frequent dosing regimens and might extend the window for effective treatment of infection. Further preclinical assessment, will be required to predict a human dose and to determine optimal dosing regiments of killing effects of **AVI-4773** in animals stands out among other recently disclosed M^Pro^ inhibitors. In part, these properties derive from the unique mode of inhibition conferred by an unactivated N-propargyl side chain. We posit that successful capture of Cys145 with this very weak electrophile requires placement of the alkyne function near the oxyanion hole, which promotes nucleophilic attack of Cys145 by stabilizing the developing negative charge at the terminal carbon and its eventual protonation, plausibly by His 41 of the catalytic dyad. Thus, the irreversible inhibition of M^Pro^ by **AVI-4516**, **AVI-4773** and related analogs distinguish this chemotype mechanistically from nirmatrelvir (reversible-covalent) and ensitrelvir (noncovalent), while offering apparent advantages in terms of target engagement and pharmacodynamic effect, which we continue to explore. Additional features of this new chemotype include a simple achiral structure and straightforward synthesis in four or five steps, which implies good potential for low manufacturing costs, an important criterion in the context of global coronavirus pandemic preparedness and the stockpiling of drug substance.

In nirmatrelvir-resistant mutants, **AVI-4516** performs at least as well as nirmatrelvir while C6-aryl congeners (*e.g.* **AVI-4694**) exhibit even more potent antiviral activity, likely due to a combination of improved permeability and the formation of additional active-site contacts beyond those of nirmatrelvir. Importantly the C6-chlorophenyl substituent of **AVI-4692** contacts His41 of the catalytic dyad, which is absolutely conserved across all coronavirus M^Pro^ enzymes and may, in part, explain the enhanced spectrum of this chemotype. The C6-aryl subcategory is thus a promising one for further expansion of antiviral spectrum to combat current and future variants. Synergy has been explored previously as a treatment modality for SARS-CoV-2 that has the potential to circumvent mutational pressure(*33*). Promisingly, **AVI-4516** has synergy with an orally available RdRp inhibitor in a cellular infection model with the omicron strain of SARS-CoV-2, XBB.1.16.

Coronaviruses can infect the brain leading to inflammatory syndrome and diverse neurological symptoms, and possibly even contributing to poorly understood conditions like “long COVID”. As demonstrated here, **AVI-4773** crosses the blood brain barrier in mice, with an unbound brain concentration ∼8% of that in plasma and at least 1000-fold higher that the antiviral EC90 at 8 hours. This suggests that **AVI-4773** could serve as an in vivo test article to better understand coronavirus infection and the brain. Above all, the discovery of compounds such as **AVI-4516** and **AVI-4773** reveals an advanced pre-clinical lead series with differentiated properties and excellent prospects to deliver a pan-coronavirus therapeutic development candidate. This discovery approach and unique mechanism of inhibition of these compounds also provide a roadmap for the discovery of antiviral scaffolds that target cysteine proteases of other viruses of concern.

## Materials and Methods

Materials and methods are reported in the supplementary information.

## Acknowledgements

Advanced light Source beamline 8.3.1 and SLAC National Accelerator Laboratory for the access to the synchrotron and Dr. Violla Bassim for the help for the data collection. NIAID AVIDD grant number U19AI171110. Assessment of ADMET targets (Cardiac Channel Profiling, CYP Induction, Peptidase Selectivity Panel, and Secondary Pharmacology Profiling) was performed via NIAID’s NIH preclinical services for *in vitro* assessment (Contract No. 75N93019D00021/75N93023F00001/HHSN272201800007I*) In vivo* antiviral screening (pan-coronavirus assays) was performed via NIAID’s preclinical services (SRF No. 2021-1229-003). We thank Prof. James Fraser for his insight and help with the structural biology. We thank Dr. Norbert Bischofberger, Dr. Zach Sweeney, and Dr. Spiros Liras for their guidance and input throughout this project. We would also like to thank Dr. Saumya Gopalkrishnan for critical reading of the manuscript and help with coordination of experiments. MO received support from the Roddenberry Foundation, from P. and E. Taft, and the Gladstone Institutes.

## Supplementary information for

### Supplementary Figures and Tables

**Supplementary Fig. 1.**
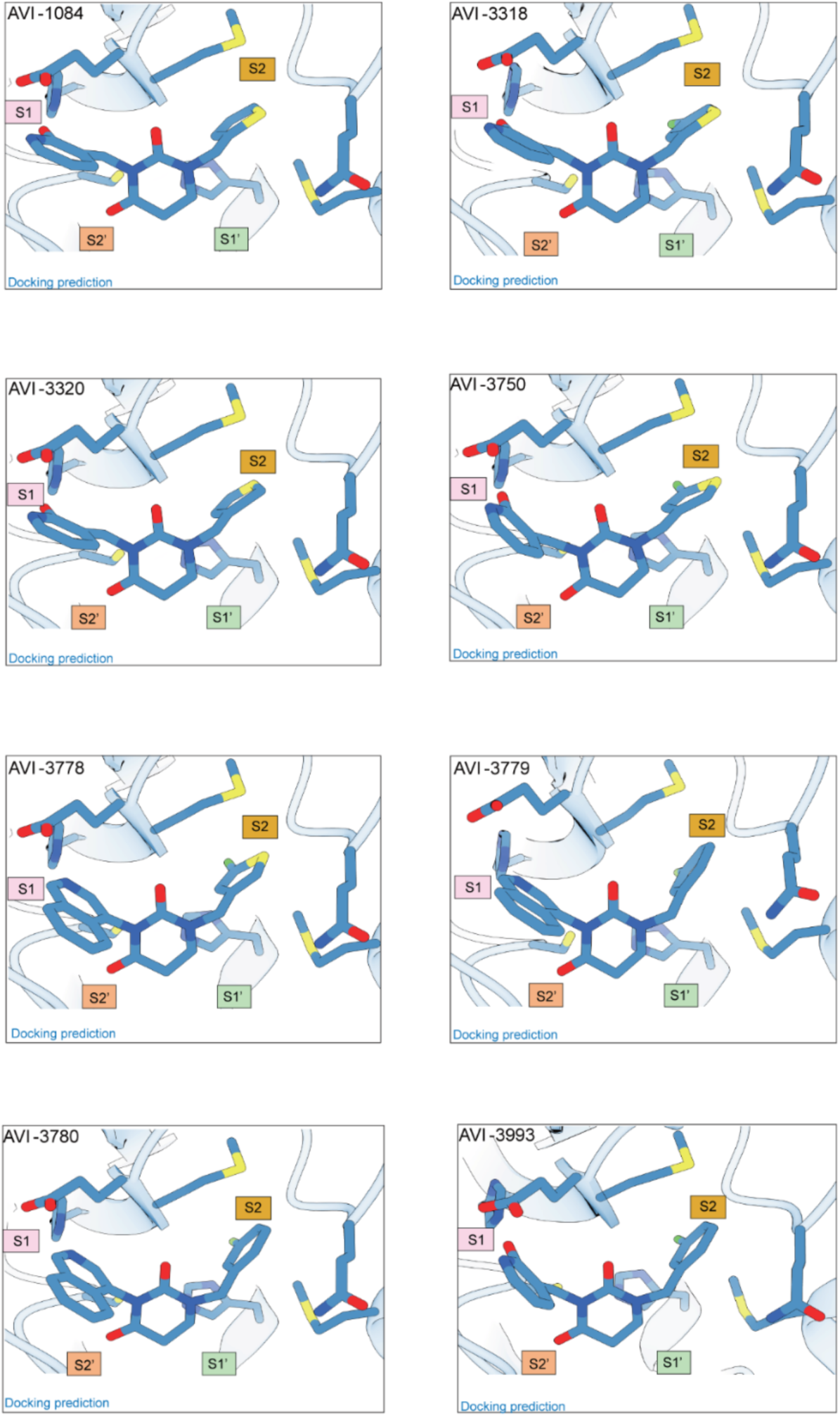
Docking poses for compounds in Fig. 1. The nearby M^Pro^ S subsites are annotated.

**Supplementary Fig. 2.**
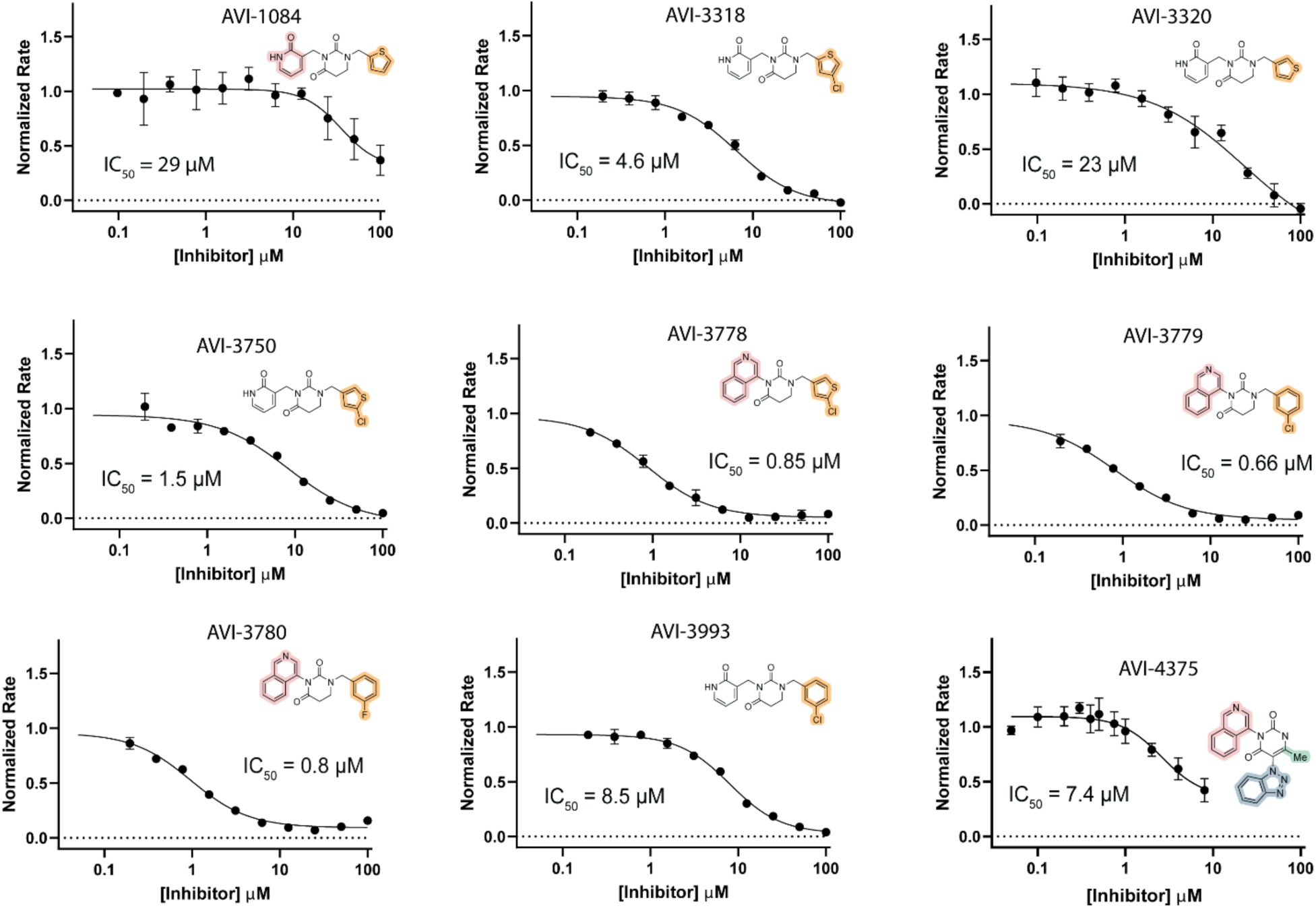
Dose response curves of selected compounds. (AVI-1084, AVI-3318, AVI-3320, AVI-3750, AVI-3778, AVI-3779, AVI-3780, AVI-3993, AVI-4375). Each point was performed in technical triplicate. All compounds were fit using four parameter inhibitor vs response equation in GraphPad Prism to obtain an IC_50_ and error bars are plotted as ± S.D. All rates were normalized to DMSO control.

**Supplementary Table 1.**
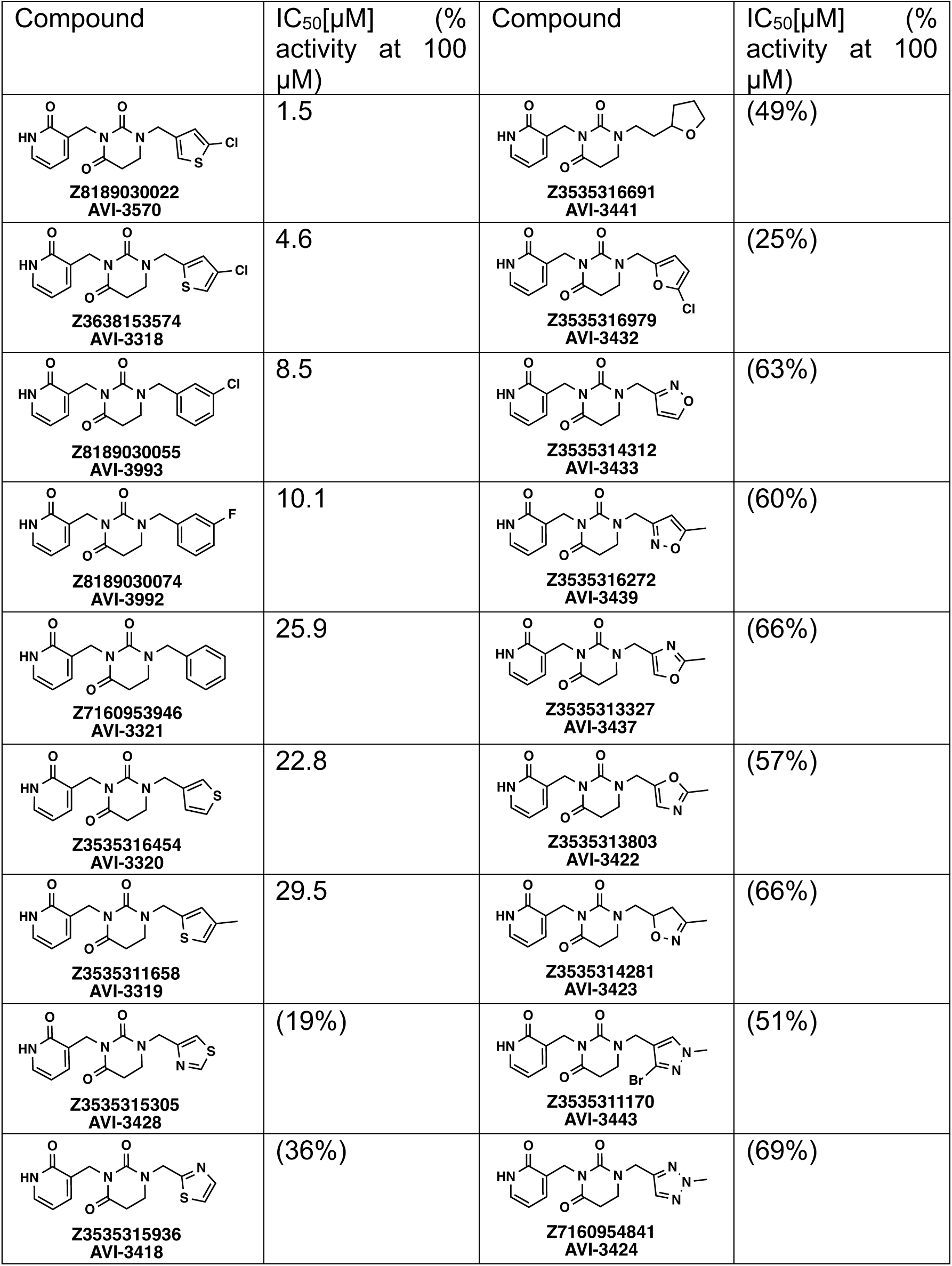
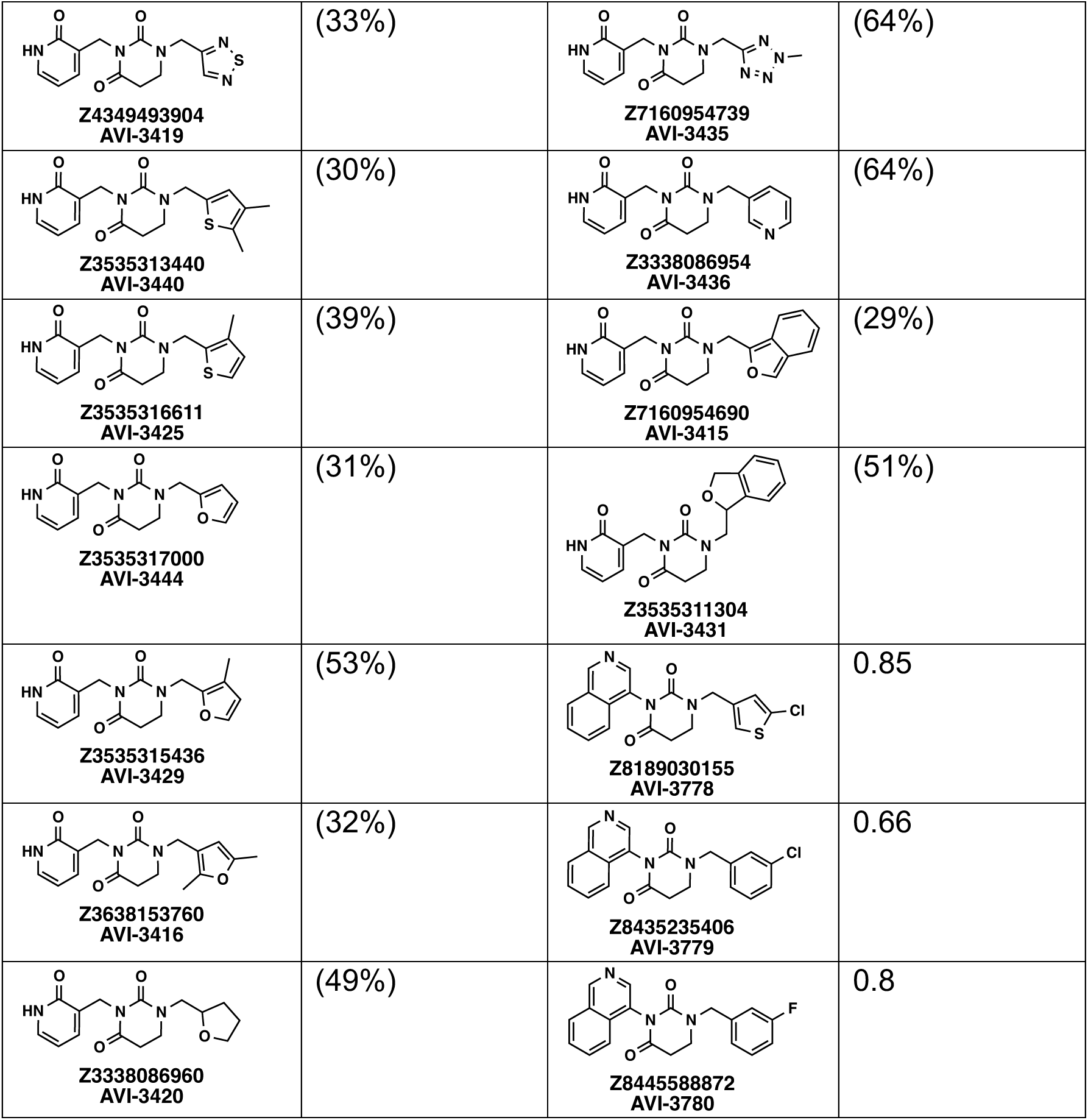
Chemical structure and *in vitro* activity of compounds identified by docking. Table contains IC_50_s or % M^Pro^ activity when treated with 100 µM of compound.

| Compound | IC<sub>50</sub>[μM] (% activity at 100 μM) | Compound | IC<sub>50</sub>[μM] (% activity at 100 μM) |
| --- | --- | --- | --- |
| **Z8189030022**<br>**AVI-3570** | 1.5 | **Z3535316691**<br>**AVI-3441** | (49%) |
| **Z3638153574**<br>**AVI-3318** | 4.6 | **Z3535316979**<br>**AVI-3432** | (25%) |
| **Z8189030055**<br>**AVI-3993** | 8.5 | **Z3535314312**<br>**AVI-3433** | (63%) |
| **Z8189030074**<br>**AVI-3992** | 10.1 | **Z3535316272**<br>**AVI-3439** | (60%) |
| **Z7160953946**<br>**AVI-3321** | 25.9 | **Z3535313327**<br>**AVI-3437** | (66%) |
| **Z3535316454**<br>**AVI-3320** | 22.8 | **Z3535313803**<br>**AVI-3422** | (57%) |
| **Z3535311658**<br>**AVI-3319** | 29.5 | **Z3535314281**<br>**AVI-3423** | (66%) |
| **Z3535315305**<br>**AVI-3428** | (19%) | **Z3535311170**<br>**AVI-3443** | (51%) |
| **Z3535315936**<br>**AVI-3418** | (36%) | **Z7160954841**<br>**AVI-3424** | (69%) |
| **Z4349493904**<br>**AVI-3419** | (33%) | **Z7160954739**<br>**AVI-3435** | (64%) |
| **Z3535313440**<br>**AVI-3440** | (30%) | **Z3338086954**<br>**AVI-3436** | (64%) |
| **Z3535316611**<br>**AVI-3425** | (39%) | **Z7160954690**<br>**AVI-3415** | (29%) |
| **Z3535317000**<br>**AVI-3444** | (31%) | **Z3535311304**<br>**AVI-3431** | (51%) |
| **Z3535315436**<br>**AVI-3429** | (53%) | **Z8189030155**<br>**AVI-3778** | 0.85 |
| **Z3638153760**<br>**AVI-3416** | (32%) | **Z8435235406**<br>**AVI-3779** | 0.66 |
| **Z3338086960**<br>**AVI-3420** | (49%) | **Z8445588872**<br>**AVI-3780** | 0.8 |

**Supplementary Fig. 3.**
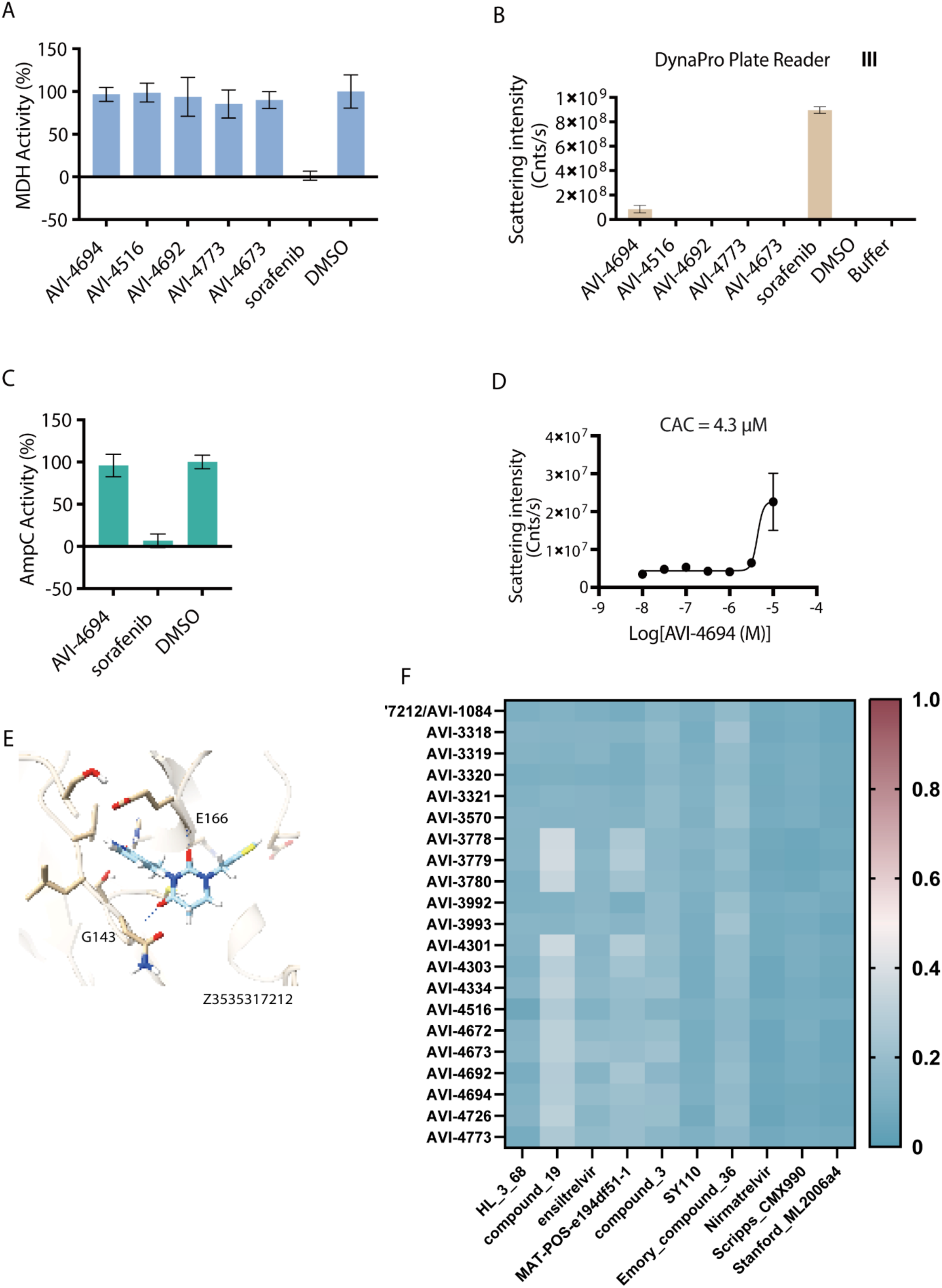
Aggregation testing of inhibitors. **A:** MDH activity when treated with compounds listed to test for non-specific aggregation-based inhibition. Sorafenib is used as a positive control. **B:** DLS measurement for detection of aggregation at 10 μM. **C:** AMPc activity in the presence of AVI-4694 and positive control to test for nonspecific aggregation. **D:** Dose response scattering measurement of AVI-4694 to determine critical aggregation concentration (CAC) **E:** Docking pose of Z3535317212 (AVI-1084) **F:** Comparison of Tanimoto coefficients for compounds reported here vs potent M^Pro^ small molecule inhibitors compounds: HL-3-68(*1*), compound_19(*2*), ensitrelvir(*3*), MAT-POS-e194df51-1(*4*), compound_3(*5*), SY110(*6*), Emory_compound_36(*7*), nirmatrelvir(*8*), Scripps_CMX990(*9*), Stanford_ML2026a4(*10*).

**Supplementary Fig. 4.**
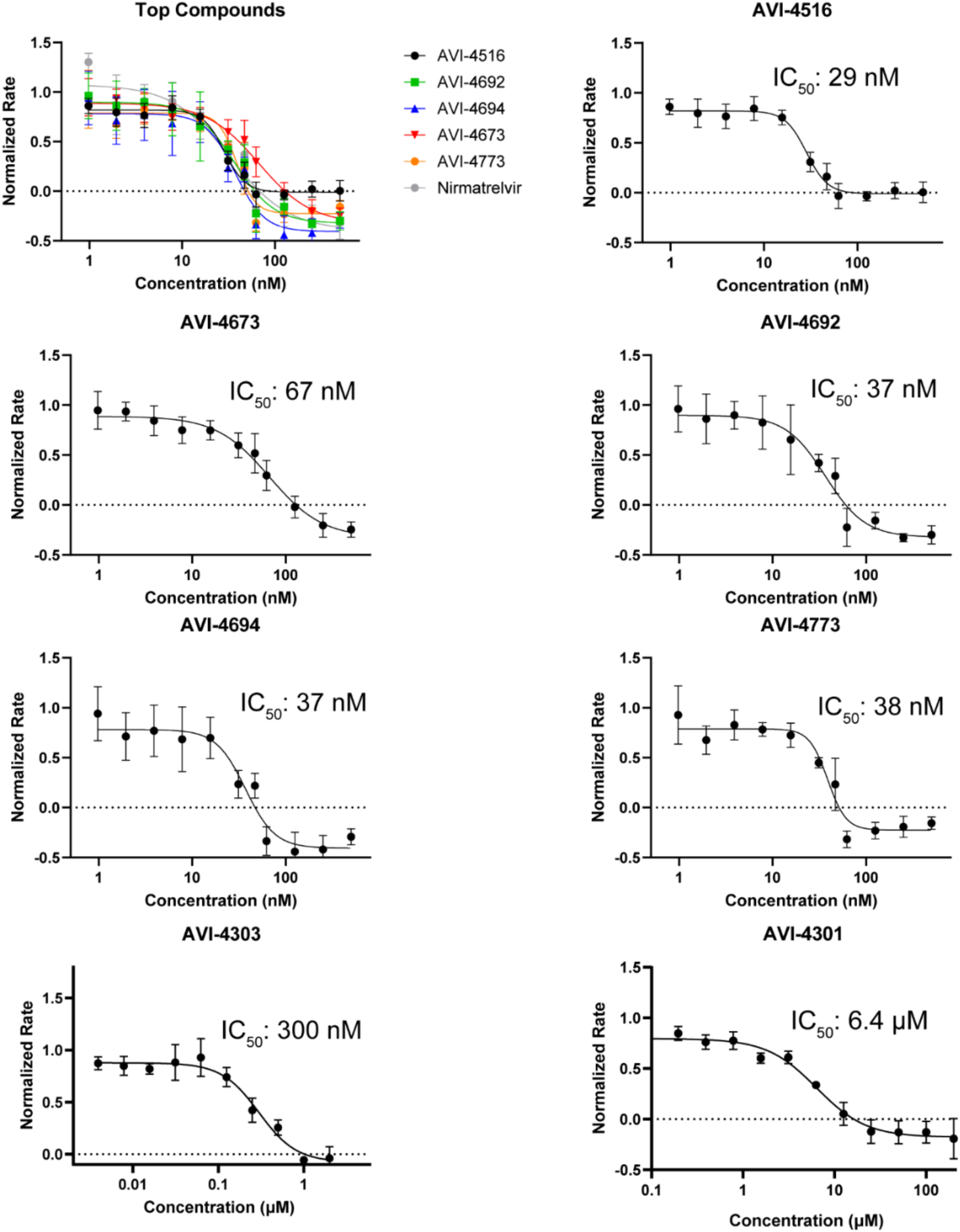
Biochemical assay dose response of compounds from Figure 2. Compounds were incubated with M^Pro^ for 1 h and activity was then measured to generate the curve. All rates were normalized to DMSO control. Each assay was performed in technical triplicate and plotted ± S.D. and fit to a four parameter IC_50_ equation using Prism. Of note, while we present these IC_50_s, for any covalent inhibitor, this number is time dependent and is used only as a comparison with the noncovalent compounds in the rest of the series. Additionally, lower enzyme concentrations often do not exhibit reliable signal which may overestimate the IC_50_s in some cases.

**Supplementary Fig. 5.**
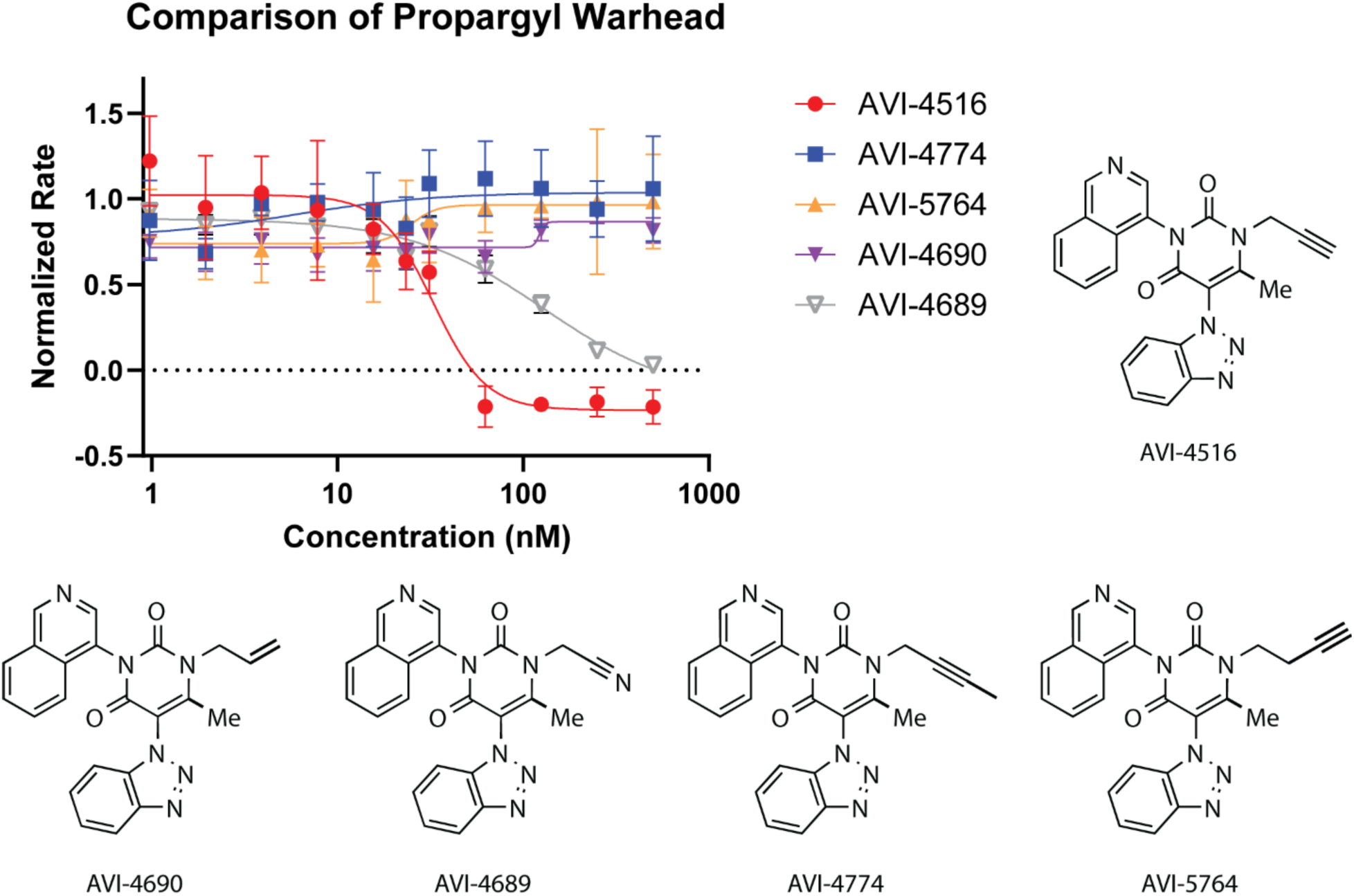
Biochemical *in vitro* dose response and structures of warheads for 4516 analogs. Only the compound (AVI-4689) with the nitrile version retained activity in this dose range. All rates were normalized to DMSO control. Each assay was performed in technical triplicate and plotted ± S.D. and fit to a four parameter IC_50_ equation using Prism.

**Supplementary Fig. 6.**
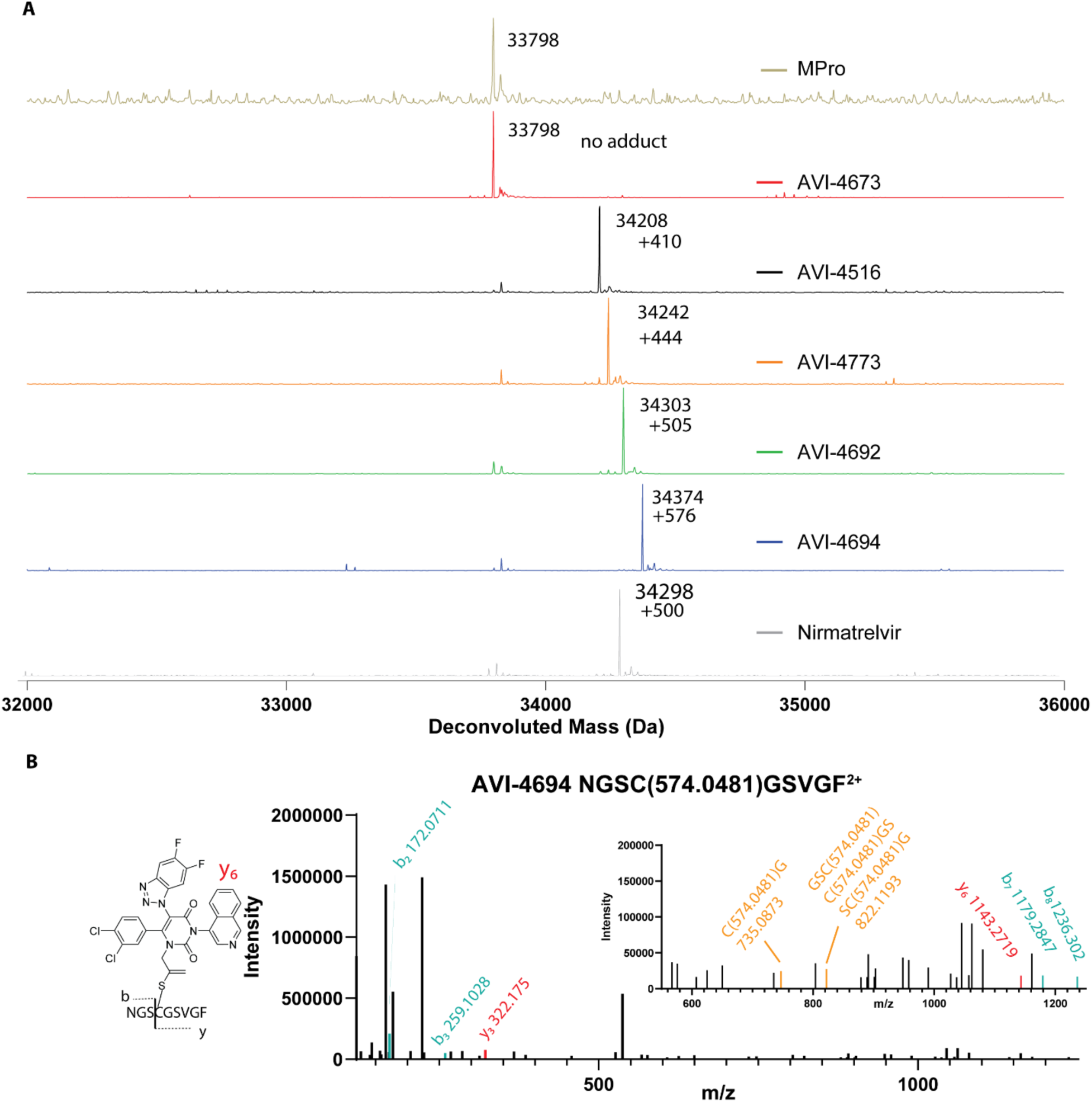
Mass spectrometry investigation of covalency for propargyl warhead. **A:** Deconvoluted whole protein denaturing MS experiment. Comparison of top inhibitors in this study: AVI-4516, AVI-4673, AVI-4773, AVI-4692, AVI-4694. 10 µM of M^Pro^ was treated with 100 µM of compound overnight, then diluted to 500 nM and analyzed via MS. The observed adduct after deconvolution is noted next to main peak. All cases where an adduct was consistent with one modification. **B:** Structure of predicted adduct based on previous literature(*11*) and the structure of AVI-4692/AVI-4516 bound to M^Pro^ MS2 spectra of M^Pro^ treated with 4694 and then digested with chymotrypsin. The MS1 ion that was selected was NGSC(574.0481)GSVGF^2+^ Y_6_ and b_3_ ions are noted in the spectra. The ion that comprises C(574.0481)G shows that the modification is localized to the cysteine.

**Supplementary Fig. 7.**
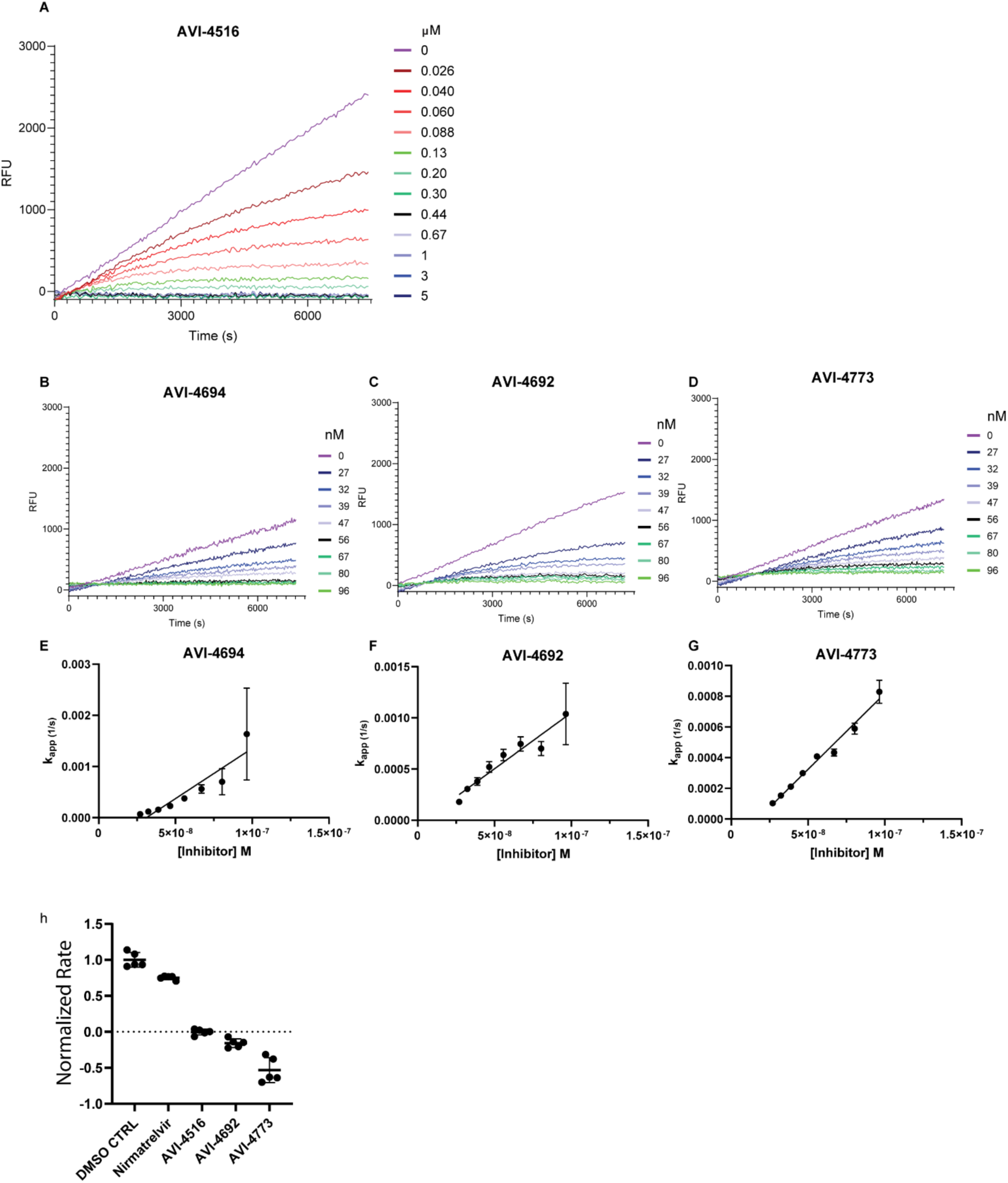
Inhibitor-kinetic experiments for AVI-4516, AVI-4773, AVI-4692, and AVI-4694. **A:** Average blank subtracted raw traces for AVI-4516. **B-D:** Average blank subtracted raw traces for AVI-4694, AVI-4692, and AVI-4773. **E-G:** A plot of *k_app_* vs inhibitor concentration and linear fit used to determine *k_inact_/K_I_*. All inhibition kinetic experiments were done in technical quintuplicate. Error bars are plotted as 95% CI. **H.** Dialysis experiment after 7 days of incubation with compound. Normalized rates when 100 µL of 1 µM M^Pro^ is treated with 1.5 µM inhibitor and then dialyzed against 300 mL of assay buffer for 7 d at RT then diluted to 50 nM enzyme in kinetic assay (final 60,000 x dilution). Each rate was then measured in technical quintuplicate error bars are ± S.D.

**Supplementary Fig. 8.**
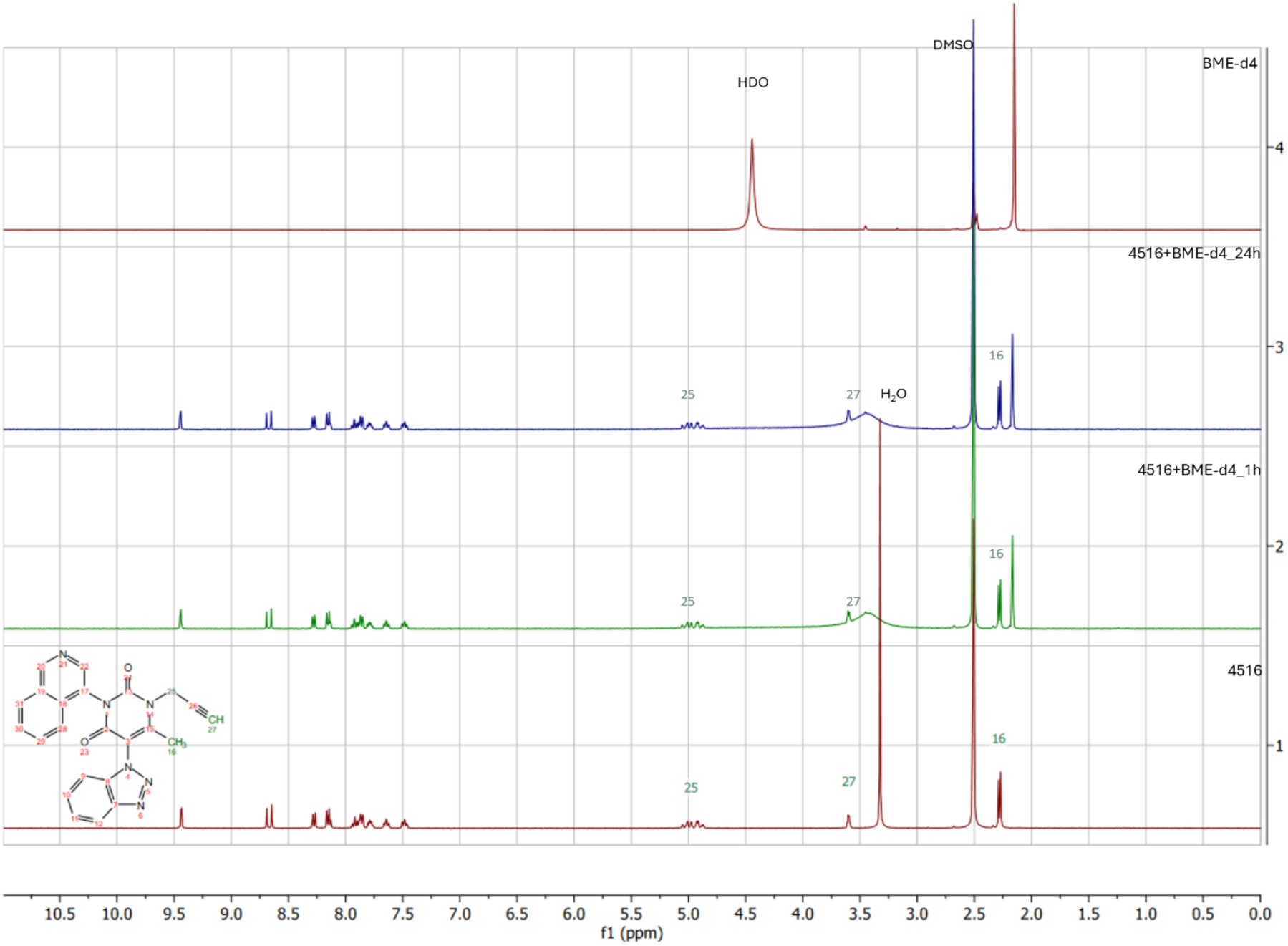
AVI-4516 is unreactive with a large excess (10 equiv.) of the ^1^H NMR-silent thiol *d4*-betamercaptoethanol (*d4*-BME). ^1^H NMR spectrum of AVI-4516 alone (bottom red trace), and in the presence 10 equiv. of *d4*-BME after 1 h (green trace) and 24 h (blue trace) vs. *d4*-BME alone (top red trace). These spectra indicate the lack of reactivity with *d4*-BME as a thiol nucleophile as the AVI-4516 spectrum is unchanged whereas obvious changes to the alkyne resonance (denoted as peak 27) and appearance of new alkene resonances for the thioenol ether product was not observed.

**Supplementary Table 2:** CC_50_ in A549 cells and percent cell viability at 100 µM.

| **Molecule Name** | **A549 CC<sub>50</sub> [µM]** | **A549 Viability at 100 µM (%)** | **Molecule Name** | **A549 CC<sub>50</sub> [µM]** | **A549 Viability at 100 µM (%)** |
| --- | --- | --- | --- | --- | --- |
| **AVI-3778** | >100 | 95.7 | **AVI-3443** | >100 | 100 |
| **AVI-3779** | >100 | 100 | **AVI-3444** | >100 | 96.9 |
| **AVI-3780** | >100 | 100 | **AVI-3570** | >100 | 100 |
| **AVI-4301** | >100 | 100 | **AVI-4434** | >100 | 100 |
| **AVI-4303** | >100 | 92.3 | **AVI-4435** | >100 | 94.7 |
| **AVI-4434** | >100 | 96.9 | **AVI-4436** | >100 | 99.7 |
| **AVI-4516** | >100 | 90.4 | **AVI-3419** | >100 | 94.4 |
| **AVI-4673** | >100 | 94.4 | **AVI-3420** | >100 | 100 |
| **AVI-1027** | >100 | 100 | **AVI-3421** | >100 | 100 |
| **AVI-1084** | >100 | 100 | **AVI-3422** | >100 | 100 |
| **AVI-1242** | >100 | 100 | **AVI-3423** | >100 | 96.8 |
| **AVI-3318** | >100 | 88.9 | **AVI-3425** | >100 | 94.2 |
| **AVI-3319** | >100 | 100 | **AVI-3426** | >100 | 100 |
| **AVI-3320** | >100 | 100 | **AVI-3428** | >100 | 100 |
| **AVI-3321** | >100 | 100 | **AVI-3429** | >100 | 100 |
| **AVI-3415** | >100 | 100 | **AVI-3430** | >100 | 100 |
| **AVI-3416** | >100 | 100 | **AVI-3431** | >100 | 100 |
| **AVI-3417** | >100 | 100 | **AVI-3432** | >100 | 78.1 |
| **AVI-3418** | >100 | 99.6 | **AVI-3433** | >100 | 98.2 |
| **AVI-3437** | >100 | 100 | **AVI-3434** | >100 | 100 |
| **AVI-3438** | >100 | 70.3 | **AVI-3435** | >100 | 100 |
| **AVI-3439** | >100 | 97.1 | **AVI-3436** | >100 | 100 |
| **AVI-3440** | >100 | 99.1 | **AVI-3441** | >100 | 100 |

**Supplementary Table 3:** Permeability of selected compounds.

| **Molecule Name** | **PAMPA-Gut Papp**<br>**(10<sup>-6</sup>cm/s)** |
| --- | --- |
| **AVI-4143** | 42.3 |
| **AVI-4692** | 31.8 |
| **AVI-3778** | 26.4 |
| **AVI-3779** | 23.9 |
| **AVI-3780** | 19.6 |
| **AVI-4301** | 31.8 |
| **AVI-4303** | 6.6 |
| **AVI-4434** | 12.9 |
| **AVI-4516** | 22.4 |
| **AVI-4673** | 4.99 |

**Supplementary Fig. 9.**
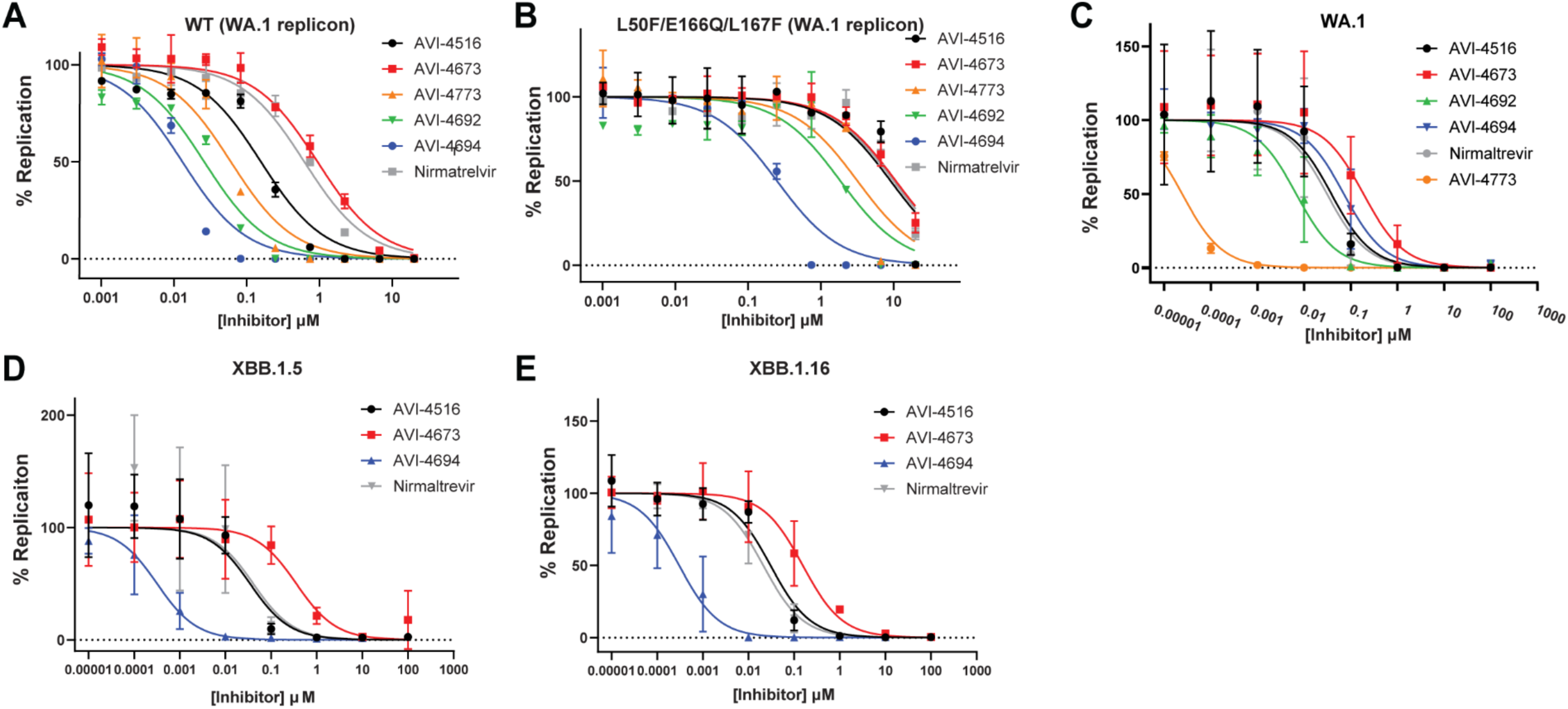
Dose response curves of SARS-CoV-2 replicon and viral infection. **A-B:** Replicon-based dose response curves. **C-E:** Live virus Incucyte-based measurements. Each point was measured in biological triplicate. Error bars were plot ± S.D. Inhibition curves were fit using Prism.

**Supplementary Fig. 10.**
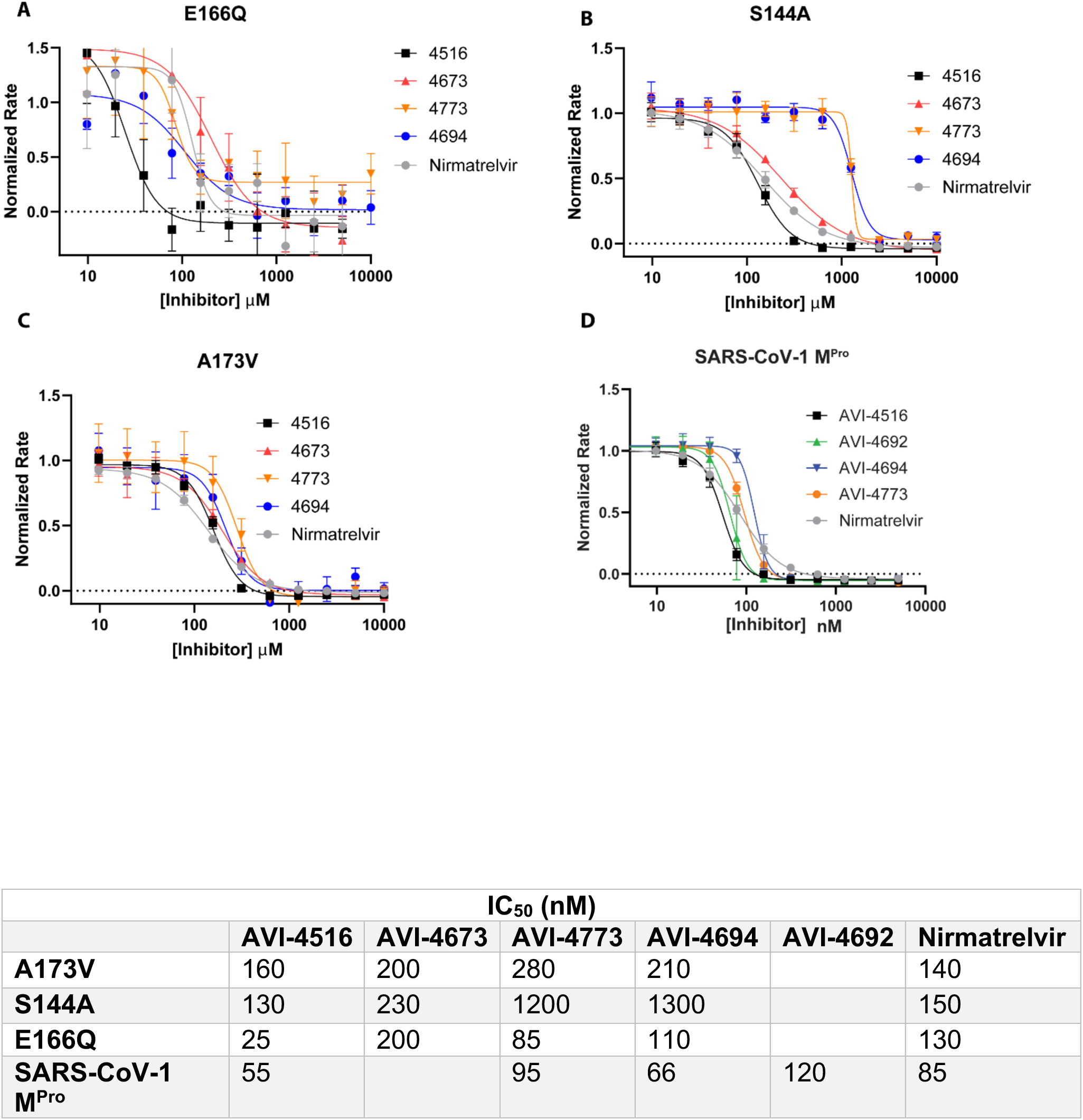
Dose response of AVI-4516, AVI-4673, AVI-4773, AVI-4694, and N=nirmatrelvir vs activity of selected nirmatrelvir resistant mutants and SARS-CoV-1 M^Pro^. Each point was performed in technical triplicate. All compounds were fit using four parameter inhibitor vs response equation in Prism to obtain an IC_50_. All rates were normalized to DMSO control. Error bars were plotted as ± S.D.

**Supplementary Fig. 11.**
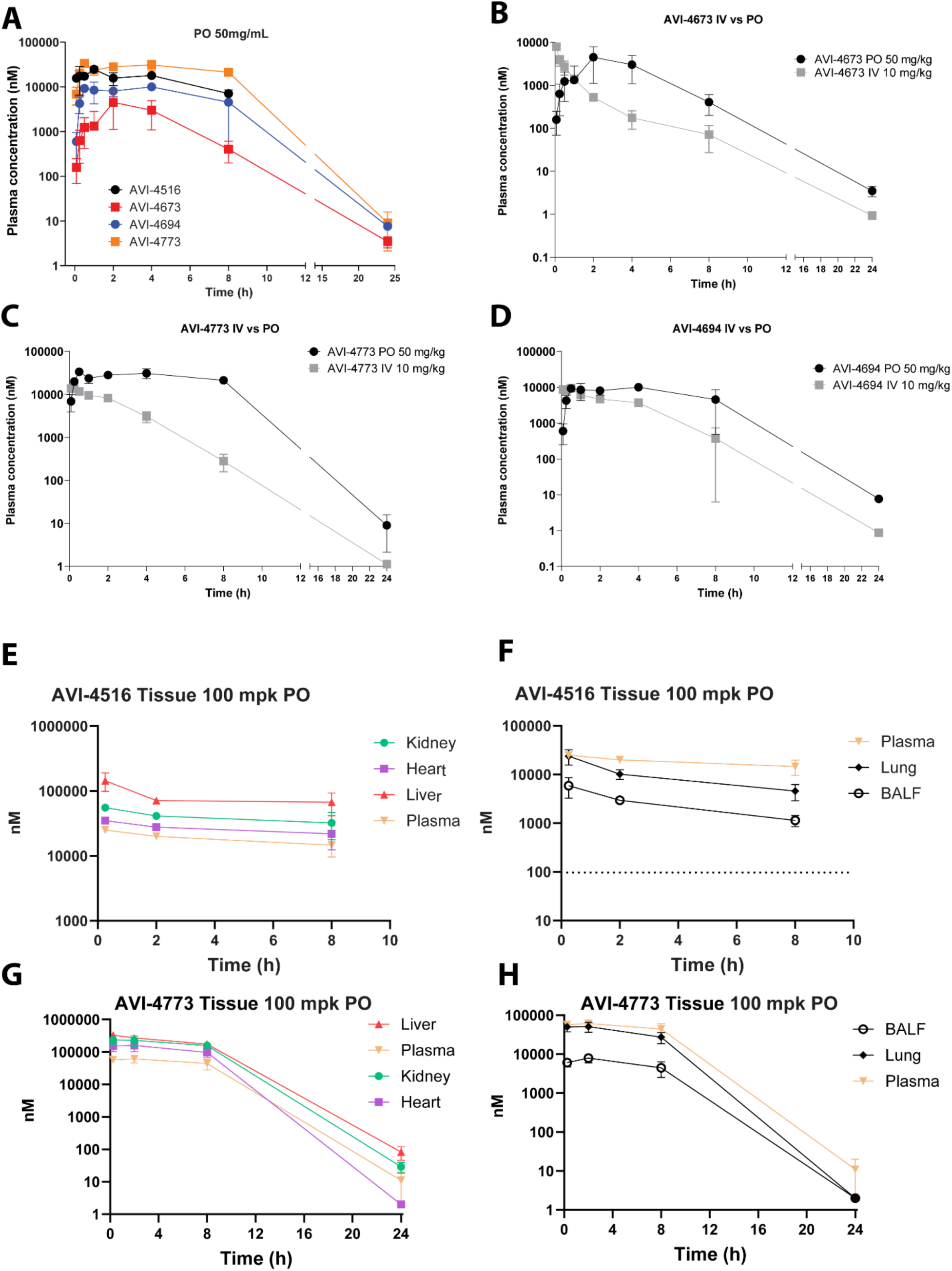
Mouse PK comparison of oral dosing and IV dosing. **A:** comparison of concentration in plasma through oral dosing (50mg/kg) of AVI-4516, AVI-4694, AVI-4673 and AVI-4773. **B-D:** Plasma concentration comparison of PO at 50 mg/kg and IV at 10mg/kg dosing scheme for AVI-4673, AVI-4773, and AVI-4694 respectively. **E:** Kidney, heart, and liver distribution compared to plasma of AVI-4516 after 100 mg/kg PO. Error bars are plotted as ±SD. **F:** Comparison of AVI-4516 (100 mg/kg PO) in lung, plasma and BAL fluid. Error bars are plotted as ±SD. **G**: Kidney, heart, and liver distribution compared to plasma of AVI-4773 after 100 mg/kg PO. **H**: Comparison of AVI-4773 (100 mg/kg PO) in lung, plasma and BAL fluid. Error bars are plotted as ±SD.

**Supplementary Table 4.** Plasma concentration of AVI-4516 after IV (10mg/kg) and PO (50mg/kg) in male CD-1 mice.

| AVI-4516 IV dose 10 mg/kg |  |  |  |  |  |  |  |  | AVI-4516 PO dose 50 mg/kg |  |  |  |  |  |  |
| --- | --- | --- | --- | --- | --- | --- | --- | --- | --- | --- | --- | --- | --- | --- | --- |
| Dose<br>(mg/kg) | Sampling<br>time<br>(h) | Concentration<br>(ng/mL) |  |  | Mean<br>(ng/mL) | SD | CV(%) | Dose<br>(mg/kg) | Sampling<br>time<br>(h) | Concentration<br>(ng/mL) |  |  | Mean<br>(ng/mL) | SD | CV(%) |
|  |  | Individual |  |  |  |  |  |  |  | Individual |  |  |  |  |  |
| 10 IV<br>LLOQ=1.00<br>ng/mL<br>BQL<LLOQ | perdose | BQL | BQL | BQL | **BQL** | NA | NA | 50 PO<br>LLOQ=1.00<br>ng/mL<br>BQL<LLOQ | perdose | BQL | BQL | BQL | **BQL** | NA | NA |
|  | 0.083 | 7850 | 7710 | 7280 | **7613** | 297 | 3.90 |  | 0.083 | 5080 | 6540 | 7490 | **6370** | 1214 | 19.1 |
|  | 0.25 | 4680 | 4520 | 5130 | **4777** | 316 | 6.62 |  | 0.25 | 5630 | 3590 | 12100 | **7107** | 4443 | 62.5 |
|  | 0.5 | 3140 | 3680 | 2250 | **3023** | 722 | 23.9 |  | 0.5 | 8310 | 5590 | 7260 | **7053** | 1372 | 19.4 |
|  | 1 | 2080 | 1350 | 2140 | **1857** | 440 | 23.7 |  | 1 | 9050 | 9280 | 12000 | **10110** | 1641 | 16.2 |
|  | 2 | 615 | 178 | 545 | **446** | 235 | 52.6 |  | 2 | 5560 | 4940 | 8730 | **6410** | 2033 | 31.7 |
|  | 4 | 43.2 | 96.4 | 30.6 | **56.7** | 34.9 | 61.6 |  | 4 | 7520 | 8490 | 6040 | **7350** | 1234 | 16.8 |
|  | 8 | 10.2 | 7.28 | 12.9 | **10.1** | 2.81 | 27.8 |  | 8 | 2630 | 2730 | 3300 | **2887** | 361 | 12.5 |
|  | 24 | BQL | BQL | BQL | **BQL** | NA | NA |  | 24 | BQL | BQL | BQL | **BQL** | NA | NA |
| PK<br>parameters | Unit | Estimated Value |  |  |  |  |  | PK<br>parameters | Unit | Estimated Value |  |  |  |  |  |
| CL | L/hr/kg | 1.74 |  |  |  |  |  | T<sub>max</sub> | hr | 1.00 |  |  |  |  |  |
| V<sub>ss</sub> | L/kg | 1.39 |  |  |  |  |  | C<sub>max</sub> | ng/mL | 10110 |  |  |  |  |  |
| T<sub>1/2</sub> | hr | 0.866 |  |  |  |  |  | T<sub>1/2</sub> | hr | 4.70 |  |  |  |  |  |
| AUC<sub>last</sub> | hr*ng/mL | 5732 |  |  |  |  |  | AUC<sub>last</sub> | hr*ng/mL | 49944 |  |  |  |  |  |
| AUC<sub>INF</sub> | hr*ng/mL | 5744 |  |  |  |  |  | AUC<sub>INF</sub> | hr*ng/mL | 69535 |  |  |  |  |  |
| MRT<sub>INF</sub> | hr | 0.798 |  |  |  |  |  | F | % | 174 |  |  |  |  |  |

**Supplementary Table 5.**
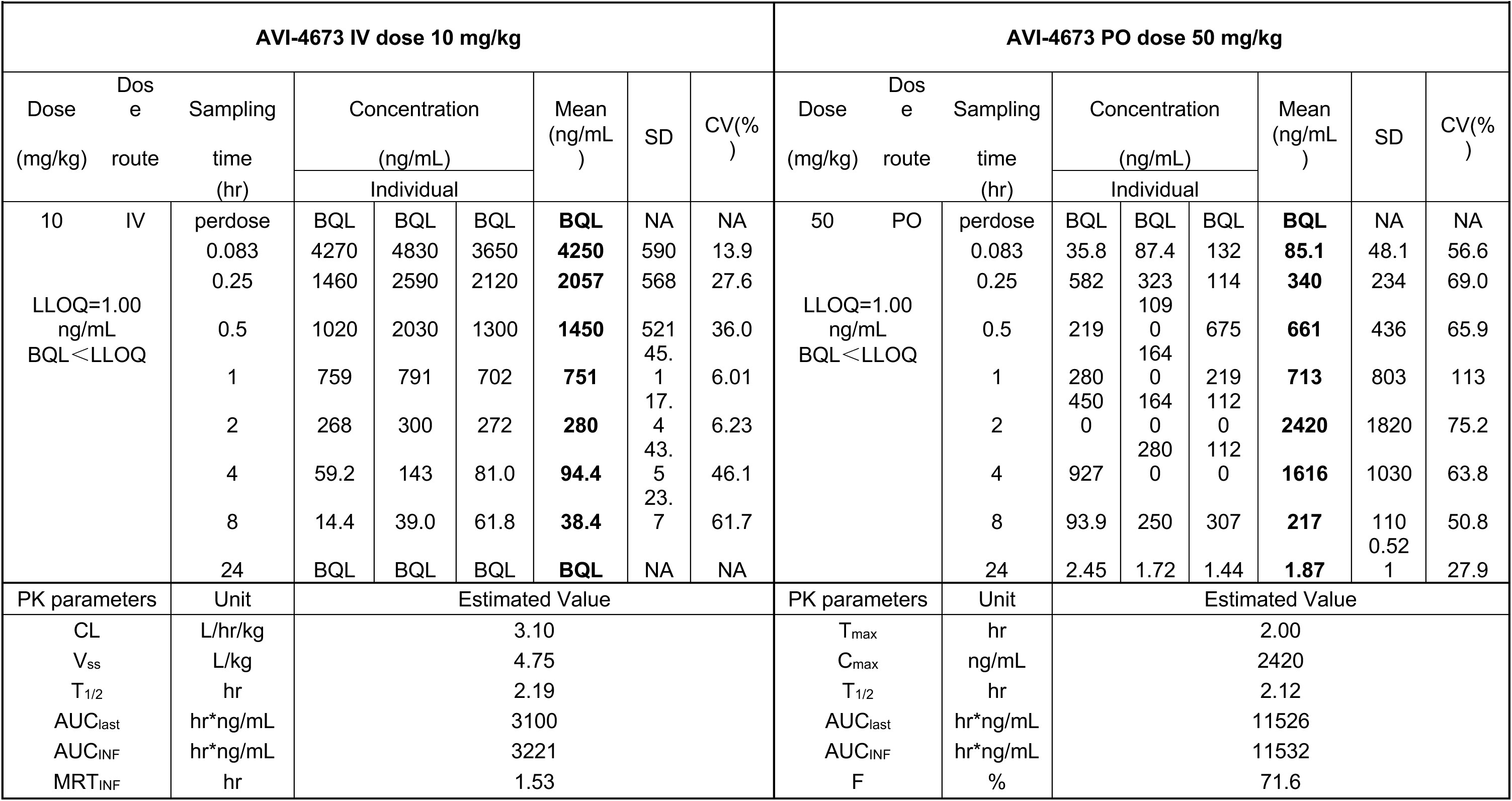
Plasma concentration of AVI-4673 after IV (10mg/kg) and PO (50mg/kg) in male CD-1 mice.

**Supplementary Table 6.** Plasma concentration of AVI-4773 after IV (10mg/kg) and PO (50mg/kg) in male CD-1 mice.

| AVI-4773 IV dose 10 mg/kg |  |  |  |  |  |  |  |  | AVI-4773 PO dose 50 mg/kg |  |  |  |  |  |  |  |  |
| --- | --- | --- | --- | --- | --- | --- | --- | --- | --- | --- | --- | --- | --- | --- | --- | --- | --- |
| Dose<br>(mg/kg) | Dose<br>route | Sampling<br>time<br>(h) | Concentration<br>(ng/mL) |  |  | Mean<br>(ng/mL) | SD | CV(%) | Dose<br>(mg/kg) | Dose<br>route | Sampling<br>time<br>(h) | Concentration<br>(ng/mL) |  |  | Mean<br>(ng/mL) | SD | CV(%) |
|  |  |  | Individual |  |  |  |  |  |  |  |  | Individual |  |  |  |  |  |
| 10 | IV | perdose | BQL | BQL | BQL | **BQL** | NA | NA | 50 | PO | perdose | BQL | BQL | BQL | **BQL** | NA | NA |
|  |  | 0.083 | 6340 | 6270 | 5800 | **6137** | 294 | 4.79 |  |  | 0.083 | 3040 | 4390 | 1760 | **3063** | 1315 | 42.9 |
|  |  | 0.25 | 6080 | 5700 | 6170 | **5983** | 249 | 4.17 |  |  | 0.25 | 8540 | 8950 | 9020 | **8837** | 259 | 2.93 |
|  |  | 0.5 | 4940 | 5640 | 5030 | **5203** | 381 | 7.32 |  |  | 0.5 | 15100 | 14800 | 14900 | **14933** | 153 | 1.02 |
|  |  | 1 | 4110 | 4500 | 4080 | **4230** | 234 | 5.54 |  |  | 1 | 10100 | 13300 | 8230 | **10543** | 2564 | 24.3 |
|  |  | 2 | 3920 | 3180 | 3850 | **3650** | 409 | 11.2 |  |  | 2 | 10800 | 12800 | 13900 | **12500** | 1572 | 12.6 |
|  |  | 4 | 1270 | 1780 | 1030 | **1360** | 383 | 28.2 |  |  | 4 | 14500 | 9920 | 16900 | **13773** | 3546 | 25.7 |
|  |  | 8 | 84.6 | 186 | 103 | **125** | 54.0 | 43.4 |  |  | 8 | 8640 | 9450 | 10300 | **9463** | 830 | 8.77 |
|  |  | 24 | BQL | BQL | BQL | **BQL** | NA | NA |  |  | 24 | 3.35 | 1.34 | 7.35 | **4.01** | 3.06 | 76.2 |
| PK parameters |  | Unit | Estimated Value |  |  |  |  |  | PK parameters |  | Unit | Estimated Value |  |  |  |  |  |
| CL |  | L/hr/kg | 0.574 |  |  |  |  |  | T<sub>max</sub> |  | hr | 0.500 |  |  |  |  |  |
| V<sub>ss</sub> |  | L/kg | 1.18 |  |  |  |  |  | C<sub>max</sub> |  | ng/mL | 14933 |  |  |  |  |  |
| T<sub>1/2</sub> |  | hr | 1.22 |  |  |  |  |  | T<sub>1/2</sub> |  | hr | 13.0 |  |  |  |  |  |
| AUC<sub>last</sub> |  | hr*ng/mL | 17200 |  |  |  |  |  | AUC<sub>last</sub> |  | hr*ng/mL | 94730 |  |  |  |  |  |
| AUC<sub>INF</sub> |  | hr*ng/mL | 17420 |  |  |  |  |  | AUC<sub>INF</sub> |  | hr*ng/mL | 272740 |  |  |  |  |  |
| MRT<sub>INF</sub> |  | hr | 2.06 |  |  |  |  |  | F |  | % | 110 |  |  |  |  |  |

**Supplementary Table 7.** Plasma concentration of AVI-4694 after IV (10mg/kg) and PO (50mg/kg) in male CD-1 mice.

| AVI-4694 IV dose 10 mg/kg |  |  |  |  |  |  |  |  | AVI-4694 PO dose 50 mg/kg |  |  |  |  |  |  |  |  |
| --- | --- | --- | --- | --- | --- | --- | --- | --- | --- | --- | --- | --- | --- | --- | --- | --- | --- |
| Dose<br>(mg/kg) | Dose<br>route | Sampling<br>time<br>(h) | Concentration<br>(ng/mL) |  |  | Mean<br>(ng/mL) | SD | CV(%) | Dose<br>(mg/kg) | Dose<br>route | Sampling<br>time<br>(h) | Concentration<br>(ng/mL) |  |  | Mean<br>(ng/mL) | SD | CV(%) |
|  |  |  | Individual |  |  |  |  |  |  |  |  | Individual |  |  |  |  |  |
| 10 | IV | perdose | BQL | BQL | BQL | **BQL** | NA | NA | 50 | PO | perdose | BQL | BQL | BQL | **BQL** | NA | NA |
|  |  | 0.083 | 4850 | 5280 | 4830 | **4987** | 254 | 5.10 |  |  | 0.083 | 456 | 112 | 468 | **345** | 202 | 58.5 |
|  |  | 0.25 | 4410 | 5030 | 4650 | **4697** | 313 | 6.66 |  |  | 0.25 | 2020 | 1740 | 3580 | **2447** | 991 | 40.5 |
|  |  | 0.5 | 5160 | 5670 | 5420 | **5417** | 255 | 4.71 |  |  | 0.5 | 5490 | 4920 | 5530 | **5313** | 341 | 6.42 |
|  |  | 1 | 2770 | 3580 | 4060 | **3470** | 652 | 18.8 |  |  | 1 | 6730 | 2070 | 5830 | **4877** | 2472 | 50.7 |
|  |  | 2 | 2220 | 2450 | 3420 | **2697** | 637 | 23.6 |  |  | 2 | 4370 | 3630 | 5870 | **4623** | 1141 | 24.7 |
|  |  | 4 | 1710 | 2420 | 2220 | **2117** | 366 | 17.3 |  |  | 4 | 6470 | 5940 | 4890 | **5767** | 804 | 13.9 |
|  |  | 8 | 30.8 | 166 | 441 | **213** | 209 | 98.3 |  |  | 8 | 1460 | 5280 | 1070 | **2603** | 2326 | 89.4 |
|  |  | 24 | BQL | BQL | BQL | **BQL** | NA | NA |  |  | 24 | 3.88 | 4.05 | 5.25 | **4.39** | 0.747 | 17.0 |
| PK parameters |  | Unit | Estimated Value |  |  |  |  |  | PK parameters |  | Unit | Estimated Value |  |  |  |  |  |
| CL |  | L/hr/kg | 0.562 |  |  |  |  |  | T<sub>max</sub> |  | hr | 4.00 |  |  |  |  |  |
| V<sub>ss</sub> |  | L/kg | 1.46 |  |  |  |  |  | C<sub>max</sub> |  | ng/mL | 5767 |  |  |  |  |  |
| T<sub>1/2</sub> |  | hr | 1.72 |  |  |  |  |  | T<sub>1/2</sub> |  | hr | 1.87 |  |  |  |  |  |
| AUC<sub>last</sub> |  | hr*ng/mL | 17270 |  |  |  |  |  | AUC<sub>last</sub> |  | hr*ng/mL | 56507 |  |  |  |  |  |
| AUC<sub>INF</sub> |  | hr*ng/mL | 17797 |  |  |  |  |  | AUC<sub>INF</sub> |  | hr*ng/mL | 56519 |  |  |  |  |  |
| MRT<sub>INF</sub> |  | hr | 2.60 |  |  |  |  |  | F |  | % | 63.5 |  |  |  |  |  |

**Supplementary Table 8.** AVI-4516 tissue distribution after 100 mg/kg PO dose.

| Sampling time (h) | Plasma (ng/mL) |  |  |  | Brain (ng/g) |  |  |  | Kidney (ng/g) |  |  |  |
| --- | --- | --- | --- | --- | --- | --- | --- | --- | --- | --- | --- | --- |
|  | Average |  |  |  | Average |  |  |  | Average |  |  |  |
| 0.25 | 3.57* | 10400 | 10000 | **10200** | 17.1* | 805 | 666 | **736** | 8.86* | 22600 | 22700 | **22650** |
| 2 | 7080 | 9120 | 8370 | **8190** | 542 | 671 | 609 | **607** | 16300 | 18800 | 15400 | **16833** |
| 8 | 6100 | 3870 | 7970 | **5980** | 471 | 262 | 500 | **411** | 12700 | 7500 | 19200 | **13133** |
| Sampling time (h) | Liver (ng/g) |  |  |  | Heart (ng/g) |  |  |  | *=below limit of quantification |  |  |  |
|  | Average |  |  |  | Average |  |  |  |  |  |  |  |
| 0.25 | 38.2* | 72000 | 45700 | **58850** | 9.05* | 15200 | 13400 | **14300** |  |  |  |  |
| 2 | 28700 | 29700 | 29000 | **29133** | 10900 | 12300 | 10800 | **11333** |  |  |  |  |
| 8 | 27100 | 17200 | 38200 | **27500** | 8970 | 5060 | 12900 | **8977** |  |  |  |  |

| Sampling time (h) | Plasma (ng/mL) |  |  |  | Lung (ng/g) |  |  |  |
| --- | --- | --- | --- | --- | --- | --- | --- | --- |
|  | Average |  |  |  | Average |  |  |  |
| 0.25 | 23400 | 18100 | 9870 | **17123** | 13000 | 10000 | 6270 | **9757** |
| 2 | 11000 | 9310 | 7880 | **9397** | 5090 | 4200 | 3200 | **4163** |
| 8 | 3750 | 4380 | 4390 | **4173** | 2650 | 1540 | 1410 | **1867** |
| Sampling time (h) | BALF (ng/mL) |  |  |  | BALF (Sample Volume (mL)) |  |  |  |
|  | Average |  |  |  | Average |  |  |  |
| 0.25 | 3250 | 2810 | 1190 | **2417** | 0.510 | 0.330 | 0.390 | **0.410** |
| 2 | 1420 | 1190 | 1020 | **1210** | 0.800 | 0.710 | 0.750 | **0.753** |
| 8 | 607 | 398 | 396 | **467** | 0.460 | 0.490 | 0.710 | **0.553** |

**Supplemental Table 9.** Brain Distrubition of Ensitrelvir after a 100 mg/kg PO dose in male CD1 mouse.

|  |  |  |  |  |  |  |
| --- | --- | --- | --- | --- | --- | --- |
| Dose<br>(mg/kg) | Dose<br>route | Sampling<br>time<br>(hr) | Concentration<br>(ng/g) |  |  | Mean<br>(ng/g) |
|  |  |  | Individual |  |  |  |
| 100 | PO | 0.25 | 1540 | 1280 | 1970 | 1597 |
|  |  | 2 | 1190 | 1590 | 1780 | 1520 |
|  |  | 8 | 1200 | 914 | 2150 | 1421 |
|  |  | 24 | 29.3 | 46.8 | 67.7 | 47.9 |

**Supplementary Table 10.**
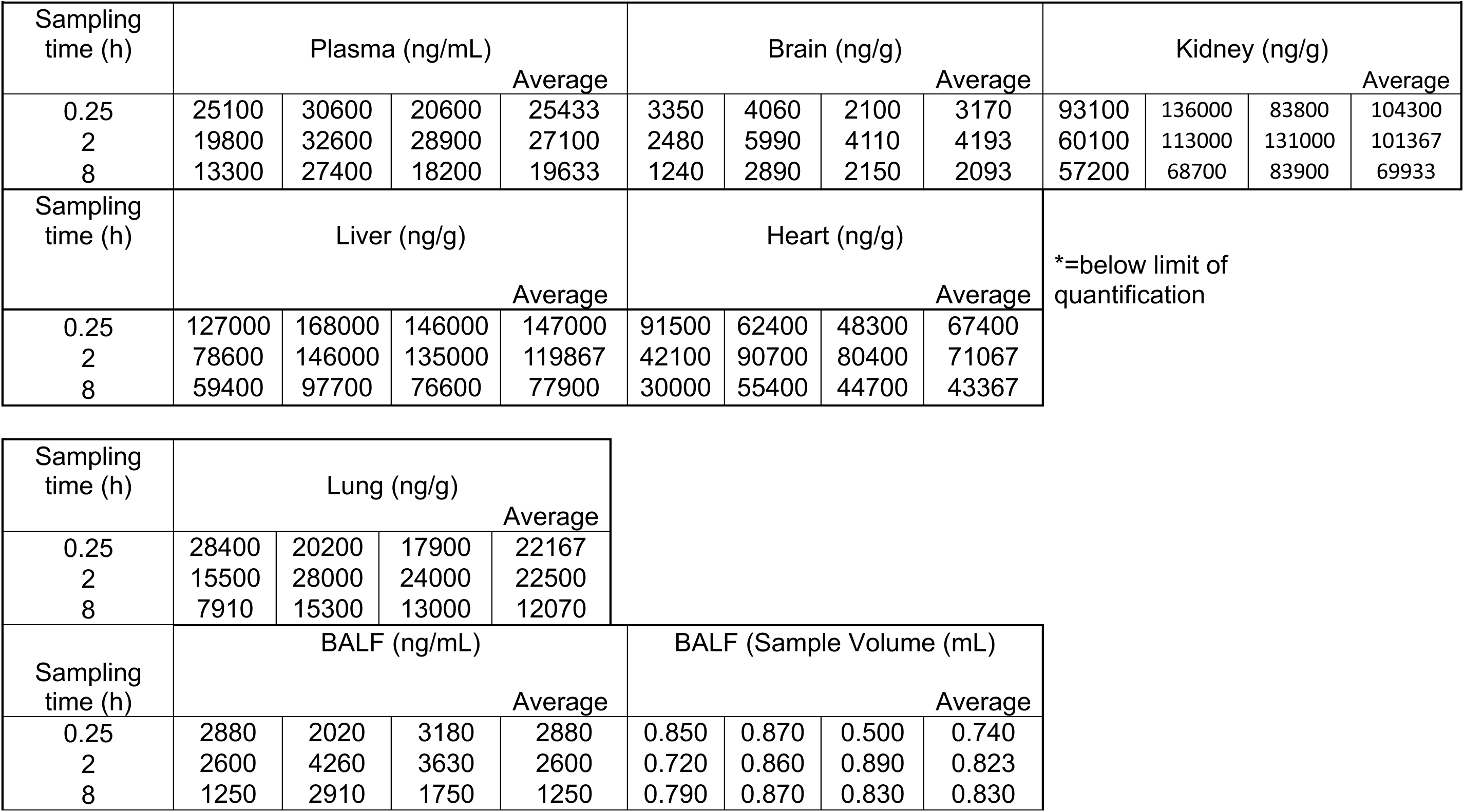
AVI-4773 tissue distribution after 100 mg/kg PO dose.

**Supplementary Fig. 12.**
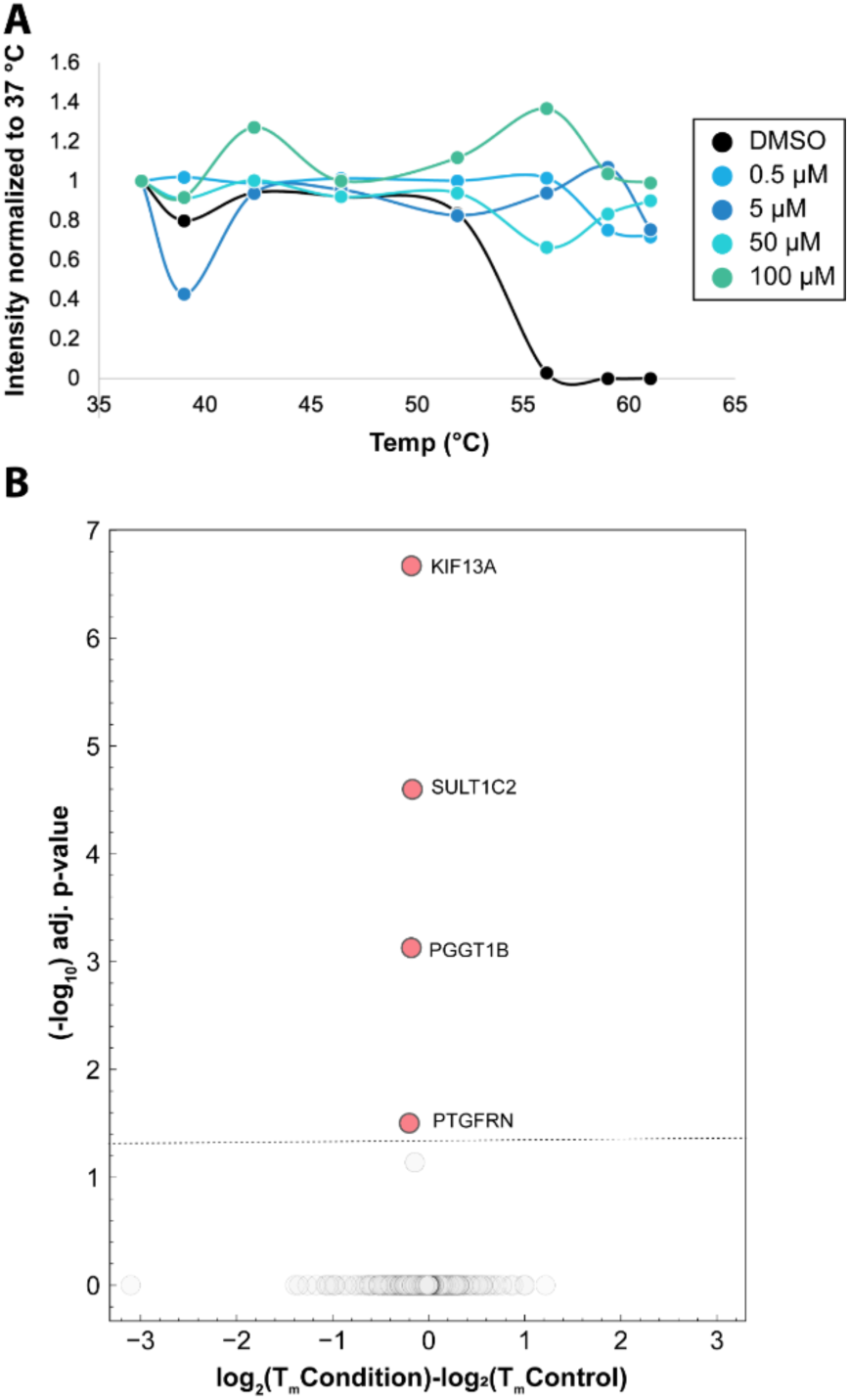
M^Pro^ T_m_ shift and TPP data. **A:** The T_m_ of purified M^Pro^ alone with increasing concentrations of AVI-4516, at even 0.5 µM the compound stabilizes the protein >10 °C. **B:** TPP on lysates treated with **AVI-4516**, there are no significant proteins that exhibit an increased T_m_, and only four proteins that have a statistically significant shift, but very minor changes to the Tm and in the negative direction.

**Supplementary Fig. 13.**
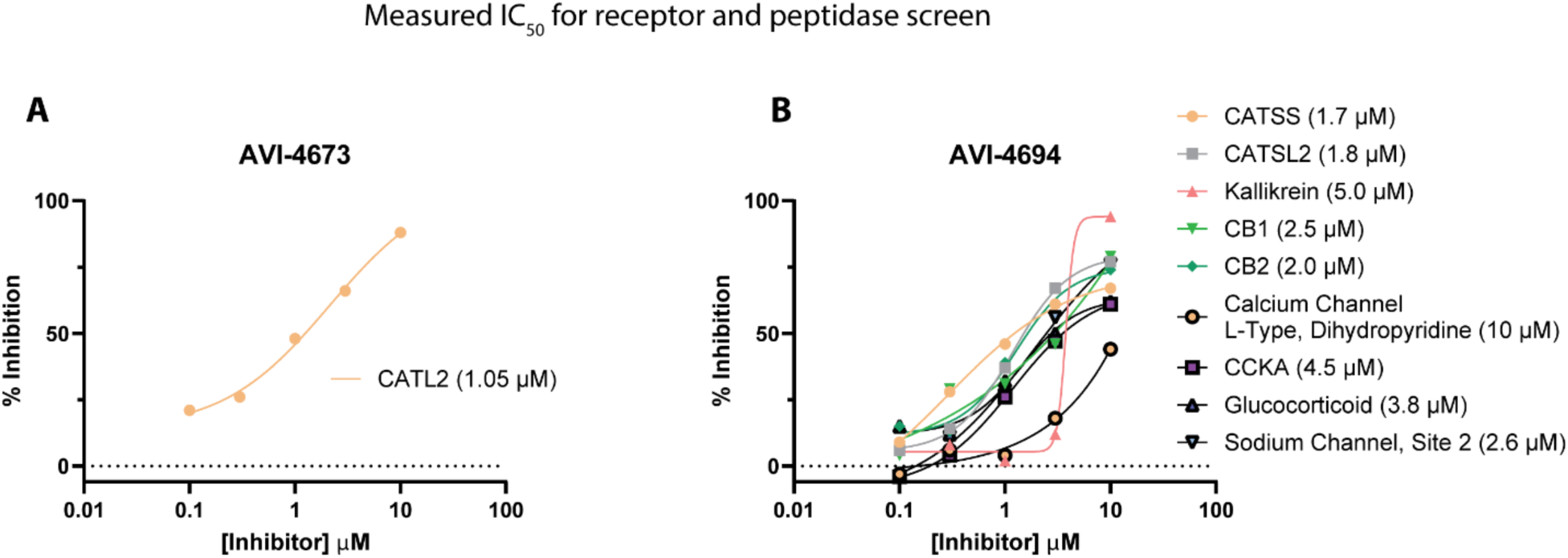
Measured IC_50_s from i*n vitro* safety screen. **A**: AVI-4673 dose response curve for Cathepsin L. **B**: AVI-4694 dose response curve for Cathepsin L, Cathepsin S, Kallikrein, cannabinoid receptor CB1 and CB2, Calcium Channel L-Type, CCK, Glucocorticoid and Sodium channel. Computed IC_50_s are noted in the legend.

**Supplementary Fig. 14.**
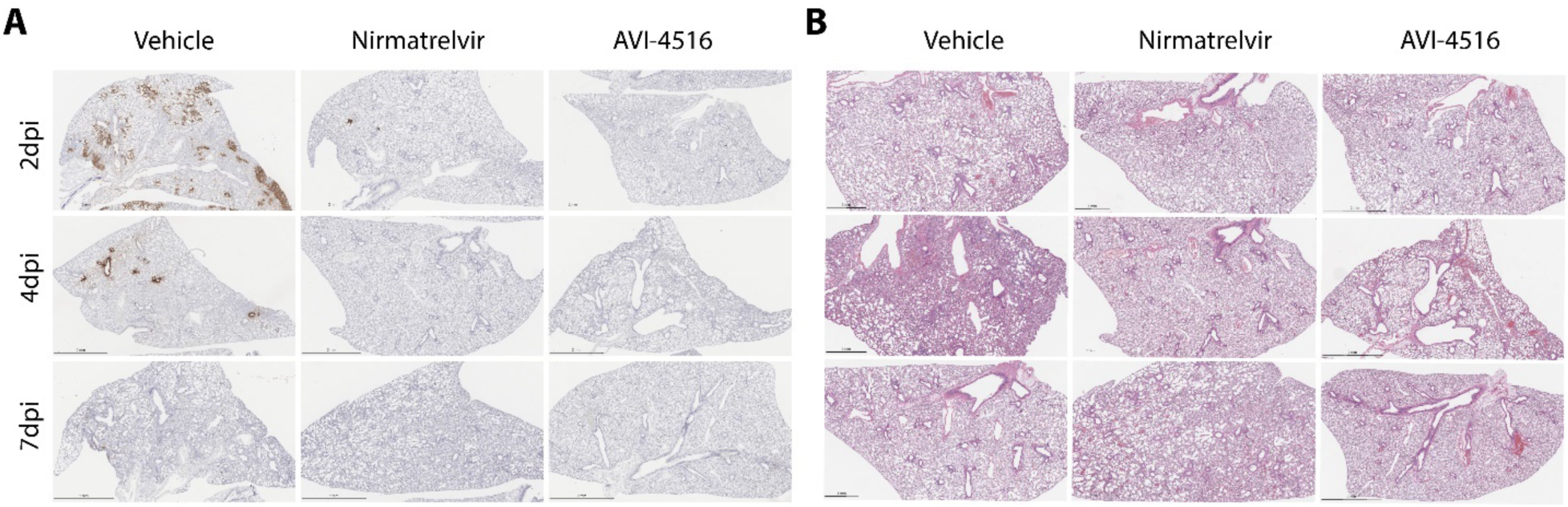
Representative images of immunohistochemistry of the SARS-CoV-2 N protein. (**A**) and Hematoxylin and eosin (H&E) staining (**B**) in the left lung lobe of mice from different treatment groups at the specified time points. Scale bars represent 2mm.

**Supplementary Fig. 15.**
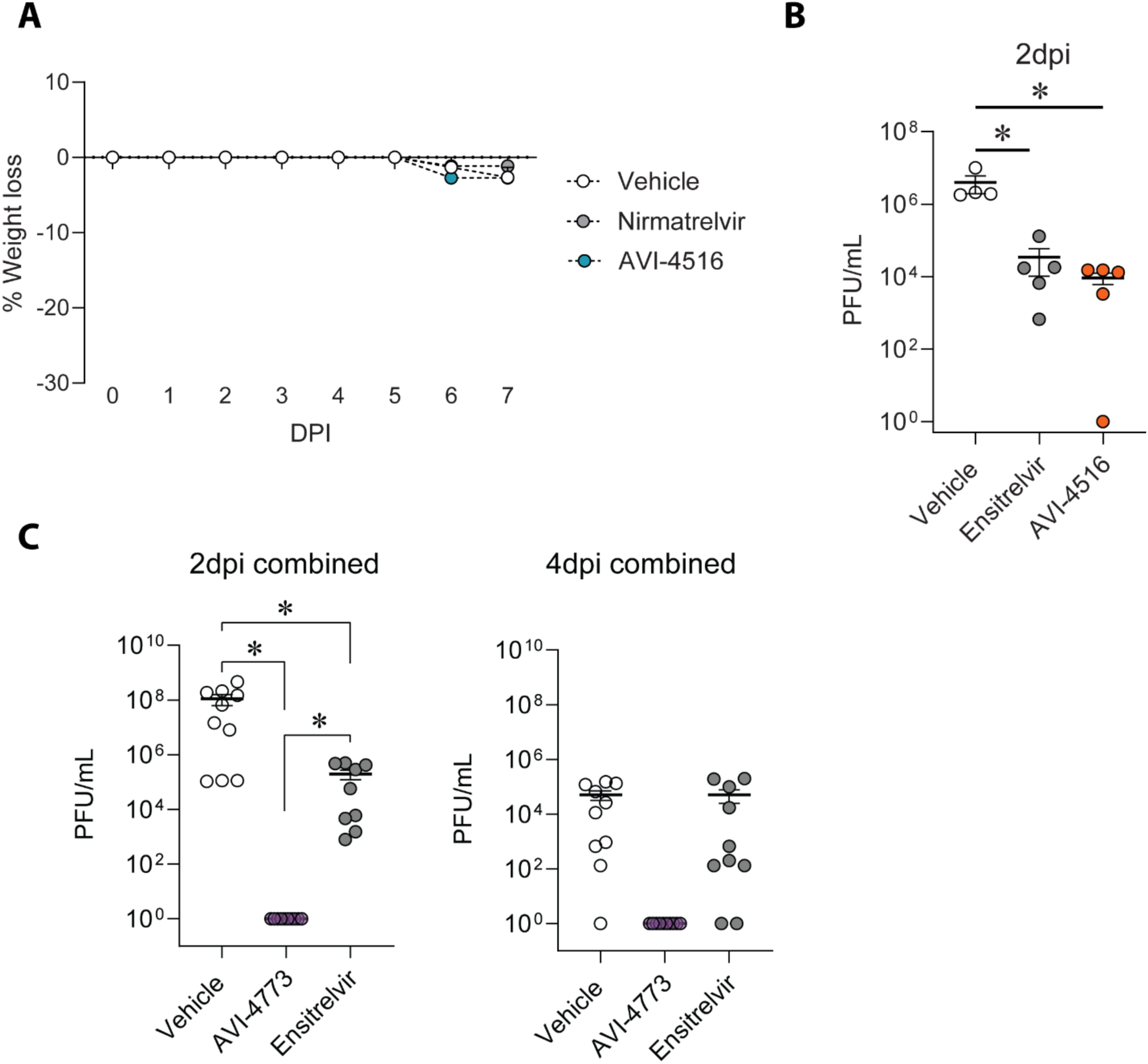
Mouse weight loss and ensitrelvir vs AVI-4516 mouse efficacy experiment. A: Weight loss of mice during antiviral study. Minimal weight loss was observed in all arms of the study. **B:** Comparison of ensitrelvir and AVI-4516 during mouse antiviral study. **C:** Duplicate experiments of AVI-4773 compared to ensitrelvir according to the study design in Fig. 5F combined data points.

**Supplementary Table 11.** Pan-coronavirus activity of M^Pro^ inhibitors.

| Compound Name | Virus | Virus Strain | Cell line | Test Compound |  |  |  |  | Positive Control |  |  |  |  |
| --- | --- | --- | --- | --- | --- | --- | --- | --- | --- | --- | --- | --- | --- |
|  |  |  |  | EC<sub>50</sub> | EC<sub>90</sub> | CC<sub>50</sub> | SI<sub>50</sub> | SI<sub>90</sub> | EC<sub>50</sub> | EC<sub>90</sub> | CC<sub>50</sub> | SI<sub>50</sub> | SI<sub>90</sub> |
| AVI - 4516 | HCoV | Alpha 229E | Huh7 | 2.9 | 3.5 | >100 | >34 | 29 | 0.44 | 0.37 | >10 | >23 | >27 |
| AVI - 4694 |  |  |  | >1.6 | >1.6 | 1.6 | 0 | 0 | 0.44 | 0.37 | >10 | >23 | >27 |
| AVI - 4516 | HCoV | Beta OC43 | Rhabdomyosarcoma | 0.59 | 0.52 | >100 | >170 | >190 | 0.035 | 0.032 | 6.1 | 170 | 190 |
| AVI - 4694 |  |  |  | 0.055 | 0.061 | >100 | >1800 | >1600 | 0.035 | 0.032 | 6.1 | 170 | 190 |
| AVI - 4516 | MERS-CoV | EMC | Vero E6 | <0.032 | 0 | >100 | >3200 | 2900 | 0.0037 | 0.039 | 2 | 540 | 51 |
| AVI - 4694 |  |  |  | 0.045 | 0 | >100 | >2200 | 1300 | 0.0037 | 0.039 | 2 | 540 | 51 |
| AVI - 4516 | SARS-CoV | Urbani |  | 0.57 | 1 | >100 | >180 | >100 | 0.12 | 0.14 | >10 | >83 | >71 |
| AVI - 4694 |  |  |  | 0.052 | 0.091 | 54 | 1000 | 590 | 0.12 | 0.14 | >10 | >83 | >71 |
| AVI - 4516 | SARS-CoV-2 | B. 1. 617.2 (delta) |  | 0.24 | 0.18 | >100 | >420 | 560 | 0.17 | 0.05 | 36 | 210 | 720 |
| AVI - 4694 |  |  |  | 0.063 | 0.047 | 32 | 510 | 680 | 0.17 | 0.05 | 36 | 210 | 720 |
| AVI - 4516 |  | BA.2 (omicron) |  | 0.18 | 0.39 | >100 | >560 | 260 | 0.053 | 0.043 | 43 | 810 | 1000 |
| AVI - 4694 |  |  |  | 0.05 | 0.045 | 41 | 820 | 910 | 0.053 | 0.043 | 43 | 810 | 1000 |

**Supplementary Tale 12:** Refinement statistics for X-ray diffraction data and protein models.

| Compound | AVI-3318 | AVI-4516 | AVI-4692 | AVI-4303 |
| --- | --- | --- | --- | --- |
| PDB entry ID |  |  |  |  |
| Data collection |  |  |  |  |
| Wavelength (Å) | 0.88557 | 1.12709 | 1.12709 | 1.115830 |
| Resolution (Å) | 50.00 - 1.96 | 50.00 - 2.35 | 50.00 - 1.85 | 50.00 - 1.58 |
| Space group | C 1 2 1 | C 1 2 1 | C 1 2 1 | P 32 2 1 |
| Unit cell dimensions |  |  |  |  |
| *a*, *b*, *c* (Å) | 115.5 53.8 45.2 | 115.2 53.7 45.0 | 114.8 53.5 44.8 | 66.4 66.4 233.6 |
| $\alpha$, $\beta$, $\gamma$ (°) | 90 101 90 | 90 101.5 90 | 90 102.1 90 | 90 90 120 |
| Total number of reflections (<sup>a</sup>) | 475284 (27923) | 66793 (10390) | 155577 (25174) | 1159359 (186246) |
| Unique reflections (<sup>a</sup>) | 17848 (1774) | 11344 (1752) | 23010 (2253) | 84920 (13483) |
| Multiplicity (<sup>a</sup>) | 10.06 (15.74) | 5.88 (5.93) | 6.18 (6.29) | 13.6 (13.8) |
| Completeness (%) (<sup>a</sup>) | 90 (56) | 99.1 (96.0) | 99.6 (98.8) | 99.9 (99.4) |
| Mean *I*/$\sigma$(*I*) (<sup>a</sup>) | 45.13 (2.96) | 13.39 (1.96) | 21.74 (2.57) | 25.88 (3.31) |
| *R*<sub>merge</sub> (%) (<sup>a</sup>) | 4.7 (83.5) | 10.1 (81.0) | 5.3 (77.6) | 5.7 (76) |
| *R*<sub>meas</sub> (%) (<sup>a</sup>) | 4.8 (86.2) | 11.1 (88.9) | 5.7 (83.9) | 6.0 (78.9) |
| *CC*<sub>1/2</sub> (%) (<sup>a</sup>) | 100 (89) | 99.7 (72.1) | 100 (86.4) | 100 (91.6) |
| Wilson B-factor | 40.22 | 47.29 | 30.61 | 21.14 |
| Refinement |  |  |  |  |
| Resolution (Å) (<sup>a</sup>) | 44.44 - 1.96 (2.03 - 1.96) | 48.59 - 2.35 (2.43 - 2.35) | 38.68 - 1.843 (1.91 - 1.8) | 46.29 - 1.57 (1.62 - 1.57) |
| Reflections used (<sup>a</sup>) | 17809 (962) | 11340 (1075) | 23012 (2253) | 84857 (8380) |
| Reflections used for *R*<sub>free</sub> (<sup>a</sup>) | 862 (42) | 1135 (108) | 2000 (196) | 4166 (371) |
| *R*<sub>work</sub> (<sup>a</sup>) | 0.20 (0.29) | 0.18 (0.28) | 0.17 (0.28) | 0.15 (0.17) |
| *R*<sub>free</sub> (<sup>a</sup>) | 0.24 (0.35) | 0.20 (0.32) | 0.22 (0.38) | 0.20 (0.26) |
| Number of non-hydrogen atoms | 2407 | 2420 | 2591 | 5376 |
| macromolecules | 2353 | 2335 | 2317 | 4774 |
| ligands | 23 | 64 | 76 | 73 |
| solvent | 31 | 21 | 198 | 529 |
| Protein residues | 305 | 305 | 305 | 606 |
| RMS (bonds) | 0.049 | 0.047 | 0.036 | 0.042 |
| RMS (angles) | 1.17 | 1.68 | 2.10 | 1.08 |
| Ramachandran plot |  |  |  |  |
| favored (%) | 97.67 | 98.00 | 98.33 | 97.84 |
| allowed (%) | 2.00 | 1.67 | 1.67 | 2.16 |
| outliers (%) | 0.33 | 0.33 | 0.00 | 0.00 |
| rotamer outlier(%) | 0.00 | 1.17 | 0.00 | 0.37 |
| Clashscore | 5.35 | 6.43 | 5.19 | 2.72 |
| Average B factor | 45.71 | 54.62 | 33.58 | 29.73 |
| macromolecules | 45.76 | 54.78 | 32.93 | 28.76 |
| ligands | 43.59 | 51.48 | 33.48 | 31.23 |
| solvent | 43.33 | 46.87 | 41.22 | 38.31 |

<sup>(a)</sup> Values in parentheses correspond to the highest-resolution shell

**Supplementary Fig. 16.**
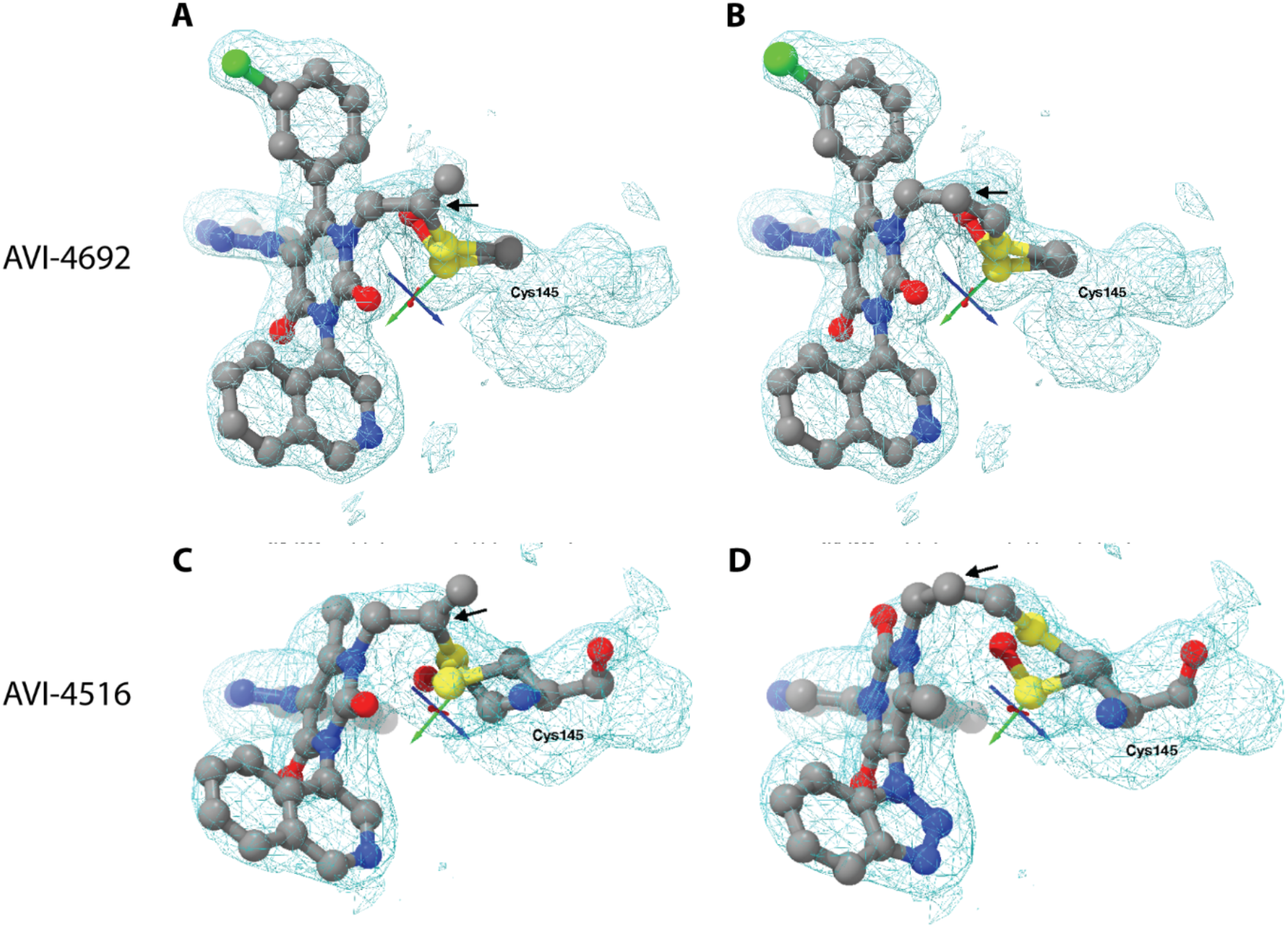
Comparison of density maps for cysteine connectivity to AVI-4692 and AVI-4516. **A-B:** Omit electron density maps were generated by omitting any modeled ligand, then AVI 4692 reacting either at its internal carbon (a) or terminal carbon (b) in their final refined states were fit into this omit density map (shown at 1 sigma). The internal carbon in both cases is shown with an arrow. Configuration while reacting with internal carbon shows marginally better fit R_work_ 0.1742 R_free_ 0.2214 for internal; R_work_ 0.1803 R_free_ 0.2238 for external.). **C-D:** comparing different binding modes for 4516. Omit electron density maps were generated by omitting any modeled ligand, then AVI 4516 reacting either at its internal carbon (c) or terminal carbon (d) in their final refined states were fit into this omit density map (shown at 0.7 sigma). The internal carbon in both cases is shown with an arrow. Configuration while reacting with internal carbon shows marginally better fit (R_work_ 0.1807 R_free_ 0.2034 for internal; R_work_ 0.1828 R_free_ 0.2047 for external.). Partial occupancy oxidized cysteine 145 is also shown in all structures.

**Supplementary Figure 17.**
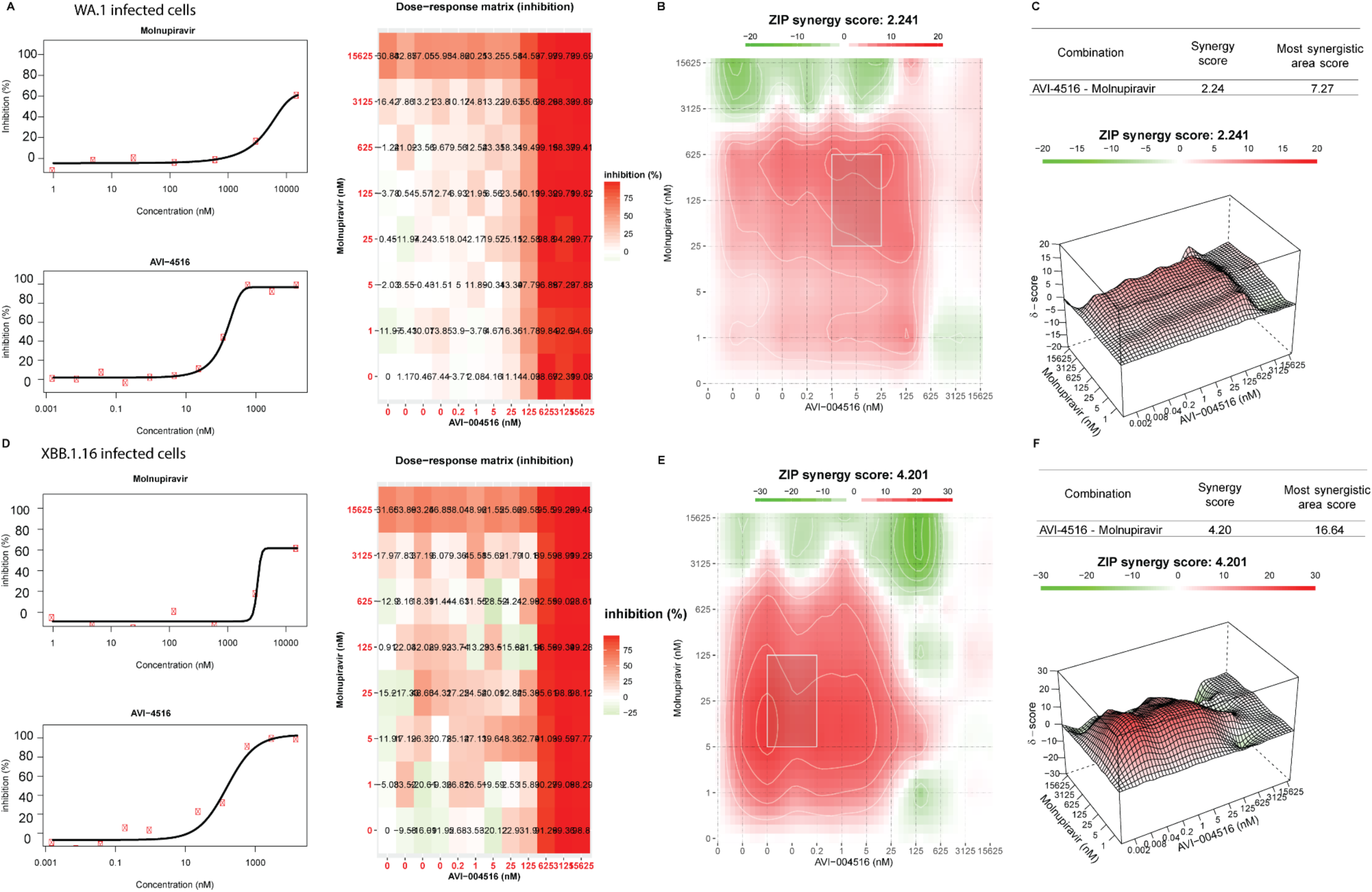
Synergy experiments with AVI-4516 and molnupiravir. **A-C**. Synergy experiments with AVI-4516 and molnupiravir with WA.1 infected cells. **A**: Dose response for each compound tested and in matrix format. **B**: 2-D plot of ZIP analysis of synergy C: 3-D plot of ZIP analysis of synergy with computed overall synergy score and highest area score. **D-F**: Synergy experiments with AVI-4516 and molnupiravir with XBB.1.16 infected cells. **D:** Dose response for each compound tested and in matrix format. **E**: 2-D plot of ZIP analysis of synergy **F**: 3-D plot of ZIP analysis of synergy with computed overall synergy score and highest area score.

## Materials and Methods

### Non-covalent optimization

Analogs for docking hit Z3535317212 were queried in SmallWorld 48 billion make-on-demand libraries (https://swp.docking.org/search.html). The resulting analogs were further filtered based on Tc > 0.5 and docked to the M^Pro^-x11612 as described in the previous docking campaign(*12*). Compounds were also designed by modifying the 2D structure and custom synthesis by Enamine Ltd. (Kyïv, Ukraine). The docked poses were visually inspected for compatibility with the site, and prioritized analogs were synthesized and tested. Make-on-demand non-covalent analogs were purchased and synthesized by Enamine Ltd. Purities of molecules were at least 90% and most active compounds were at least 95% (assessed by LC/MS data).

### M^Pro^ expression and purification

The M^Pro^ expression plasmid was generated as previously described(*12*) with slight modifications. BL21 pLyS Ros2 (DE3) cells were transformed with the expression plasmid. A single colony was used to start an overnight culture in LB media supplemented with 100 µg/mL ampicillin and 20 µg/ml of chloramphenicol. This overnight culture was diluted 1:50 to inoculate 2XYT media supplemented with 100 µg/mL ampicillin and 20 µg/ml chloramphenicol. These cultures grew at 37 °C until the OD600 reached approximately 1-2.0, at which point the temperature was reduced to 20 °C and IPTG was added to a final concentration of 1 mM induced overnight. The cultures were then harvested by centrifugation at 4,000 x g for 20 min at 4 °C. The pellet was then resuspended in 50 mM Tris pH 8.0, 400 mM NaCl, 20 mM imidazole with 0.1 mg/mL DNAseI and 0.1 mg/mL Lysozyme (Sigma). The resuspended pellet was lysed by sonication and then clarified with centrifugation at 42,000 x g for 30 min. The clarified lysate was loaded onto pre-equilibrated Ni-NTA resin (with Buffer A (400 mM NaCl 50 mM Tris 20 mM imidazole pH 8.0). The column was then washed with 20X CV of Buffer A and eluted in a step gradient of imidazole from 20 – 500mM. The fractions were analyzed with SDS-PAGE for purity. Pure fractions were pooled and concentration to <1 mL treated with his tagged HRV 3C protease (expressed as described previously(*12*)) in the ratio of 1 mg HRV 3C protease/ 50 mg of M^Pro^ (by A_280_) to cleave the histidine tag. This solution was then dialyzed overnight into HRV3C buffer (50mM Tris, pH 7.0 150 mM NaCl, 1 mM EDTA, 1 mM DTT at RT overnight. Room temperature dialysis reduced precipitation as reported previously(*13*). The dialyzed and cleaved protein was then flown over a Ni NTA column equilibrated with Buffer A. The column was washed with 5 CV of Buffer A and 5 CV of Buffer A with 500 mM imidazole. Tagless M^Pro^ was eluted in the flowthrough and concentrated using 10 kDa MWCO Amicon ultra centrifugal filters (Millipore Sigma). This protein was then passed over an S200 (GE) column in Buffer B (25 mM HEPES 150 mM NaCl 1 mM TCEP pH 7.5). Purified M^Pro^ was concentrated to ∼10mg/mL and stored at -80 for months with negligible loss of activity. The variants E166Q, S144A, Q192T, A173V were generated as previously described(*12*) and purified in the same manner as WT M^Pro^.

### M^Pro^ inhibition Assays

All M^Pro^ enzymatic assays used Buffer C (50 mM Tris 150 mM NaCl 1 mM EDTA 0.05% Tween-20 1 mM TCEP pH 7.4). TCEP was added fresh for each assay and the pH was readjusted to 7.4 upon addition. Enzyme was incubated in Buffer C with TCEP for 10 min to ensure full activation of M^Pro^. The compounds were then incubated with M^Pro^ at RT for 1h on Corning 3820 384 well black plates. After incubation with compounds, the M^Pro^ substrate, (dR)(dR)(MCA)KATVQAIAS(DNP)K which was synthesized as previously described(*12*) was used to initiate the reaction at a final concentration of 10 µM. The final DMSO in each well was 1%. Increase in fluorescence with an excitation of 328 nm and an emission of 393 nm over time was monitored for the first 30 min of the reaction with a BioTek Neo2 plate reader. All curves were performed in at least technical triplicate. Each slope was normalized to M^Pro^ with DMSO only control. These dose response curves were fitted to a four-parameter inhibitor vs response curve (IC_50_) curve in GraphPad Prism 10.2.0. For compounds that displayed an IC_50_ value > ∼1 µM, 50 nM of M^Pro^ was used in the assay. For compounds that displayed greater potency 25 nM M^Pro^ was used as enzyme concentrations lower than 25 nM resulted in high noise. All mutated SARS-CoV-2 IC_50_ assays were performed at 50 nM. SARS-CoV-1 M^Pro^ IC_50_s were measured using the same substrate and buffer conditions as the SARS-CoV-2 M^Pro^ assay.

### Inhibitor kinetics assays

A twelve-point serial dilution (starting either 1.5 dilution from 3 µM or 1.2 dilution from 100 nM) of inhibitor and DMSO only was prepared in DMSO and diluted in Buffer C to 3x the final concentration. This was added to an equal volume of a 3x solution of substrate (30 uM) diluted in buffer C. 10 µl of a 3x solution of M^Pro^ diluted in Buffer C was then add to a Corning 3820 384 well black plate. Both the substrate and inhibitor mixture and the enzyme on the plate was allowed to warm at 37 °C for 30 min to ensure accuracy of reads and full activation of M^Pro^. The substrate and inhibitor mixture was then added to the plate and increase in fluorescence was monitored with an excitation of 328 nm and an emission of 393 nm on a BioTek Neo2 plate reader. All assays were run in technical quintuplicate. An average trace of blank containing substrate alone was then subtracted from all traces. The traces were then fit similar to previous reported(*14*) in GraphPad Prism software version 9.1.1 using Y=(((V)/k)*(1-exp(-k_app_*x)))+Z to obtain *k_app_* at each inhibitor concentration. These values were then plotted vs inhibitor concentration and fit using the Michaelis Menten fitting equation in Prism to obtain *k_inact_* and *K_I_*. All inhibitors tested were compared against 25 nM due to high noise and low signal with lower concentrations of M^Pro^ and thus For AVI-4773, AVI-4692, AVI-4694 inhibitor saturation could not be observed and the resulting *k_app_* data were fit using a linear regression in prism to obtain the *k_inact/_K_I_* values from the slope. The values were corrected for substrate in the assay using previously described equations(*14*).

### Enzyme Aggregation inhibition assays

Samples were prepared in 50 mM KPi buffer, pH 7.0 with final DMSO concentration at 1% (v/v). Compounds were incubated with 2 nM Malate dehydrogenase (MDH) (Sigma Aldrich, 442610) or AmpC β-lactamase (AmpC) for 5 minutes. MDH reactions were initiated by the addition of 200 μM nicotinamide adenine dinucleotide (NADH) (Sigma Aldrich, 54839) and 200 μM oxaloacetic acid (Sigma Aldrich, 324427). The change in absorbance was monitored at 340 nm for 80 s. AmpC reactions were initiated by the addition of 50 μM CENTA chromogenic substrate (Sigma Aldrich, 219475). The change in absorbance was monitored at 405 nm for 80 s. Initial rates were normalized with the DMSO control to determine percent enzyme activity (%). Each compound was initially screened at 10 μM in triplicate. Compounds that did not inhibit MDH but formed colloidal-like particles by DLS were screened against AmpC. Data was analyzed using GraphPad Prism software version 9.1.1 (San Diego, CA).

### Dynamic light scattering (DLS)

Samples were prepared in filtered 50 mM KPi buffer, pH 7.0 with final DMSO concentration at 1% (v/v). Colloidal particle formation was detected using DynaPro Plate Reader III (Wyatt Technologies). All compounds were screened in triplicate at 10 μM. If colloidal-like particles were detected, seven-point half-log dilutions of compounds were performed in triplicate. As previously reported(*15*), critical aggregation concentrations (CACs) were determined. Analysis was performed with GraphPad Prism software version 9.1.1 (San Diego, CA).

### Intact Protein Mass Spectrometry

500 nM of M^Pro^ was incubated with DMSO only, 100 µM of AVI-4516, or AVI-4694 in 50 mM Ammonium Acetate 1 mM TCEP pH 7.4 with a final DMSO concentration of 1% for 24 h. 8 µL of this reaction was then injected onto an I-Class Acquity UPLC (Waters) equipped with an Acquity UPLC protein BEH C4 column (Waters). Mass spectra were measured by a Xevo G2-XS Quadrupole Time of Flight mass spectrometer with a ZSpray ion source. The gradient and mass spectrum collection were performed as described previously(*16*). For comparison of modified proteins, the spectra were deconvoluted using MaxEnt1 software and the resulting data were visualized in Prism 10.

### Determination of modified residue for covalent inhibitors 4516 and 4694 using chymotryptic digestion

10 µM MPro was incubated in 50 mM ammonium acetate 5 mM DTT pH 7.4 for 20 h at RT with either 100 µM of compound AVI-4516 or AVI-4694. The protein was then denatured with addition of Guanidinum HCL (Sigma Aldrich) to a final concentration of 1 M and heated at 60 °C for 20 min then alkylated with 15 mM of iodoacetamide and digested according to the manufacturer’s protocol for chymotrypsin (Promega). These samples were desalted using preequilibrated (3 x 15 µL of 50% ACN 0.2% Formic acid then 3 x 15 µL 0.2% formic acid) Cleanup C18 pipette tips (Agilent) by pipetting 15 µL of the acidified solution 10X to ensure full binding of the peptides to the C18 plug in the tips. The tips were then washed 5 x with 15 µL of 0.2% formic acid, and finally with 5 x 15 µL of 50% ACN 0.2% Formic acid eluted into a to a nonstick 0.5 ml Axygen maximum recovery tube (Corning). This eluant was then evaporated in a speed vac and reconstituted with 15 µL of 0.1% formic acid. 5 µL of each sample was then injected into a PepMap RSLC C18 (Thermo Scientific ES900) attached to a 10,000-psi nanoACQUITY Ultra Performance Liquid Chromatography System (Waters) followed by a Q Exactive Plus Hybrid Quadrupole-Orbitrap (Thermo Fisher Scientific). The peaks were assigned using PAVA and the peaks were searched using ProteinProspector for any modification on cysteine.

### Reactivity of AVI-4516 with betamercaptoethanol

A 4 mM solution of AVI-4516 was prepared in 0.5 mL DMSO-d_6_ and analyzed by ^1^H-NMR as T_0_. Then, 0.5 mL of 40 mM BME-d_4_ in DMSO-d_6_ was added to the compound and incubated at room temperature. The sample was analyzed by ^1^H-NMR after 1 and 24 hours. The 3 obtained spectra were aligned and then stacked for comparison.

### PAMPA Assay

Parallel artificial membrane permeability assay (PAMPA) measurements were made using the PAMPAExplorer kit (pION, PN 120670-10) as described by the manufacturer. Prisma HT Buffer (pION, PN110151) at pH 7.4 was added to each well of the 96-well High Sensitivity UV Plate (pION, PN 110286) and the UV absorption was read from 250nm to 500nm using 10 nm steps using the Molecular DevicesFlexStation 3 Multi-Mode Microplate Reader to obtain the baseline signal. Once the compounds were added to the 96 well deep well plate (pION, PN 110023) as directed by the kit, then the plate was agitated for 1 h at 1000 rpm. Afterwards, the contents of the plate were transferred to a 96 well filter plate (AcroPrep™ Advance, PN 8129) with an empty deep well plate underneath and spun down in a centrifuge. The filtered solutions were then transferred to a UV plate and read as described above to determine the initial signal. The PAMPA plate sandwich was then prepared as directed in the kit. 3 controls as a reference of permeation speed and DMSO as a blank. All the compounds were done in technical triplicate. The GIT-0 lipid solution (pION, PN 110669) was used to mimic the gastrointestinal tract (GIT) conditions. The control references for the GIT assay were Verapamil for high permeability, Antipyrine for low/moderate permeability, and Ranitidine for low permeability. After 15 min, the wells of the acceptor plate were filled with the acceptor sink buffer (ASB) for the gut PAMPA as described in the kit. The sandwich was then placed in a humidity-controlled chamber and incubated at room temperature for 18 h without any well stirrers. After the 18-hour incubation, the contents from the plates were transferred to the UV plate. The permeation speed was then determined in the PAMPA Explorer software using the initial, baseline, and final measurements for each well.

### Cell Cytotoxicity Assay

A549-ACE2h were used for the cytotoxicity assay. Briefly, 2×10^4^ cells/well were seeded in Nunc Edge 2.0 96-well plates (Thermo Scientific) filled with 1.5 mL PBS for outer moats and 100 µl for in-between wells and incubated for 24 h at 37°C and 5% CO_2_. Next, cells were treated with compounds at the respective concentrations and vehicle control for 50 h at 37°C and 5% CO_2_. After the incubation, Cell Titer-Glo^Ⓡ^ reagent was added 1:1 to cells and incubated at rt for 5 min prior transfer of 100 µl of mixture to a white 96-well plate. Luciferase was measured in an infinite M Plex plate reader (Tecan). Cell viability was analyzed as the percentage of viability normalized to the vehicle control. Compound cytotoxicity was assessed in parallel to infection experiments with cells of the same passage.

Compound plates were created using an Echo acoustic dispenser with a final DMSO concentration of 0.5% in a 782080 Greiner 384-well plate. 2000 A540 cells in 25 µL of media were added to each well. The plate was incubated at 37°C under a 5% CO2 atmosphere for 48 hours followed by the addition of 25 µL of Cell-Titer glo. Percent viability was measured using a PerkinElmer Envision plate reader. Data processing was completed using GraphPad Prism software version 9.1.1 with a 4-PL logistic curve fit with DMSO only control set as 100% viability and media only control as 0%.

### Cells and viruses

A549-ACE2h were generated by stable expression and selection for hACE2 expression(*17*) followed by sorting of cells expressing high levels of the receptor by FACS using a hACE-2 Alexa Fluor® 647-conjugated mab (FAB9332R, R&D systems). Cells were maintained with DMEM supplemented with 10% FBS, blasticidin (10 μg/ml) (Sigma), 1X NEAA (Gibco), and 1% L-Glutamine (Corning) at 37°C and 5% CO2. Vero-ACE2/TMPRRS2 (VAT) (gifted from A. Creanga and B. Graham at NIH) were maintained in DMEM supplemented with 10% FBS, 1x Penicillin-Streptomycin, and 10 μg/mL of puromycin at 37°C and 5% CO2.The mNeonGreen SARS-CoV-2 (icSARS-CoV-2-mNG) was a kind gift from <u>Pei-Yong Shi</u> (University of Texas Medical Branch, Galveston). Virus was propagated in VAT cells and viral sequence verified.

### SARS-CoV-2 replicon assay

SARS-CoV-2 single-round infectious particles were generated as previously described with some modifications(*18*). BHK-21 cells were seeded in 10-cm dish (1×10^6^) and were transfected the next day 10 µg pBAC SARS-CoV-2 Spike replicon plasmid (WA1, WA1 nsp5 L50F/E166Q/L167F, or BA.2.86.1), 5 µg Spike Delta variant plasmid(*19*), and 5 ug Nucleocapsid R203M plasmid(*20*) using Xtremegene 9 DNA transfection reagent (Sigma Aldrich). The media was changed the next day, and the cells were incubated at 37 °C and 5% CO2. At 70 hours post transfection, 20K VAT cells in 50 µL culture medium were mixed with 50 µL compound at 4x final concentration and plated in 96-well tissue culture plates. At 72 hours post transfection, the supernatant was 0.45 µm filtered and 100 µL was added to each well of compound treated VAT cells and the cells were incubated for 6-8 hours at 37 °C and 5% CO2. The cells were washed once with culture medium and 100 µL of compound containing culture medium was added. The cells were incubated for 24 hours and 50 µL of supernatant was transferred to white 96-well plate. 50 µL of Promega nanoGlo reagent was added and luminescence was recorded in a Tecan plate reader. Experiments were conducted in two biological replicates.

### In cell drug antiviral screening and dose-dependent curves

Compound antiviral activity was determined using the Incucyte^Ⓡ^ live cell analysis system. A549-ACE2h cells were seeded and incubated as for the cytotoxicity assay. The next day, cells were pre-treated with compounds for 2 h prior to removal of compounds and infection with the mNeon expressing viruses icSARS-CoV-2-mNG (MOI 0.1), SARS-CoV-2/XBB.1.5 (MOI 0.13), SARS-CoV-2/XBB.1.16 (MOI 1), or SARS-CoV-2/EG.5.1 (MOI 0.13) Cells were infected with 50 µl viral inoculum for 2 h before removal and addition of fresh compounds and controls. Fresh compounds and controls were diluted in DMEM complete (10% FBS, 1% L-Glutamine, 1X P/S, 1X NEAA) supplemented with Incucyte® Cytotox Dye (4632, Sartorius) to control for cell death. After addition of fresh compounds, infected cells were placed in an Incucyte S3 (Sartorius) and infection/cell death measured for 48 h in 1 h intervals using a 10x objective and capturing 3 images/well per time point under cell maintenance conditions (37°C, 5% CO2). Infection was quantified as Total Green Object Integrated Intensity (GCU x µm^2^/Image) with an acquisition time of 300 ms and cell death as Red Object Integrated Intensity (GCU x µm^2^/Image) for 400 ms. Image analysis for measurements were done with the following parameters: Phase, AI confluence segmentation. Green, Top-hat segmentation with a 50 µM Radius, GCU threshold of 0.5, and Edge Split On. Red was similar to Green with a 100 µM radius and a threshold of 1 RCU. A 2 % spectral unmixing of the red channel into the green was predefined to prevent signal spillover. Post in-built software analysis, raw data was exported and antiviral efficacy determined as the percentage of infection normalized to the vehicle control. A positive control (Nirmatrelvir, HY-138687, MedChemExpress) at efficacious concentrations and uninfected cells were used as an intra-assay positive and negative control. Unless otherwise stated, experiments were performed in triplicate with 3 technical replicates. EC_50_ values were calculated using GraphPad Prism 10 (La Jolla, CA, USA) using a dose-response inhibition equation with non-linear fit regression model.

### Pancoronavirus inhibiton

*In vivo* antiviral screening (pan-coronavirus assays) was performed via NIAID’s preclinical services (SRF No. 2021-1229-003). The general procedure for testing compounds is as follows:

Reduction of virus-induced cytopathic effect (Primary CPE assay) Confluent or near-confluent cell culture monolayers of Vero 76 cells (or another appropriate cell line) are prepared in 96-well disposable microplates the day before testing. Cells are maintained in MEM supplemented with 5% FBS. For antiviral assays, the same medium is used but with FBS reduced to 2% and supplemented with 50-µg/ml gentamicin. Compounds are dissolved in DMSO. The test compound is prepared at eight serial half-log10 concentrations, usually 32, 10, 3.2, 1.0, 0.32, 0.1, 0.032 and 0.01 µM. Five microwells are used per dilution: three for infected cultures and two for uninfected toxicity cultures. Controls for the experiment consist of six microwells that are infected and not treated (virus controls) and six that are untreated and uninfected (cell controls) on every plate. A known active drug is tested in parallel as a positive control drug using the same method as is applied for test compounds. On the testing day, the growth media is removed from the cells and the test compound is applied in 0.1 ml volume to wells at 2X concentration. Virus, normally at a titer that will cause >80% CPE (usually an MOI 80% CPE for most virus strains) is observed in virus control wells. The plates are then stained with 0.011% neutral red for approximately two hours at 37 °C in a 5% CO_2_ incubator. The neutral red medium is removed by complete aspiration, and the cells may be rinsed 1X with phosphate-buffered solution (PBS) to remove the residual dye. The PBS is completely removed, and the incorporated neutral red is eluted with 50% Sorensen’s citrate buffer/50% ethanol for at least 30 minutes. Neutral red dye penetrates into living cells, thus, the more intense the red color, the larger the number of viable cells present in the wells. The dye content in each well is quantified using a spectrophotometer at 540 nm wavelength. The dye content in each set of wells is converted to a percentage of dye present in untreated control wells using a Microsoft Excel-based spreadsheet and normalized based on the virus control. The 50% effective (EC_50_, virus-inhibitory) concentrations and 50% cytotoxic (CC_50_, cell-inhibitory) concentrations are then calculated by regression analysis. The quotient of CC_50_ divided by EC_50_ gives the 50% selectivity index (SI_50_) value. Compounds showing EC50 5 are considered minimally active. Reduction of virus yield (VYR assay) Active compounds are further tested in a confirmatory VYR assay. This assay is run for compounds that have an EC50 < 10 µM and SI50 ≥ 5. After sufficient virus replication occurs (generally 3 days for many viruses), a sample of supernatant is taken from each infected well (replicate wells are pooled) and held frozen at -80 °C for later virus titer determination. After maximum CPE is observed, the viable plates are stained with neutral red dye. The incorporated dye content is quantified as described above to generate the EC_50_ and CC_50_ values. The VYR test directly determines how much test compound is required to inhibit 90% virus replication. The virus yielded in the presence of the test compound is titrated and compared to virus titers from the untreated virus controls. The viral samples (collected as described in the paragraph above) are titrated by the endpoint dilution. Serial 10-fold dilutions of supernatant are made and plated into 4 replicate wells containing fresh cell monolayers of Vero 76 cells. Plates are then incubated, and cells are scored for the presence or absence of the virus after distinct CPE is observed, and the CCID_50_ is calculated using the Reed-Muench method. The 90% (one log10) effective concentration (EC_90_) is calculated by regression analysis by plotting the log10 of the inhibitor concentration versus log10 of the virus produced at each concentration. Dividing EC_90_ by the CC_50_ gives the SI_90_ value for this test.(*21*) The positive control compound for SARS-CoV-1 and MERS-CoV-1 infection was Remdesivir. The positive control for alpha 229E CoV, Beta OC43 and SARS-CoV-2 strains was EIDD-1931.

### Peptidase Selectively panel

Peptidase selectively was tested using NIH PCS services contract No: HHSN272201800007I/75N93022F00001. Eurofins completed this analysis under the NIH contract. Compounds were screened against a panel of ∼30 mammalian serine and cysteine peptidases. First a single concentration at 10 µM inhibition screen and IC_50_ was determined in follow-up for assays where the compound displayed >50% inhibition at the 10 µM. This data was then visualized in GraphPad Prism 10 and all negative values were set to 0.

### Secondary Pharmacology Screening

Secondary Pharmacology Screening selectively was tested using NIH PCS services contract No: HHSN272201800007I/75N93022F00001.

In vitro assays against a panel of ∼50 mammalian receptors and enzymes to assess potential off-target pharmacology that might lead to toxicity(*22*). Eurofins completed this analysis under the NIH contract. Test compound initially measured at a single concentration of 10 µM to determine % Inhibition relative to controls. Follow-up IC50 analysis was done where the compound exhibited >50% inhibition using 5 concentrations of test compound to enable determination of an IC_50_. This data was then visualized in GraphPad Prism 10 and all negative values were set to 0.

### Crystallography

Apo crystals of SARS-CoV-2 M^Pro^ wild type and mutants were obtained via vapor diffusion in sitting drops using Swiss 24 well plates using a concentration of 8 mg/mL mixed with the well solution containing 20 to 24% w/v polyethylene glycol 8000 and 100 mM Tris pH 7.4. Plates were incubated at 20° C and crystals grew in 3-4 days. For crystals with compounds, proteins were incubated for 1 h with 10-fold IC_50_ of the compound and trays were prepared in the same conditions as the APO crystals. For some compounds, several rounds of seeding were required to obtain good diffracting crystals. An initial seeding with crystal from the APO protein was performed to obtain small crystals for each compound, then each small crystals were harvested to obtained seeds for each compound for the second seeding. Crystals were soaked with cryoprotectant containing the well buffer, 20% glycerol, and 100 µM of compound before being flash frozen in liquid nitrogen. X-ray diffraction data were collected at the beamline 8.3.1 at the Advanced Light Source or beamline 12-1 at the Stanford Synchroton Radiation Lightsource. The data were indexed, integrated and scaled with XDS. The structure determination and refinement was performed with Phenix. Structures were first modeled and refined (with phenix.refine) without ligands to generate a difference density for the ligand. Ligand restraints were generated from SMILES using phenix.elbow and ligands were placed and refined with the rest of the protein in phenix.refine. For covalent ligands, once binding pose was identified as above, a new restraints file was generated describing covalently linked compound to the cys side chain which was then linked through “LINK” command to the c-alpha carbon. This allowed specification of the proper geometry for the covalent link to the sp2 carbon. These were refined as a mixture of: covalently bound ligand, non-covalently bound ligand and an oxidized cys with a mixed occupancy. The waters were automatically added at the end of the refinement and then manually examined. Statistics for the refined structures are reported in the Supplementary Table 10. The crystallography datasets have been deposited in the Protein Data Bank under the deposition XXX.

*Kinetic solubility, Microsomal stability, MDCK-MDR1 Bi-directional transport assay, Plasma protein binding assay and CYP450 inhibition studies were conducted at Quintara Discovery, Hayward (California, US)*.

### Kinetic Solubility

Kinetic solubility of drug substances in various buffer systems can be determined using samples supplied in DMSO solution. A sample dissolved in DMSO (typically 10 mM) is diluted with the appropriate amount of buffer (typically PBS, pH 7.4) and mixed by shaking for 1.5 hours followed by vacuum filtration. The sample is then assayed via reverse phase HPLC with UV detection. Quantitation is achieved by the reference to a three-point standard curve constructed via serial dilution of drug substance dissolved in 100% DMSO. Reference compounds (such as testosterone) are included in each test. Each compound was sent as a DMSO stock from. DMSO stocks and control compounds (such as testosterone) are thawed. Add 190 μL of buffer solution (PBS, pH 7.4 as the default buffer) to all wells on a 96-well Millipore Solubility filter plate. Transfer 10 μL of compound DMSO stocks in triplicate to the buffer wells to a final concentration of 500 μM. The filter plate is shaken for 1.5 hours at room temperature. Samples are filtered via a vacuum system into a fresh 96-well plate. Dilute compounds to 500 μM (highest concentration) in DMSO and further dilute them 1:10 for calibration curve (three-point). HPLC/UV analysis (220 nm, 254 nm, and 280 nm).

### Microsomal Stability Assay

Metabolic stability of testing compound can be evaluated using human, rat, mouse, or other animal liver or intestine microsomes to predict intrinsic clearance. The assay is carried out in 96-well microtiter plates at 37°C. Reaction mixtures (25 μL) contain a final concentration of 1 μM test compound, 0.5 mg/mL liver microsomes protein, and 1 mM NADPH and/or 1 mM UDPGA (with alamethicin) in 100 mM potassium phosphate, pH 7.4 buffer with 3 mM MgCl_2_. The incubation is done with N=2. At each of the time points example, 0, 15, 30, and 60 minutes, 150 μL of quench solution (100% acetonitrile with 0.1% formic acid) with internal standard is transferred to each well. Besides the zero-minute controls, mixtures containing the same components except the NADPH can also be prepared as the negative control. Verapamil is included as a positive control to verify assay performance. Plates are sealed, vortexed, and centrifuged at 4°C for 15 minutes at 4000 rpm. The supernatant is transferred to fresh plates for LC/MS/MS analysis. All samples are analyzed on LC/MS/MS using an AB Sciex API 4000 instrument, coupled to a Shimadzu LC-20AD LC Pump system. Analytical samples are separated using a Waters Atlantis T3 dC18 reverse phase HPLC column (20 mm x 2.1 mm) at a flow rate of 0.5 mL/min. The mobile phase consists of 0.1% formic acid in water (solvent A) and 0.1% formic acid in 100% acetonitrile (solvent B). The extent of metabolism esd calculated as the disappearance of the test compound, compared to the 0-min time incubation. Initial rates are calculated for the compound concentration and used to determine t_1/2_ values and subsequently, the intrinsic clearance, CL_int_ = (0.693)(1/t_1/2_ (min))(g of liver/kg of body weight)(mL incubation/mg of microsomal protein)(45mg of microsomal protein/g of liver weight).

### MDCK-MDR1 Bi-Directional Transport Assay

MDCK-MDR1 cells are plated into 96-well Millipore Millicell-96 plates at 7,500 cells/75 μL/well and incubated for three days at 37°C with 5% CO_2_. Cells are washed with Hank’s Balanced Salt Solution (HBSS) with 5mM HEPES for 30 minutes before starting the experiment. Test compound solutions are prepared by diluting DMSO stock into HBSS buffer, resulting in a final DMSO concentration of 0.1%. Prior to the experiment, cell monolayer integrity is verified by transendothelial electrical resistance (TEER). Transport experiment is initiated by adding test compounds to the apical (75 μL) or basal (250 μL) side.

Transport plates are incubated at 37°C in a humidified incubator with 5% CO_2_. Samples are taken from the donor and acceptor compartments after one hour and analyzed by liquid chromatography with tandem mass spectrometry (LC/MS/MS).

Digoxin is typically used as reference control. Apparent permeability (Papp) values are calculated using the following equation:

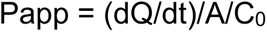

where dQ/dt is the initial rate of amount of test compound transported across cell monolayer, A is the surface area of the filter membrane, and C_0_ is the initial concentration of the test compound, calculated for each direction using a 4-point calibration curve by LC/MS/MS.

Net flux ratio between the two directional transports is calculated by the following equation:

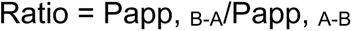

where Papp, _B-A_ and Papp, _A-B_ represent the apparent permeability of test compound from the basal-to-apical and apical-to-basal side of the cellular monolayer, respectively.

Recovery is calculated based on the compound concentration at the end of the experiment, compared to that at the beginning of the experiment, adjusted for volumes. A net flux ratio greater than two is considered a positive result for substrate determination.

### Plasma Protein Binding Assay

The rapid equilibrium dialysis (RED) device inserts along with a Teflon base plate (Pierce, Rockford, IL) are used for the binding studies. Human or animal plasma is obtained commercially. The pH of the plasma is adjusted to 7.4 prior to the experiment.

DMSO stocks (1 mM) are spiked into the plasma to make a final concentration of 2 μM. Aliquots of (100 μL) were transferred to a fresh 96-well deep-well plate as the T4 (recovery) samples. An equal volume of blank PBS buffer is added to the plate to make the matrix as 50:50 plasma:buffer. The T4 recovery samples are incubated at 37°C for 4 hours. The spiked plasma solutions (300 μL) were placed into the sample chamber (indicated by the red ring); and 500 μL of PBS buffer, pH 7.4, is placed into the adjacent chamber. The plate is sealed with a self-adhesive lid and incubated at 37°C on an orbital shaker (250 rpm) for 4 hours. After 4 hours, from the RED plate, aliquots (100 μL) are removed from each side of the insert (plasma and buffer) and dispensed into the 96-well plate. Subsequently, 100 μL of blank plasma is added to the buffer samples and 100 μL of blank buffer is added to all the collected plasma samples. At last, 300 μL of quench solution (50% acetonitrile, 50% methanol, and 0.05% formic acid, warmed up at 37°C) containing internal standards is added to each well. Plates are sealed, vortexed, and centrifuged at 4°C for 15 minutes at 4000 rpm. The supernatant is transferred to fresh plates for LC/MS/MS analysis. Reference compound propranolol was included in every experiment. All samples were analyzed on LC/MS/MS using an AB Sciex API 4000 instrument, coupled to a Shimadzu LC-20AD LC Pump system. Analytical samples are separated using a Waters Atlantis T3 dC18 reverse phase HPLC column (20 mm x 2.1 mm) at a flow rate of 0.5 mL/min. The mobile phase consisted of 0.1% formic acid in water (solvent A) and 0.1% formic acid in acetonitrile (solvent B).

The percentage of test compound bound to protein is calculated by the following equation:

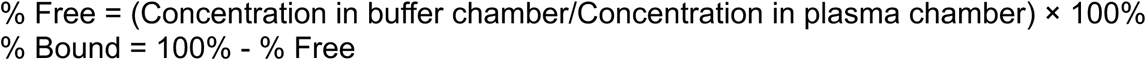

the percentage of test compound recovered was calculated by the following equation:

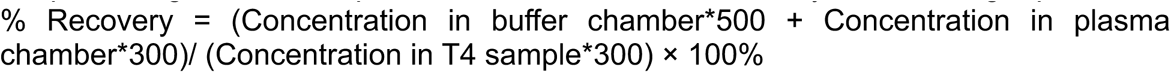

All the samples are diluted by quench solution to around 400 nM to be within compounds’ linear ranges.

### CYP450 Inhibition Assay

Selective substrates are incubated with pooled human liver microsomes as single substrates. The assays were performed in 384-well plates using a final volume of 40 μL at 37°C. All assays employ 100 mM potassium phosphate buffer, pH 7.4, with 3 mM MgCl2 and 1 mM cofactor NADPH. Compounds were tested at 10 µM to obtain % inhibition. Human cytochrome specific inhibitors are also included within each assay as reference compounds. The quantitation window can be defined as 100% enzyme activity (NADPH added) in the presence of vehicle control (DMSO). Analysis was via LC/MS/MS where enzyme activity is based on the detection of appearance of the respective substrate metabolites.

Briefly the conditions for all tested CYP450 are as follows: For CYP1A2 to 0.1 mg/mL of human liver microsome (HLM) the substrate phenacetin was used at 30 µM and the metabolite acetaminophen was monitored after 10 min. For CYP2B6, to 0.1 mg/mL of HLM, the substrate bupropion was used at 100 uM and the metabolite hydroxybupropin was monitored after 10 min. For CYP2C8 to 0.1 mg/mL of HLM the substrate paclitaxel was used at 2 µM and the metabolite 6α-Hydroxypaclitaxel was monitored after 10 min. For CYP2C9, to 0.1 mg/mL of HLM, the substrate diclofenac was used at 4 µM and the metabolite hydroxydiclofenac was monitored after 10 min. For CYP2C19, to 0.2 mg/mL of HLM, the substrate mephenytoin was used at 35 µM and the metabolite 4’-hydroxymephenytoin was monitored after 20 min. For CYP2D6, to 0.1 mg/mL of HLM, the substrate bufuralol was used at 10 µM and the metabolite hydroxybufuralol was monitored after 10 min. For CYP3A4, to 0.05 mg/mL of HLM, either testosterone (30 µM) or midazolam (5 µM) was added and the metabolites 6β-hydroxytestosterone or 1’-hydroxymidazolam were monitored respectively, after 5 min.

All samples were analyzed on LC/MS/MS using an AB Sciex API 4000 instrument, coupled to a Shimadzu LC-20AD LC Pump system. Analytical sample of 1A2-ACE is separated using a Thermo Hypersil Gold C18 (50 x 2.1 mm) column, and other samples were separated using a Waters Atlantis T3 dC18 reverse phase HPLC column (20 mm x 2.1 mm) at a flow rate of 0.5 mL/min. The mobile phase consists of 0.1% formic acid in water (solvent A) and 0.1% formic acid in acetonitrile (solvent B).

### Thermal proteome profiling (TPP) assay

For TPP optimization experiments, 5 µM MPro was treated with either DMSO or compound at a final concentration of 0.5, 5, 50, or 100 µM at room temperature for 30 minutes. Each condition was split into 20 µL aliquots, and aliquots from each condition were heated for 4 minutes on a BioRad C1000 Touch Thermal cycler at the following temperatures: 37, 39, 42.3, 46.4, 51.9, 56.1, 59, 61°C. Samples were centrifuged at 20,000xG for 60 minutes. Supernatant was incubated with 8M urea, 100 mM tris, 10 mM TCEP/44 mM CAA (pH ∼ 7.5) for 60 minutes. The urea concentration was diluted to 1 M with 100 mM tris (pH ∼7.5). Samples were digested overnight with 1 µL trypsin (Promega, 0.4 µg/µL). Samples were desalted with a 96-well mini 20MG PROTO 300 C18 plate (HNS S18V, The Nest Group) according to manufacturer’s directions. Peptide concentration was determined by NanoDrop (Thermo).

For optimization experiments, peptides were injected onto an Orbitrap Exploris 480 MS system (Thermo) equipped with an Easy nLC 1200 system (Thermo). Peptides were separated on a PepSep reverse-phase C18 column (1.9 mm particles, 1.5 mm x 15 cm, 150 mm ID) (Bruker). Mobile phase A consisted of 0.1% FA and mobile phase B consisted of 80% acetonitrile (ACN)/0.1% FA. Peptide mixtures were separated by mobile phase B ranging from 0% to 28% over 27 minutes, followed by an increase to 45% B over 4 minutes, then held at 95% B for 9 minutes at a flow rate of 500 nL/min. Samples were analyzed by DDA with an MS1 resolution of 120K (@200 m/z), a scan range of 350-1250 m/z, an MS1 normalized AGC target of 300%, and an exclusion duration of 30 s. MS2 cycle time was set to 1 s, with an isolation window of 1.3 m/z. Samples were fragmented at 28% HCD (higher-energy collisional dissociation) in Auto Scan Range Mode using an AGC target of 200%. MS2 Orbitrap resolution was set to 15,000.

For lysate experiments, pelleted A549 cells were resuspended in extraction buffer (1x PBS + phosphatase and protease inhibitors (phosSTOP (Roche) and cOmplete Mini Protease Inhibitor Cocktail (Roche)) with gentle pipetting followed by rotation at 4°C for 30 minutes. Lysates were centrifuged at 1000xG for 10 minutes at 4°C and supernatant was transferred to new tubes. Lysates (2 replicates per condition) were distributed into 10 20 uL aliquots in PCR tubes. Samples were heated from 37 to 64 in 3°C increments on a BioRad C1000 Touch Thermal cycler and held for four minutes at the specified temperature. Samples were held at room temperature for three minutes. Samples were flash frozen, followed by thawing at 35°C (x2). Aggregated proteins were removed by centrifugation at 20,000 x G for 60 mins. 20 µL of lysis buffer (8 M urea, 100 mM tris, pH ∼7.5) was added to each well and samples were incubated for 30 minutes at room temperature. Samples were reduced and alkylated by the addition of TCEP (100mM final) and 2-chloroacetamide (44mM final) followed by incubation at room temperature for 30 mins. Urea concentration was diluted to 1 M with 100 mM tris (pH ∼7.5). Samples were digested overnight with LysC (Wako, 1:100 enzyme: protein ratio) and trypsin (Promega, 1:50 enzyme:protein ratio). Samples were desalted with a 96-well mini 20MG PROTO 300 C18 plate (HNS S18V, The Nest Group) according to manufacturer’s directions. Peptide concentration was determined by NanoDrop (Thermo).

For lysate experiments, equal amounts of peptides were injected onto a timsTOF SCP (Bruker) connected to a EASY-nLC 1200 system (Thermo). Peptides were separated on a PepSep reverse-phase C18 column (1.9 mm particles, 1.5 mm x 15 cm, 150 mm ID) (Bruker) with a gradient of 5-28% buffer B (0.1% formic acid in acetonitrile) over buffer A (0.1% formic acid in water) over 20 minutes, an increase to 32% B in 3 minutes, and held at 95% B for 7 minutes. DIA-PASEF analyses were acquired from 100 to 1700 m/z over a 1/Kø of 0.70 to 1.30 Vs/cm^2^, with a ramp and accumulation time set to 75 ms. Library DDA PASEF runs were collected over the same m/z and 1/Kø range and a cycle time of 1.9 s.

All data was searched against the Uniprot Human database (downloaded 05/25/23) appended with the SARS-CoV-2 database (downloaded 02/20/2024) using a combined DDA and DIA library in Spectronaut (Biognosys, version 16.0). Default settings, including trypsin digestion, variable modifications of methionine oxidation and N-termini acetylation, and fixed modification of cysteine carbamidomethylation, were used. Missing values were imputed for each run using background intensity. Data was filtered to obtain a false discovery rate of 1% at the peptide spectrum match and protein level. Lysate experiments were normalized(*23*) and melting points were determined in R using the Inflect package(*24*).

### Mice

All animal use protocols (AN203103-00A) were approved by the Institutional Animal Care and Use Committees at the University of California, San Francisco, and Gladstone Institutes, and were conducted in strict accordance with the National Institutes of Health Guide for the Care and Use of Laboratory Animals (National Research Council (US) Committee for the Update of the Guide for the Care and Use of Laboratory Animals, 2011). The studies involved 6–8 week old female wild-type mice (The Jackson Laboratory, 000664). The mice were housed in a pathogen-free facility with controlled temperature and humidity, a 12-hour light/dark cycle, and *ad libitum* access to water and standard laboratory rodent chow.

### SARS-CoV-2 culture for mice studies

The SARS-CoV-2 Beta variant was used for all the mice infection studies. All live virus experiments were performed in a Biosafety Level 3 laboratory. SARS-CoV-2 stocks were propagated in Vero-ACE2-TMPRSS2 cells, and their sequence verified by next-generation sequencing. Viral stock titer was calculated using plaque forming assays.

#### Antiviral screening of compounds in wild type mice

Forty-five wild-type mice were infected with the SARS-CoV-2 Beta variant at a dose of 10³ PFU and divided into three treatment groups: AVI-4206 (100 mg/kg), vehicle, and Nirmatrelvir (300 mg/kg) as a positive control, with each group containing 15 mice. Treatment commenced 4 hours post-infection with oral BID dosing for 5 days, during which the animals were closely monitored for disease parameters such as weight loss, hypothermia, and posture. At 2, 4, and 7 days post-infection, five animals from each group were euthanized, and their lung tissue was harvested and homogenized for downstream analysis using plaque assay. The left lung lobe tissue from an additional subset of animals were processed for histological observations.

### Evaluating dose-dependent efficacy of AVI-4516

The dose-dependent antiviral efficacy of AVI-4516 was evaluated in SARS-CoV-2 Beta-infected WT mice. A group of 25 wild-type female mice aged 6-8 weeks were infected with the SARS-CoV-2 Beta variant at 103 PFUs. The mice were orally dosed with a range of concentrations of AVI-4516 starting from 12.5 to 100mg/kg. The treatment was started at 4 hours post-infection followed by BID on day 1 post-infection. All the mice were euthanized at day 2 post-infection and their lung tissues were harvested to estimate the virus titers in the lungs.

### Plaque assays

The lung homogenates were clarified by centrifugation, and the supernatants were serially diluted to infect Vero ACE2-TMPRSS2 cells. Following a one-hour absorption period, 2.5% Avicel (Dupont, RC-591) was applied to the cells and incubated for 48 hours. After incubation, the Avicel was removed, and the cells were fixed in 10% formalin for one hour, and then stained with crystal violet for 10 minutes. Plaques were counted, and the data were presented as plaque-forming units.

## Chemical Synthesis

### General Experimental Procedures

Unless otherwise noted all chemical reagents and solvents used are commercially available. Air and/or moisture sensitive reactions were carried out under an argon atmosphere in oven-dried glassware using anhydrous solvents from commercial suppliers. Air and/or moisture sensitive reagents were transferred via syringe or cannula and were introduced into reaction vessels through rubber septa. Solvent removal was accomplished with a rotary evaporator at ca. 10-50 Torr. NMR spectra were recorded on a Bruker Avance III HD 400 MHz spectrometer. Chemical shifts are reported in δ units (ppm). NMR spectra were referenced relative to residual NMR solvent peaks. Coupling constants (*J*) are reported in hertz (Hz). Chromatography was carried out using Isolera Four and CombiFlash NextGen 300 flash chromatography systems with Silia*Sep* silica gel and C18 cartridges from Silicycle. Reverse phase chromatography was carried out on Waters 2535 Separation module with Waters 2998 Photodiode Array Detector. Separations were carried out on XBridge Preparative C18, 19 x 50 mm column at ambient temperature using a mobile phase of water-acetonitrile containing a constant 0.1% formic acid. LC/MS data were acquired on a Waters Acquity UPLC QDa mass spectrometer equipped with Quaternary Solvent Manager, Photodiode Array Detector and Evaporative Light Scattering Detector. Separations were carried out with Acquity UPLC® BEH C18 1.7 μm, 2.1 x 50 mm column at 25°C, using a mobile phase of water-acetonitrile containing a constant 0.1 % formic acid.

### General Procedure for the Synthesis of Trisubstituted Uracil Derivatives

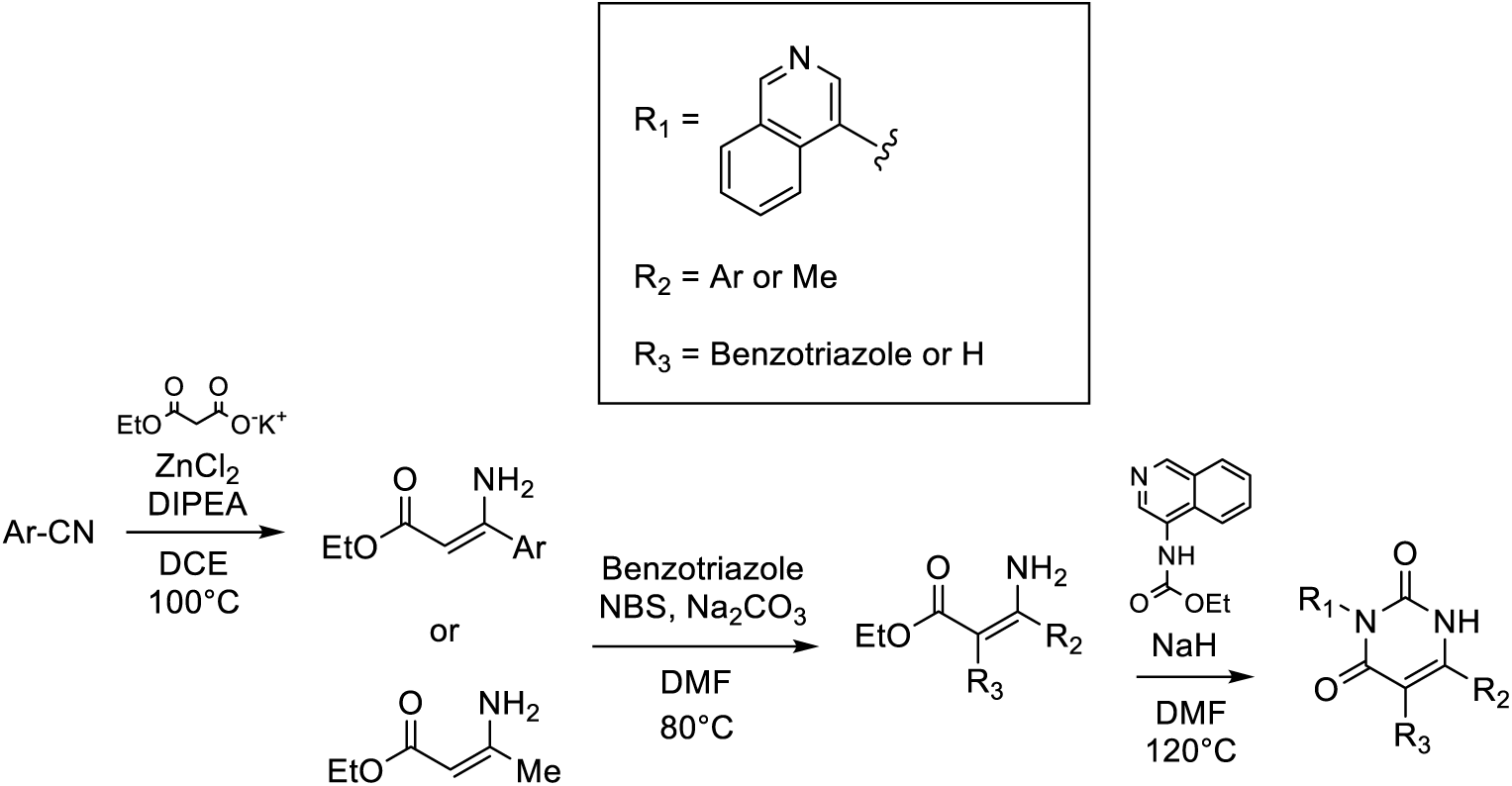

### General Procedure for the Synthesis of Tetrasubstituted Uracil Derivatives

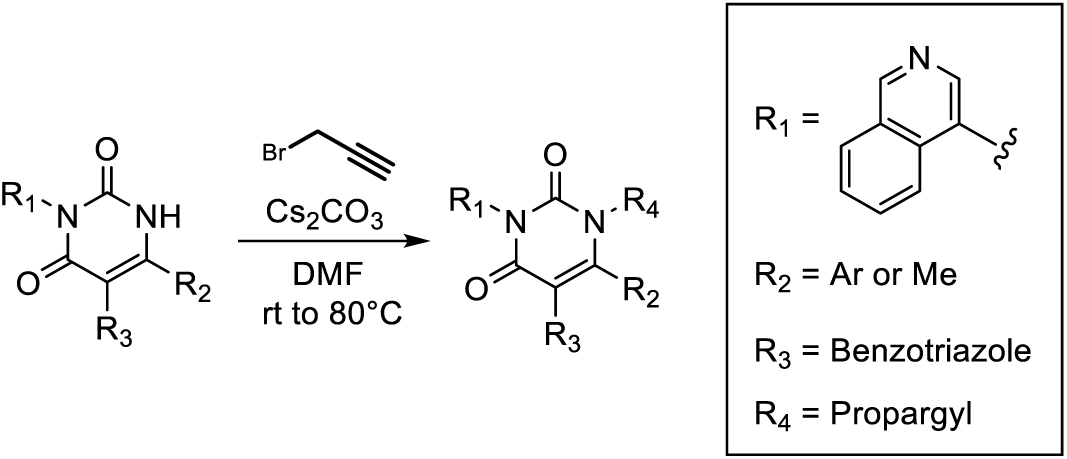

### Synthesis of carbamic acid-4-isoquinolinyl-ethyl ester

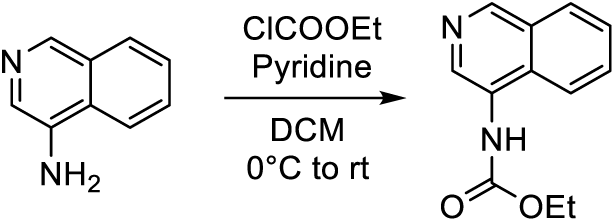

Pyridine (1.7 mL, 20.8 mmol, 3 eq) was added to a suspension of 4-aminoisoquinoline (1 g, 6.94 mmol) in DCM (25 mL) at 0°C, followed by a dropwise addition of ethyl chloroformate (0.995 mL, 10.41 mmol, 1.5 eq) dissolved in 5 mL of DCM. Then the mixture was warmed to room temperature and stirred for 1 h and quenched with 1N HCl (20 mL). The aqueous phase was extracted with DCM (3×20 mL). The organic phase was dried over Na_2_SO_4_ and concentrated in vacuo. The crude material (1 g, 6.02 mmol, 87%) was used in the next step without further purification.

**General:** C_12_H_12_N_2_O_2_; MW = 216.24.

**LCMS (ESI):** *m/z* = 217.1 [M+H]^+^.

#### General Procedure A: Blaise Reaction of Aryl Nitriles

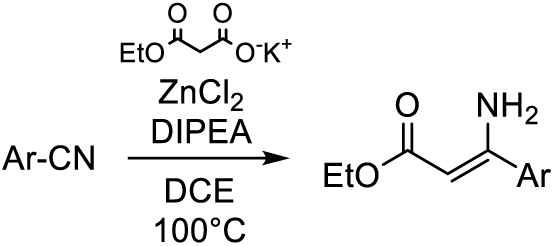

To a solution of the aryl nitrile (1 mmol) in 1,2-dichloroethane (10 mL), were added ZnCl_2_ (1.2 eq), potassium ethyl malonate (2.3 eq) and DIPEA (0.3 eq). The mixture was stirred at 100°C for 16 h, then cooled to room temperature and washed with saturated NH_4_Cl aqueous solution. The aqueous phase was extracted with DCM and the organic extracts were dried over Na_2_SO_4_ and concentrated in vacuo. The crude material was used in the next step without further purification.

#### General Procedure B: Amination of Enaminones

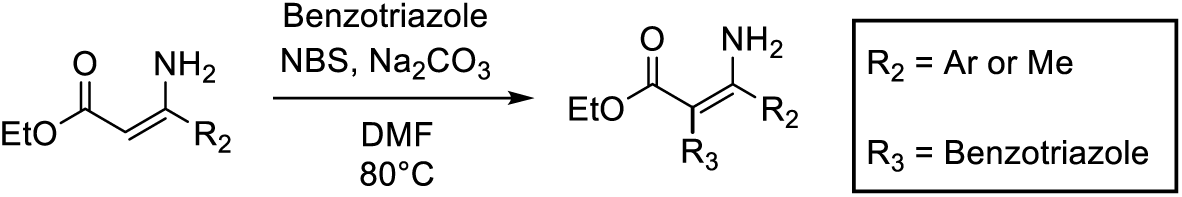

*N*-Bromosuccinimide (1.2 eq) was added to a solution of enaminone (1 mmol) in DMF (2 mL) and the mixture was stirred for 10 min at room temperature. Then, the corresponding benzotriazole (1.2 eq), and Na_2_CO_3_ (1.2 eq) were added, and the mixture was heated to 80°C and stirred for 2 h. The reaction was quenched with saturated Na_2_S_2_O_3_ aqueous solution, and the aqueous phase was extracted with EtOAc. The organic extracts were dried over Na_2_SO_4_ and concentrated in vacuo. The crude was purified by flash chromatography on silica gel (EtOAc in hexane 0% to 60%).

#### General Procedure C: Synthesis of Uracil Analogs

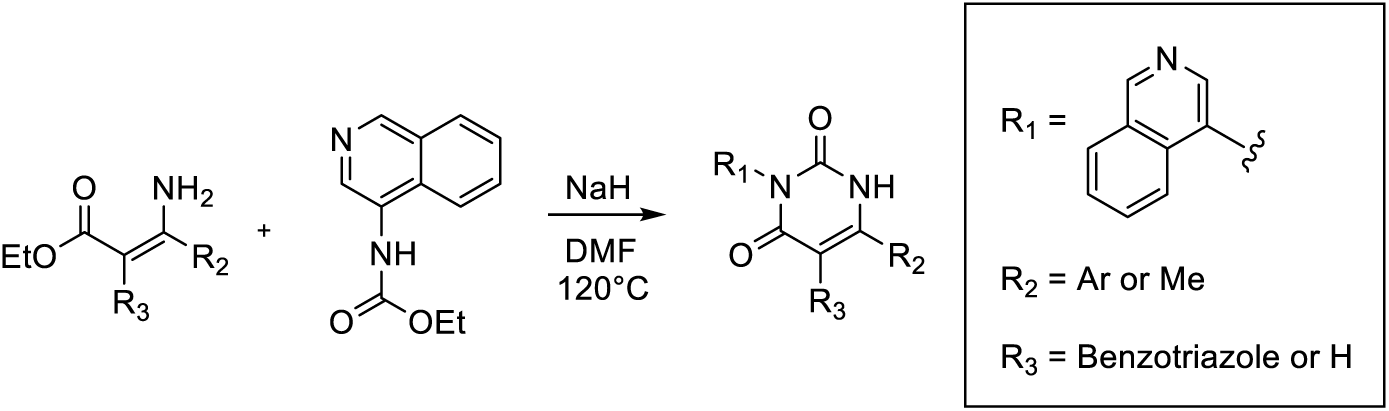

The enaminone (0.2 mmol) was dissolved in DMF (1 mL) and added dropwise to a suspension of NaH (60% in mineral oil; 2.5 eq) in DMF (0.5 mL) at 0°C. The mixture was stirred for 30 min at 0°C and then added to a solution of carbamic acid-4-isoquinolinyl-ethyl ester (1.5 eq) in DMF (1 mL) at 0°C. The mixture was heated to 120°C and stirred for 2 h; then cooled to room temperature and directly purified by preparative HPLC.

#### General Procedure D: Alkylation of Uracil

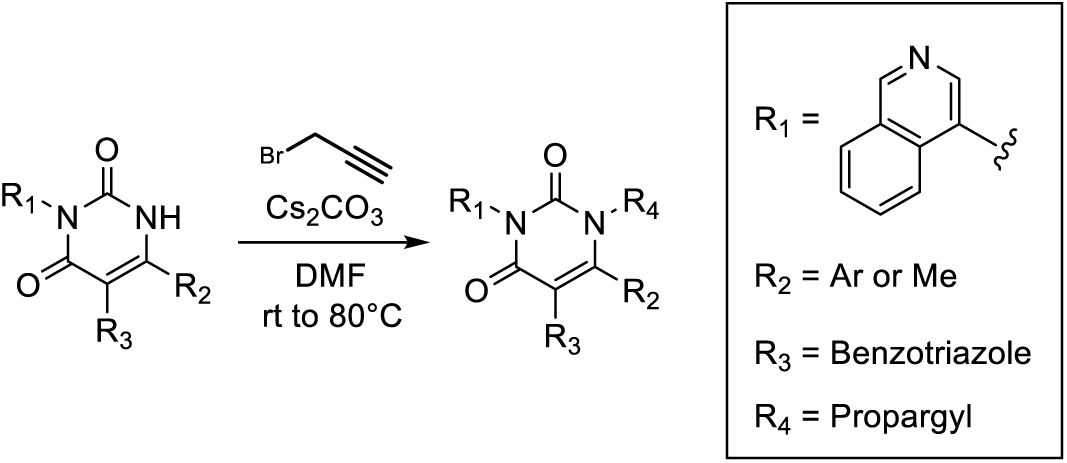

The uracil starting material (0.3 mmol) was dissolved in DMF (3 mL), then Cs_2_CO_3_ (1.5 eq) and propargyl bromide (1.2 eq) were added at room temperature. The mixture was stirred at room temperature or heated to 80°C for 4-16 h, depending on the substrate, and directly purified by preparative HPLC (40% to 90% CH_3_CN in H_2_O + 0.1% formic acid).

#### Synthesis of AVI-4301

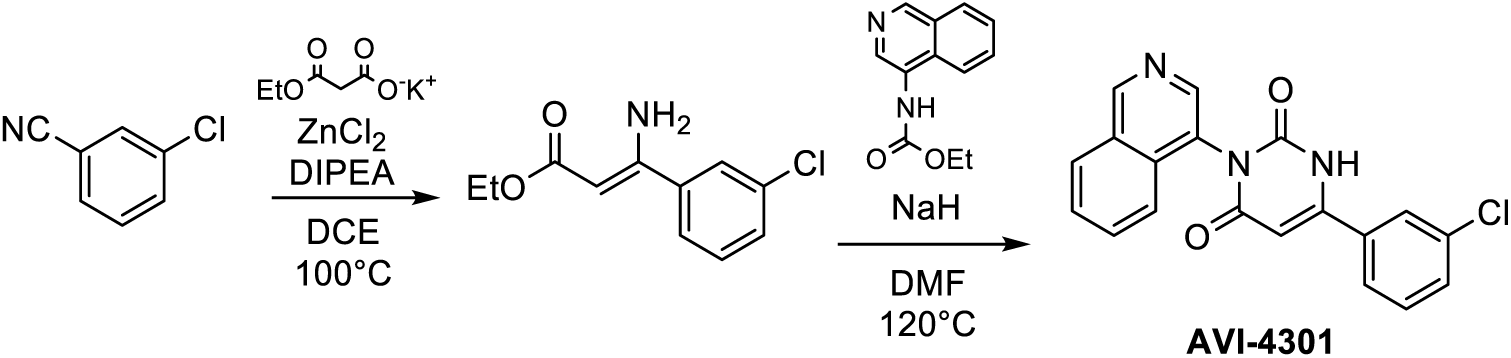

### Step 1: Ethyl 3-amino-3-(3-chlorophenyl)-2-propenoate

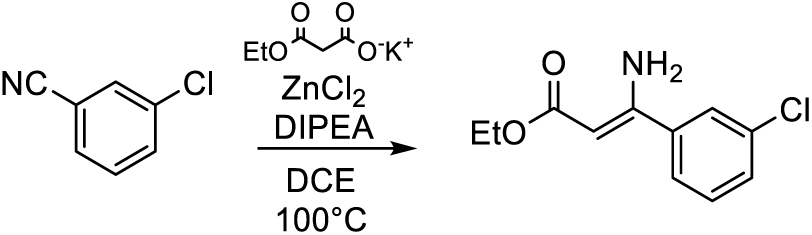

The general procedure A (Blaise Reaction) was followed, using 3-chlorobenzonitrile (1 g, 7.27 mmol).

Yield: 1.52 g, 6.75 mmol, 93%.

**General**: C_11_H_12_ClNO_2_; MW = 225.67.

**LCMS (ESI)**: *m/z* = 226.1 [M+H]^+^.

### Step 2: AVI-4301

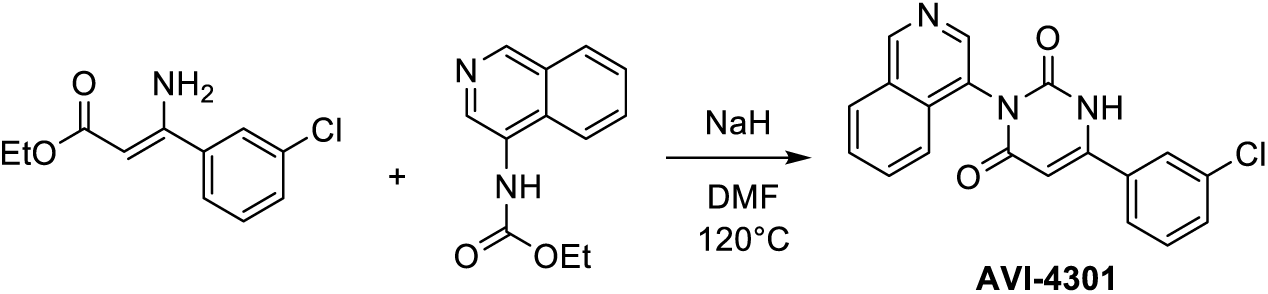

General Procedure C (Synthesis of Uracil) was followed using ethyl 3-amino-3-(3-chlorophenyl)-2-propenoate (25 mg, 0.111 mmol). Purification by preparative HPLC (20% to 70% CH_3_CN in H_2_O + 0.1% formic acid) afforded **AVI-4301** (13.5 mg, 0.0387 mmol, 35%) as a white solid.

**General**: C_19_H_12_ClN_3_O_2_; MW = 349.77.

**^1^H-NMR** (400 MHz, DMSO-d6): δ (ppm): 10.82 (brs, 1H); 9.36 (s, 1H); 8.51 (s, 1H); 8.11 (m, 1H); 7.74 (m, 1H); 7.68 (t, *J* = 5.6 Hz, 2H); 7.57 (t, *J* = 1.8 Hz, 1H); 7.42 (dd, *J* = 14.5, 8.0 Hz, 1H); 7.26 (s, 1H); 7.16 (t, *J* = 7.9 Hz, 1H); 6.17 (s, 1H).

**LCMS (ESI)**: *m/z* = 350.2 [M+H]^+^.

#### Synthesis of AVI-4303

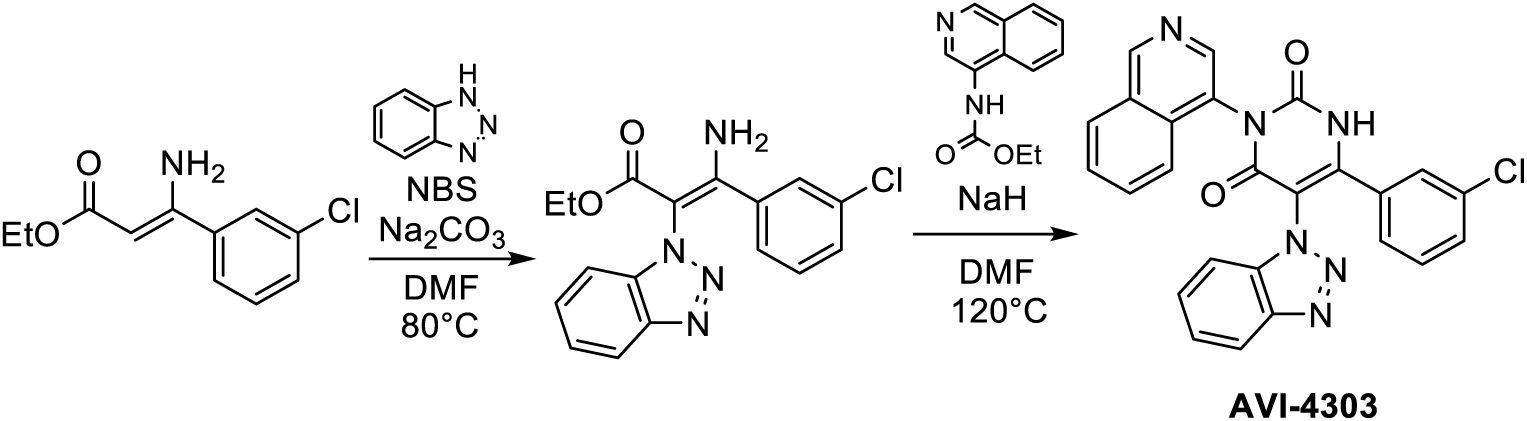

### Step 1: Ethyl 3-amino-2-(1H-benzo[d][1,2,3]triazol-1-yl)-3-(3-chlorophenyl)acrylate

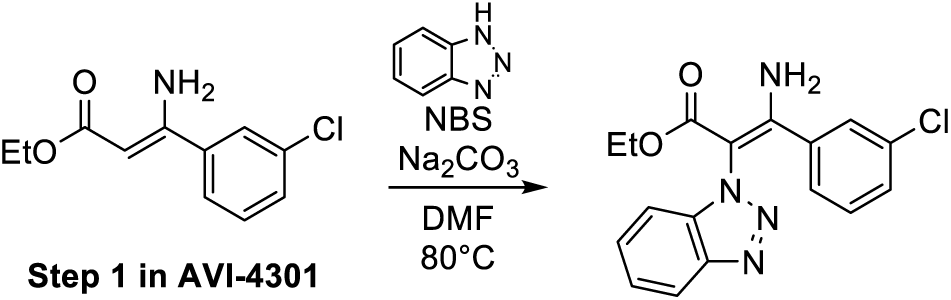

General Procedure B (Amination of Enaminone) was followed using ethyl 3-amino-3-(3-chlorophenyl)-2-propenoate (200 mg, 0.886 mmol) and 1*H*-benzotriazole. Yield: 80 mg, 0.233 mmol, 26%.

**General**: C_17_H_15_ClN_4_O_2_; MW = 342.78.

**LCMS (ESI)**: *m/z* = 343.2 [M+H]^+^.

### Step 2: AVI-4303

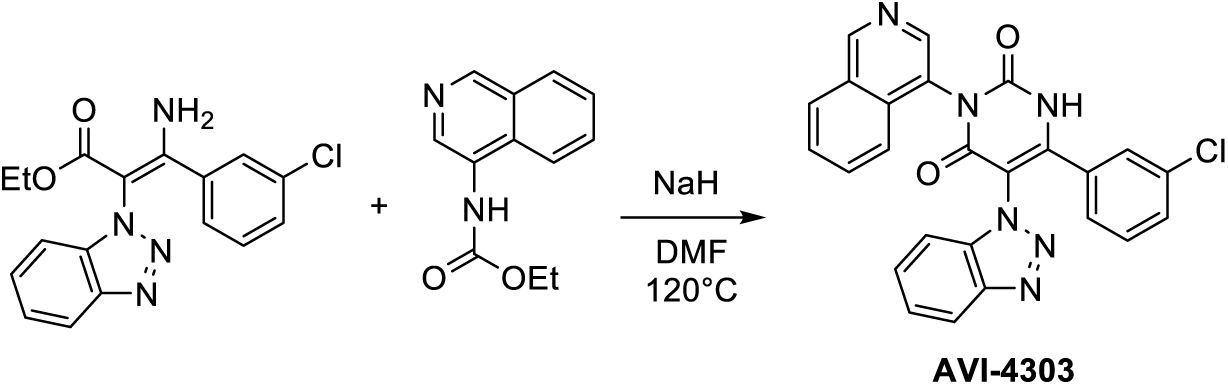

General Procedure C (Synthesis of Uracil) was followed using ethyl 3-amino-2-(1H-benzo[d][1,2,3]triazol-1-yl)-3-(3-chlorophenyl)acrylate (122 mg, 0.357 mmol). Purification by preparative HPLC (30% to 70% CH_3_CN in H_2_O + 0.1% formic acid) afforded **AVI-4303** (55 mg, 0.118 mmol, 33%) as a white solid.

**General**: C_25_H_15_ClN_6_O_2_; MW = 466.89.

**^1^H-NMR** (400 MHz, DMSO-d6): δ (ppm): 12.56 (brs, 1H); 9.44 (s, 1H); 8.65 (s, 1H); 8.28 (d, *J* = 8.3 Hz, 1H); 8.26-8.04 (m, 1H); 8.01 (d, *J* = 8.3 Hz, 1H); 7.96-7.82 (m, 2H); 7.79 (t, *J* = 7.6 Hz, 1H); 7.60-7.51 (m, 2H); 7.43 (dt, *J* = 7.7, 1.7 Hz, 1H); 7.38 (t, *J* = 7.7 Hz, 1H); 7.34-7.17 (m, 2H).

**^13^C-NMR** (101 MHz, DMSO-d6): δ (ppm): 160.3, 153.4, 152.4, 152.3, 150.5, 144.5, 143.2, 140.7, 132.9, 132.7, 131.7, 131.2, 130.9, 128.4, 128.1, 127.9, 127.1, 126.5, 126.4, 124.4, 121.9, 119.3, 110.9, 110.8, 108.1.

**LCMS (ESI)**: *m/z* = 467.1 [M+H]^+^.

#### Synthesis of AVI-4692

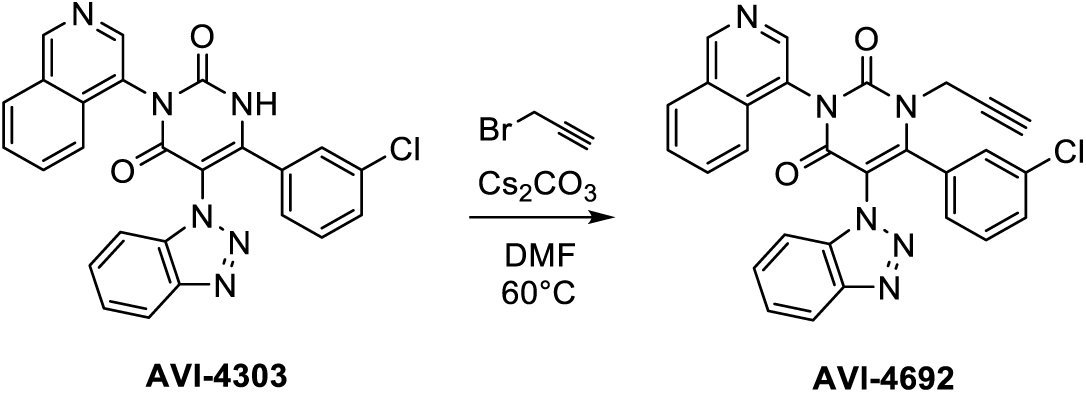

General Procedure D (Alkylation of Uracil) was followed using AVI-4303 (20 mg, 0.0428 mmol). The mixture was stirred at 60°C for 4 h. Yield: 10.5 mg, 0.0208 mmol, 49%.

**General**: C_28_H_17_ClN_6_O_2_; MW = 504.93.

**^1^H-NMR** (400 MHz, CD_3_CN): δ (ppm): 9.40 (s, 1H); 8.63 (d, *J* = 18.7 Hz, 1H); 8.22 (d, *J* = 8.3 Hz, 1H); 8.17-8.00 (m, 1H); 7.92 (q, *J* = 8.5 Hz, 2H); 7.82-7.52 (m, 4H); 7.36 (t, *J* = 7.6 Hz, 3H); 7.50-7.17 (m, 1H); 4.67-4.26 (m, 2H); 2.68 (m, 1H).

**LCMS (ESI)**: *m/z* = 505.1 [M+H]^+^.

#### Synthesis of AVI-4673

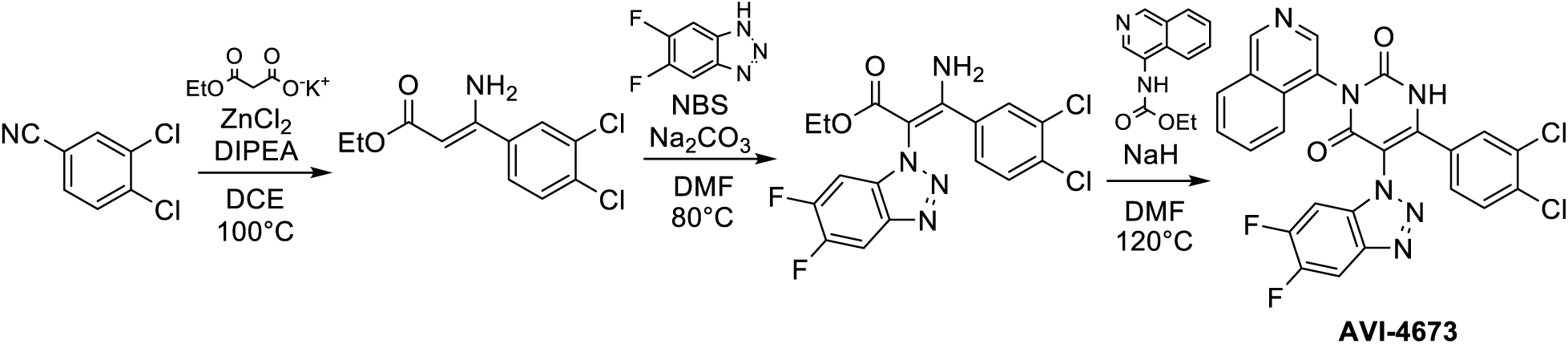

### Step 1: Ethyl 3-amino-3-(3,4-dichlorophenyl)-2-propenoate

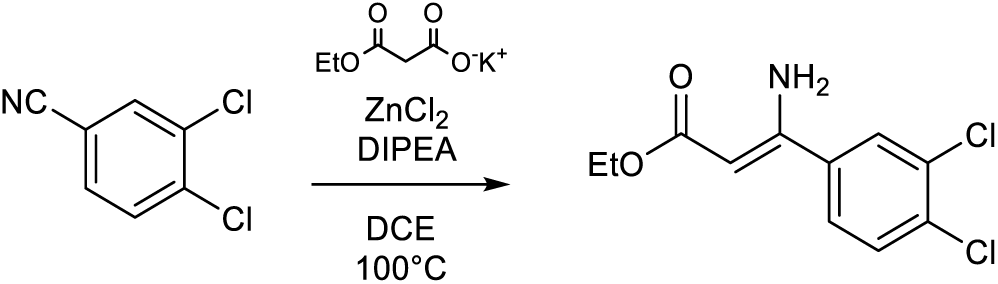

The general procedure A (Blaise Reaction) was followed, using 3,4-dichlorobenzonitrile (1 g, 5.81 mmol).

Yield: 1.42 g, 5.50 mmol, 95%.

**General**: C_11_H_11_Cl_2_NO_2_; MW = 260.11.

**LCMS (ESI)**: *m/z* = 260.1 [M+H]^+^.

### Step 2: Ethyl 3-amino-2-(5,6-difluoro-1H-benzo[d][1,2,3]triazol-1-yl)-3-(3,4-dichlorophenyl)acrylate

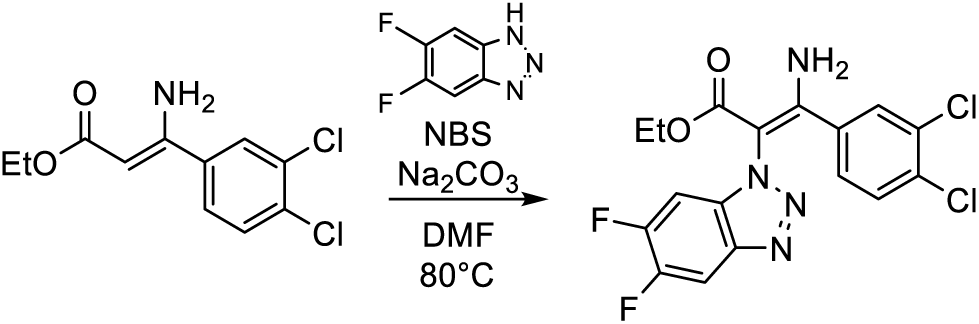

General Procedure B (Amination of Enaminone) was followed using ethyl 3-amino-3-(3,4-dichlorophenyl)-2-propenoate (80 mg, 0.309 mmol) and 5,6-difluoro-1*H*-benzotriazole. Yield: 59 mg, 0.143 mmol, 46%.

**General**: C_17_H_12_Cl_2_F_2_N_4_O_2_; MW = 413.21.

**LCMS (ESI)**: *m/z* = 413.2 [M+H]^+^.

#### Step 3: AVI-4673

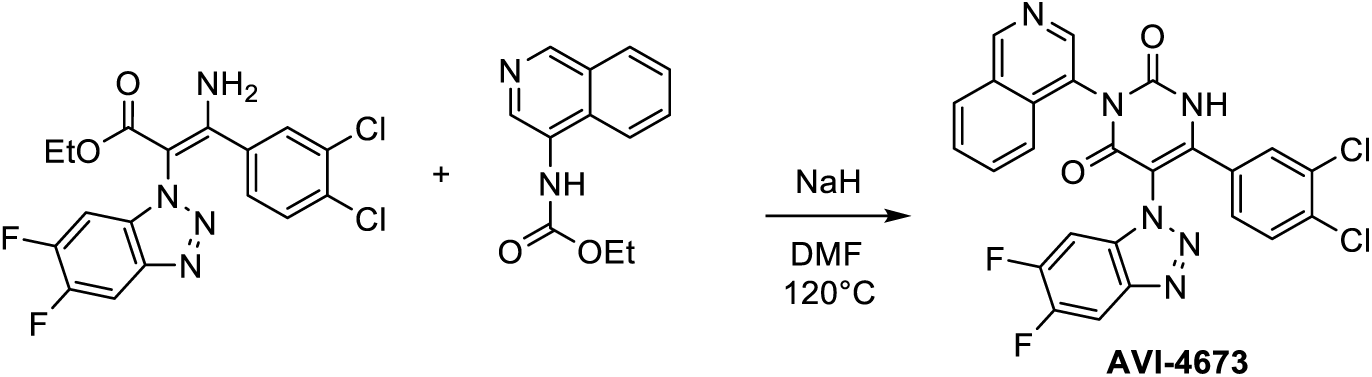

General Procedure C (Synthesis of Uracil) was followed using ethyl 3-amino-2-(5,6-difluoro-1H-benzo[d][1,2,3]triazol-1-yl)-3-(3,4-dichlorophenyl)acrylate (59 mg, 0.143 mmol). Purification by preparative HPLC (30% to 70% CH_3_CN in H_2_O + 0.1% formic acid) afforded **AVI-4673** (29 mg, 0.0540 mmol, 38%) as a white solid.

**General**: C_25_H_12_Cl_2_F_2_N_6_O_2_; MW = 537.31.

**^1^H-NMR** (400 MHz, DMSO-d6): δ (ppm): 12.68 (brs, 1H); 9.45 (s, 1H); 8.63 (s, 1H); 8.32-8.06 (m, 4H); 7.93 (brs, 1H); 7.83-7.73 (m, 2H); 7.61 (d, *J* = 8.3 Hz, 1H); 7.24 (brs, 1H).

**^13^C-NMR** (101 MHz, DMSO-d6): δ (ppm): 160.0, 153.4, 150.6, 149.9, 149.6, 147.1, 143.0, 139.7, 139.6, 133.8, 132.7, 131.7, 131.2, 131.1, 130.9, 130.2, 130.1, 128.8, 128.1, 128.0, 127.1, 121.8, 107.6, 106.8, 106.6.

**LCMS (ESI)**: *m/z* = 537.2 [M+H]^+^.

#### Synthesis of AVI-4694

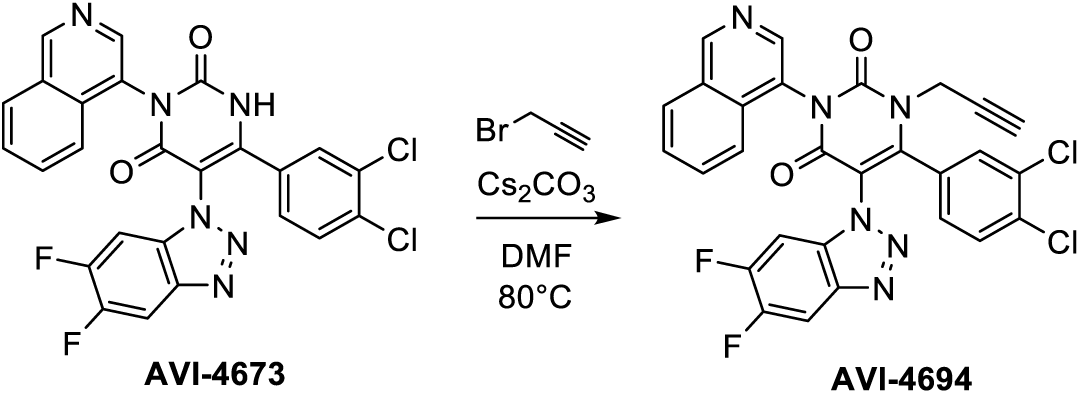

General Procedure D (Alkylation of Uracil) was followed using AVI-4673 (200 mg, 0.372 mmol). The mixture was stirred at 80°C for 4 h. Yield: 83.5 mg, 0.145 mmol, 39%.

**General**: C_28_H_14_Cl_2_F_2_N_6_O_2_; MW = 575.36.

**^1^H-NMR** (400 MHz, DMSO-d6): δ (ppm): 9.48 (s, 1H); 8.67 (m, 1H); 8.34-8.08 (m, 4H); 8.03-7.76 (m, 3H); 7.75-7.57 (m, 1H); 7.55-7.29 (m, 1H); 4.73-4.18 (m, 2H); 3.49 (m, 1H).

**^13^C-NMR** (101 MHz, DMSO-d6): δ (ppm): 158.7, 154.0, 153.7, 150.2, 150.1, 149.9, 149.7, 143.0, 142.9, 139.5, 139.4, 133.7, 132.4, 131.8, 131.1, 129.6, 128.8, 128.2, 127.1, 127.0, 121.7, 121.5, 106.7, 98.8, 98.6, 78.1, 75.9, 37.6.

**LCMS (ESI)**: *m/z* = 575.0 [M+H]^+^.

#### Synthesis of AVI-4516

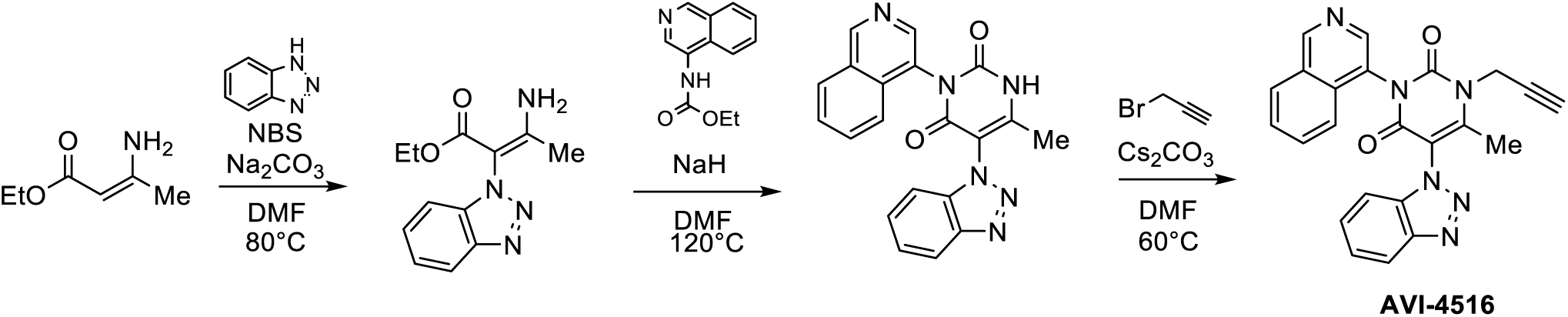

### Step 1: Ethyl 3-amino-2-(1H-benzo[d][1,2,3]triazol-1-yl)-but-2-enoate

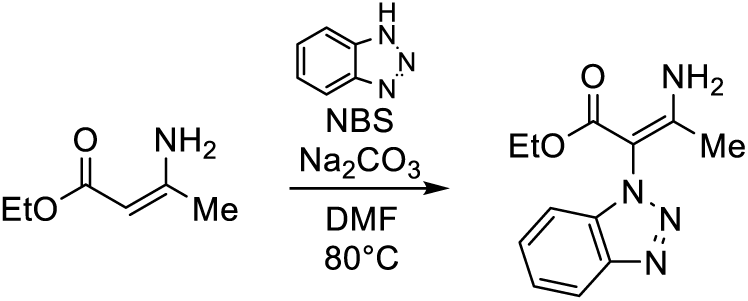

General Procedure B (Amination of Enaminone) was followed using ethyl 3-amino-but-2-enoate (50 mg, 1.56 mmol) and 1*H*-benzotriazole. Yield: 63 mg, 0.39 mmol, 66%.

**General**: C_12_H_14_N_4_O_2_; MW = 246.11.

**LCMS (ESI)**: *m/z* = 247.25 [M+H]^+^.

### Step 2: AVI-4375

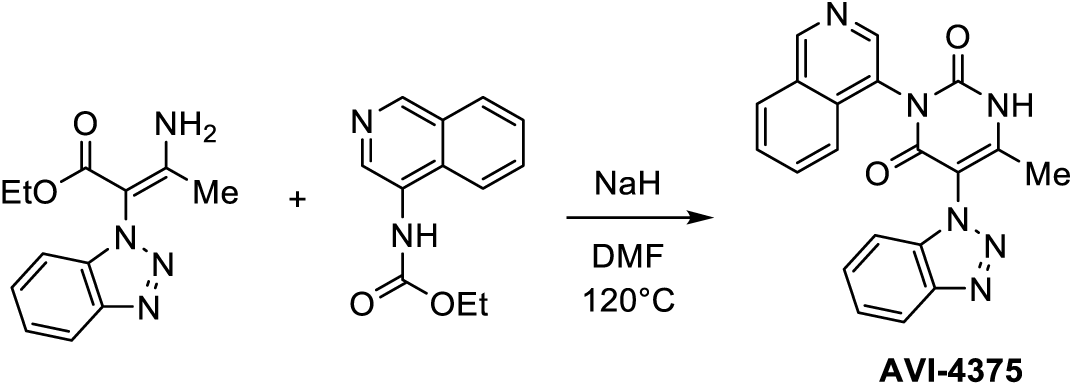

General Procedure C (Synthesis of Uracil) was followed using ethyl 3-amino-2-(1H-benzo[d][1,2,3]triazol-1-yl)-but-2-enoate (112 mg, 0.447 mmol). Purification by preparative HPLC (10% to 100% CH_3_CN in H_2_O + 0.1% formic acid) afforded **AVI-4375** (30.2 mg, 0.082 mmol, 18%) as a white solid.

**General**: C_20_H_14_N_6_O_2_; MW = 370.11.

**^1^H-NMR** (400 MHz, CD_3_OD): δ (ppm): 12.33 (brs, 1H); 9.42 (s, 1H); 8.61 (d, *J* = 12.0 Hz, 1H); 8.28 (d, *J* = 8.1 Hz, 1H); 8.13 (d, *J* = 8.6 Hz, 1H); 8.01 (q, *J* = 10.0 Hz, 1H); 7.90-7.76 (m, 2H); 7.62 (t, *J* = 7.6 Hz, 1H); 7.53-7.50 (q, 1H); 7.43 (t, *J* = 7.1 Hz, 1H); 7.38 (t, *J* = 7.7 Hz, 1H); 2.01 (s, 3H).

**LCMS (ESI)**: *m/z* = 371.30 [M+H]^+^.

### Step 3: AVI-4516

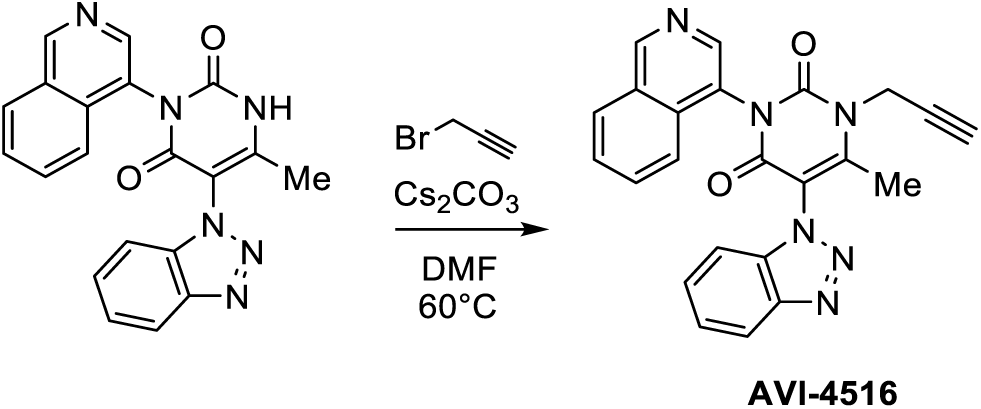

General Procedure D (Alkylation of Uracil) was followed using AVI-4375 (20 mg, 0.054 mmol). The mixture was stirred at 60°C for 4h. Yield: 9.2 mg, 0.023 mmol, 42%.

**General**: C_28_H_17_ClN_6_O_2_; MW = 408.42.

**^1^H-NMR** (400 MHz, DMSO-d6): δ (ppm): 9.43 (s, 1H); 8.67 (d, *J* = 17.4 Hz, 1H); 8.28 (d, *J* = 8.0 Hz, 1H); 8.17-8.13 (m, 2H); 7.92 (q, *J* = 8.5 Hz, 2H); 7.94-7.76 (m, 4H); 7.64 (dt, *J* = 1.9 Hz, 1H); 7.48 (dt, *J* = 2.7 Hz, 1H); 5.06 -4.87 (m, 2H); 3.60 (m, 1H); 2.28 (d, *J* = 7.4 Hz, 3H).

**LCMS (ESI)**: *m/z* = 409.3 [M+H]^+^.

#### Synthesis of AVI-4773

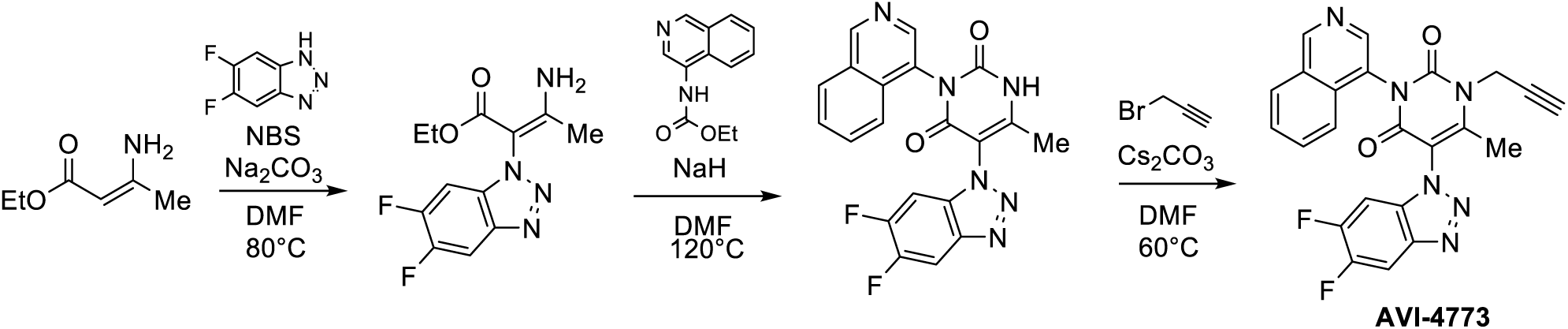

### Step 1: Ethyl 3-amino-2-(5,6-difluoro-1H-benzo[d][1,2,3]triazol-1-yl)but-2-enoate

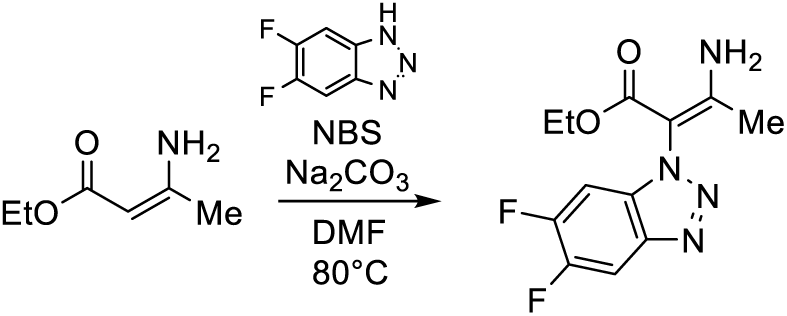

General Procedure B (Amination of Enaminone) was followed using ethyl 3-aminobut-2-enoate (83.3 mg, 0.645 mmol) and 5,6-difluoro-1*H*-benzotriazole. Yield: 137 mg, 0.485 mmol, 75%.

**General**: C_12_H_12_F_2_N_4_O_2_; MW = 282.09.

**LCMS (ESI)**: *m/z* = 283.06 [M+H]^+^.

### Step 2: AVI-4771

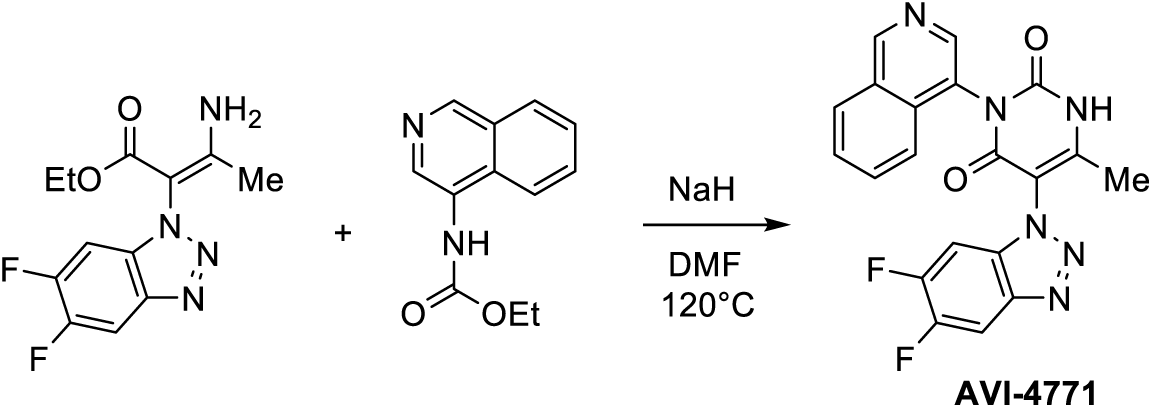

General Procedure C (Synthesis of Uracil) was followed using ethyl 3-amino-2-(5,6-difluoro-1H-benzo[d][1,2,3]triazol-1-yl)but-2-enoate (50 mg, 0.18 mmol). Purification by preparative HPLC (10% to 100% CH_3_CN in H_2_O + 0.1% formic acid) afforded **AVI-4771** (30 mg, 0.074 mmol, 42%) as a yellow solid.

**General**: C_20_H_12_F_2_N_6_O_2_; MW = 406.10.

**^1^H-NMR** (400 MHz, CD_3_CN): δ (ppm): 12.35 (s, 1H); 9.43 (s, 1H); 8.61 (s, 1H); 8.35-8.31 (dd, *J* = 2.1 Hz, 1H); 8.27 (d, *J* = 8.1 Hz, 1H); 8.17 (m, 2H); 7.89 (t, *J* = 7.6 Hz, 1H); 7.78 (t, *J* = 7.7 Hz, 1H); 7.24 (dd, *J* = 8.3, 2.1 Hz, 1H); 2.03 (s, 3H).

**LCMS (ESI)**: *m/z* = 407.07 [M+H]^+^.

### Step 3: AVI-4773

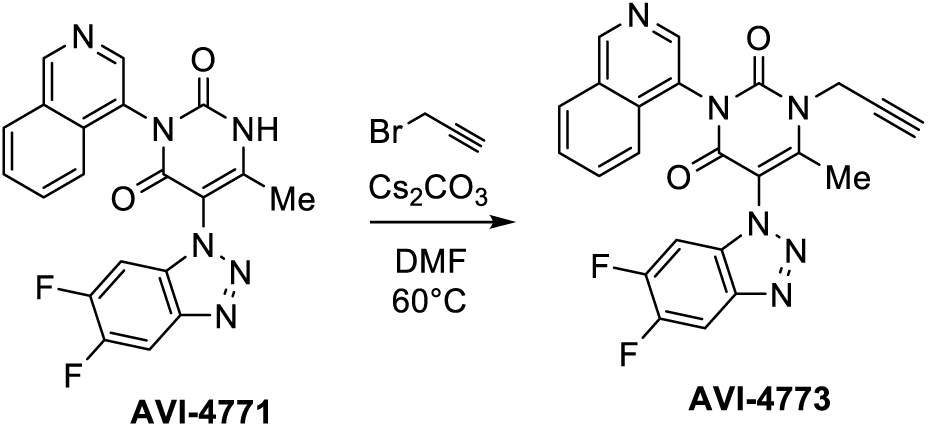

General Procedure D (Alkylation of Uracil) was followed using AVI-4771 (30 mg, 0.074 mmol). The mixture was stirred at 60°C for 4h. Yield: 8 mg, 0.02 mmol, 20%.

**General**: C_28_H_17_ClN_6_O_2_; MW = 444.11.

**^1^H-NMR** (400 MHz, DMSO-d6): δ (ppm): 9.41 (s, 1H); 8.67 (d, *J* = 18.9 Hz, 1H); 8.28 (d, *J* = 8.9 Hz, 1H); 8.14-8.02 (m, 2H); 7.89-7.82 (m, 2H); 7.76 (t, *J* = 7.6 Hz, 1H); 5.18-4.97 (m, 2H); 3.13 (m, 1H); 2.47 (d, *J* = 3.1 Hz, 3H).

**LCMS (ESI)**: *m/z* = 445.13 [M+H]^+^.

